# Transcriptome-scale spatial gene expression in the human dorsolateral prefrontal cortex

**DOI:** 10.1101/2020.02.28.969931

**Authors:** Kristen R. Maynard, Leonardo Collado-Torres, Lukas M. Weber, Cedric Uytingco, Brianna K. Barry, Stephen R. Williams, Joseph L. Catallini, Matthew N. Tran, Zachary Besich, Madhavi Tippani, Jennifer Chew, Yifeng Yin, Joel E. Kleinman, Thomas M. Hyde, Nikhil Rao, Stephanie C. Hicks, Keri Martinowich, Andrew E. Jaffe

## Abstract

We used the 10x Genomics Visium platform to define the spatial topography of gene expression in the six-layered human dorsolateral prefrontal cortex (DLPFC). We identified extensive layer-enriched expression signatures, and refined associations to previous laminar markers. We overlaid our laminar expression signatures onto large-scale single nuclei RNA sequencing data, enhancing spatial annotation of expression-driven clusters. By integrating neuropsychiatric disorder gene sets, we showed differential layer-enriched expression of genes associated with schizophrenia and autism spectrum disorder, highlighting the clinical relevance of spatially-defined expression. We then developed a data-driven framework to define unsupervised clusters in spatial transcriptomics data, which can be applied to other tissues or brain regions where morphological architecture is not as well-defined as cortical laminae. We lastly created a web application for the scientific community to explore these raw and summarized data to augment ongoing neuroscience and spatial transcriptomics research (http://research.libd.org/spatialLIBD).

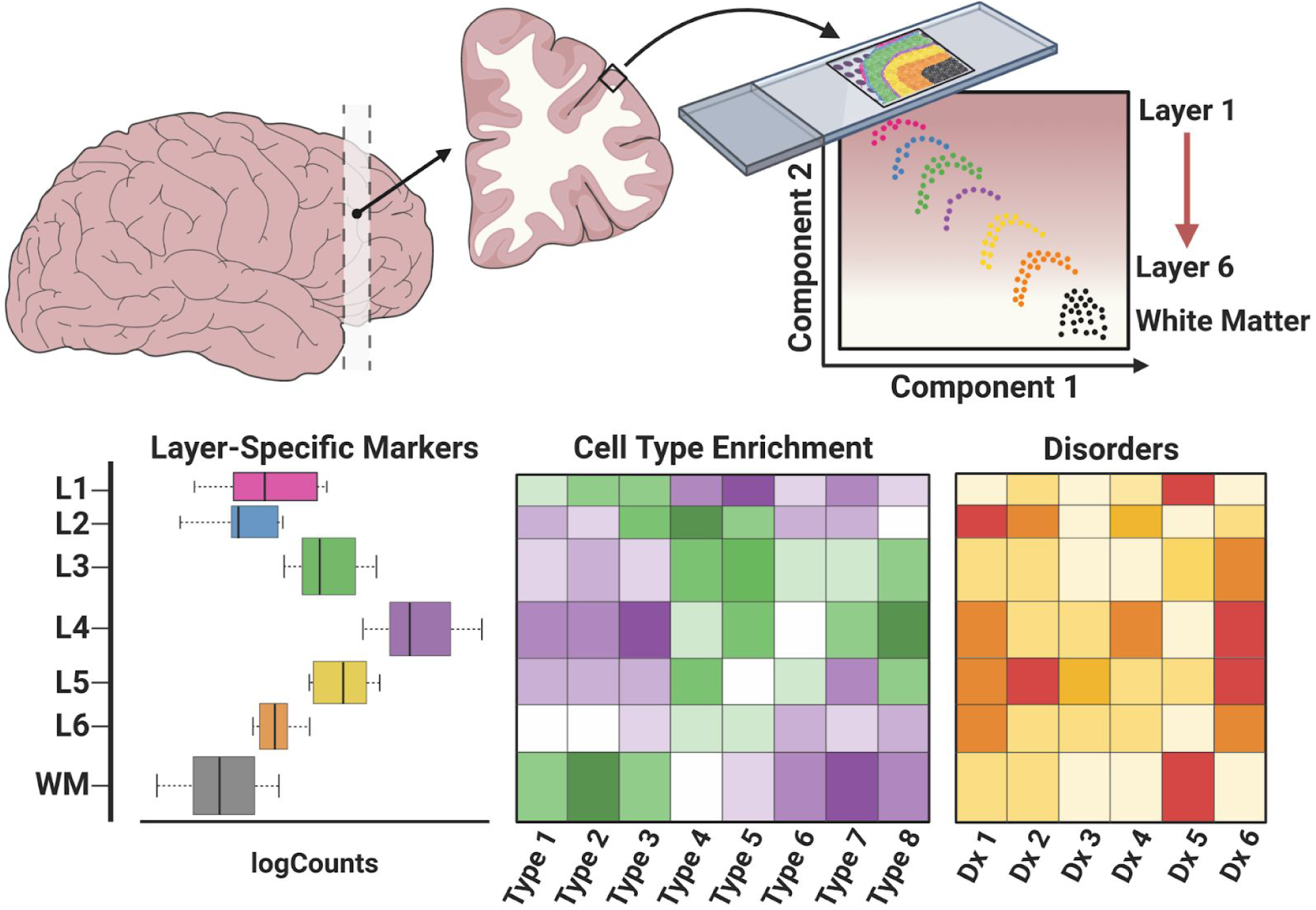

## Introduction

The spatial organization of the brain is fundamentally related to its function. This structure-function relationship is especially apparent in the context of the laminar organization of the human cerebral cortex where cells residing within different cortical layers show distinct gene expression patterns and exhibit differing patterns of morphology, physiology, and connectivity (DeFelipe and Fariñas, 1992; Harris and Shepherd, 2015; Narayanan et al., 2017; Radnikow and Feldmeyer, 2018). To the extent that structure entrains function, understanding normal brain development as well as disorders of the central nervous system will require identifying the cell types that make up the brain, and ultimately linking functional correlates of individual cell classes with structural architecture.

Major advances in single-cell (scRNA-seq) and single-nuclei (snRNA-seq) sequencing technologies have dramatically increased identification of molecularly-defined cell types in the human brain and implicated unique cell classes in risk for specific brain disorders (Darmanis et al., 2015; Hodge et al., 2019; Lake et al., 2016, 2018; Mathys et al., 2019; Nowakowski et al., 2017; Velmeshev et al., 2019). While scRNA-seq approaches are common in rodent brain tissue, the relatively large size and fragility of human neurons, coupled with the fact that most available postmortem human brain tissue is frozen, has resulted in nearly all available data in the human brain being generated on isolated nuclei with snRNA-seq approaches (Skene et al., 2018). While nuclear profiles are generally representative of whole cell profiles (Bakken et al., 2018), isolated nuclei lack the cytoplasmic compartment as well as axons and proximal dendrites, which limits our understanding of gene expression in the cytosol and neuropil (Skene et al., 2018). This is problematic for studies of brain disorders as converging evidence suggests that impairments in the formation or maintenance of synapses in critical cortical microcircuits are involved in many neuropsychiatric and neurodevelopmental disorders, including schizophrenia disorder (SCZD) and autism spectrum disorder (ASD) (Moyer et al., 2015; Sweet et al., 2010; Velmeshev et al., 2019). Indeed, studies in the postmortem brains of individuals with these disorders have implicated not only specific cell types (Gandal et al., 2018; Skene et al., 2018; Velmeshev et al., 2019), but also revealed differences in neuronal and synaptic structure that are spatially localized to specific cortical layers (Sweet et al., 2010; Velmeshev et al., 2019).

Furthermore, genes associated with increased risk for SCZD that were identified by genome-wide association studies (GWAS) are preferentially enriched for synaptic neuropil transcripts (Skene et al., 2018), suggesting that the extra-nuclear information missed by snRNA-seq approaches may be especially relevant for understanding genetic risk for brain disorders. While molecular profiles derived from sc/sn-RNAseq data can be used to predict anatomical location based on canonical marker genes described in the literature or from curated datasets, precisely assigning gene expression to the spatial coordinates of individual cell populations within intact brain cytoarchitecture of postmortem human brain tissue would significantly advance our understanding of studies of human brain development and disease.

Because it is considered a gold standard for quantifying gene expression with high spatial resolution, we recently established and optimized methods for using multiplex single-molecule fluorescent in situ hybridization (smFISH) in postmortem human brain tissue (Maynard et al., 2019). However, multiplexing with these technologies is limited, and even for methodologies that can accommodate hundreds to thousands of transcripts simultaneously, molecular crowding within cells leads to fluorescence overlap, which introduces significant microscopy-related issues and computational challenges (Burgess, 2019; Lein et al., 2017). The relatively large size of the human brain and lipofuscin-derived autofluorescence pose additional challenges for microscopy-based spatial transcriptomic methods in postmortem human tissue.

While methods such as laser capture microdissection (LCM)-seq do allow for transcriptome-wide profiling from cytosol in a spatially-defined area (Dong et al., 2018; He et al., 2017; Jaffe et al., 2019), the tissue is removed from the surrounding spatial context and processed separately, hindering the ability to analyze gradients of gene expression and examine spatial relationships within intact sections.

Emerging technologies for genome-wide spatial transcriptomics offer significant potential for providing detailed molecular maps that overcome limitations associated with sn/scRNA-seq and microscopy-based spatial transcriptomic methods. Importantly, these technologies use an on-slide cDNA synthesis approach that captures gene expression in the architecture of intact tissue, meaning that information from cytosol and neuronal processes is retained (Rodriques et al., 2019; Ståhl et al., 2016). To further our understanding of gene expression within the context of the spatial organization of the human cortex, we used the recently-released, 10x Genomics Visium platform, a novel barcoding-based transcriptome-wide spatial transcriptomics technology, to generate spatial maps of gene expression in the six-layered dorsolateral prefrontal cortex (DLPFC) of the adult human brain. The Visium platform expands the spatial resolution 5-fold beyond the first-generation ‘Spatial Transcriptomics’ approach (Ståhl et al., 2016) upon which it is based. While the original approach was successfully used to generate gene expression atlases and identify perturbations in transcriptional pathways for several normal and pathological human tissues, including the developing heart (Asp et al., 2018), invasive ductal cancer (Ståhl et al., 2016), pancreatic ductal adenocarcinoma (Moncada et al., 2018), prostate cancer (Berglund et al., 2018), postmortem spinal cord (Maniatis et al., 2019) and cerebellum (Gregory et al., 2020) of patients with amyotrophic lateral sclerosis (ALS), it lacked the necessary spatial resolution to resolve both individual cells and laminar structures in the human cortex.

Since some differences in pathology and gene expression associated with neuropsychiatric disorders are localized to specific cortical layers (Sweet et al., 2010; Velmeshev et al., 2019), the ability to localize spatial gene expression in the human brain at cellular resolution will be critical to gain further insight into disease mechanisms. Towards this end, we sought to define the laminar topography of gene expression in the human DLPFC, a brain area that has been implicated in a number of neuropsychiatric disorders. We overlaid data from recent large-scale snRNA-seq studies in the human brain with our spatial data to first confirm our layer-enriched expression signatures, and to then increase precision in manual annotation of gene expression-driven clusters to cortical laminae. To exemplify the potential of this type of data for clinical translation, we integrated our dataset with various neuropsychiatric disorder gene sets to demonstrate preferential layer-enriched expression of ASD risk genes and layer-enriched association of risk for several neuropsychiatric disorders. Finally, we compared the manually-annotated laminar clusters to entirely data-driven spatial clusters in the same human cortical tissue, using an approach that can also be applied to other human tissues and brain regions that do not have as clear morphological patterning as the cerebral cortex. We provide these data and analysis tools as a significant scientific resource for the neuroscience community to augment current molecular profiling and spatial transcriptomics efforts in the human brain.

## Results

We profiled spatial gene expression in human postmortem DLPFC tissue sections from two pairs of ‘spatial replicates’ from three independent neurotypical adult donors. Each pair consisted of two, directly adjacent 10µm serial tissue sections with the second pair located 300µm posterior from the first, resulting in a total of 12 samples run on the Visium platform (**Figure 1A**, Table S1, Method Details: Tissue processing and Visium data generation). We sequenced each sample to a median depth of 291.1M reads (IQR: 269.3M-327.7M), which corresponded to a mean 3,462 unique molecular indices (UMIs) and a mean 1,734 genes per spot. We note these rates are analogous to snRNA-seq and scRNA-seq data using the 10x Genomics Chromium platform, where a ‘cell’ barcode on the Chromium platform corresponds to a ‘spatial’ barcode on the Visium platform. However, unlike snRNA-seq data from postmortem human brain, which contains high numbers of intronic reads that map to immature transcripts, we found strong enrichment of mature mRNAs with high mean rates of exonic alignments (mean: 83.3%, IQR: 82.5-84.3%, Method Details: Visium raw data processing). Independent processing and cell segmentation of high-resolution histology images acquired before on-slide cDNA synthesis indicated an average of 3.3 cells per spot (IQR: 1-4), with a mean 15.0% (IQR: 12.8-17.9%) spots per sample containing a single cell body and 9.7% (IQR: 5.4-12.3%) ‘neuropil’ spots that lacked any cell bodies (Method Details: Histology image processing and segmentation). Tissue sections were acquired in the plane perpendicular to the pial surface that extended to the gray-white matter junction. The orientation of each sample was confirmed by delineating the border between layer 6 (L6) and the adjacent white matter (WM) and identifying layer (L5) using marker genes for gray matter/neurons (*SNAP25*), WM/oligodendrocytes (*MOBP*), and L5 (*PCP4*) in each tissue section (Figure S1, Figure S2, and Figure S3).

**Figure 1:**
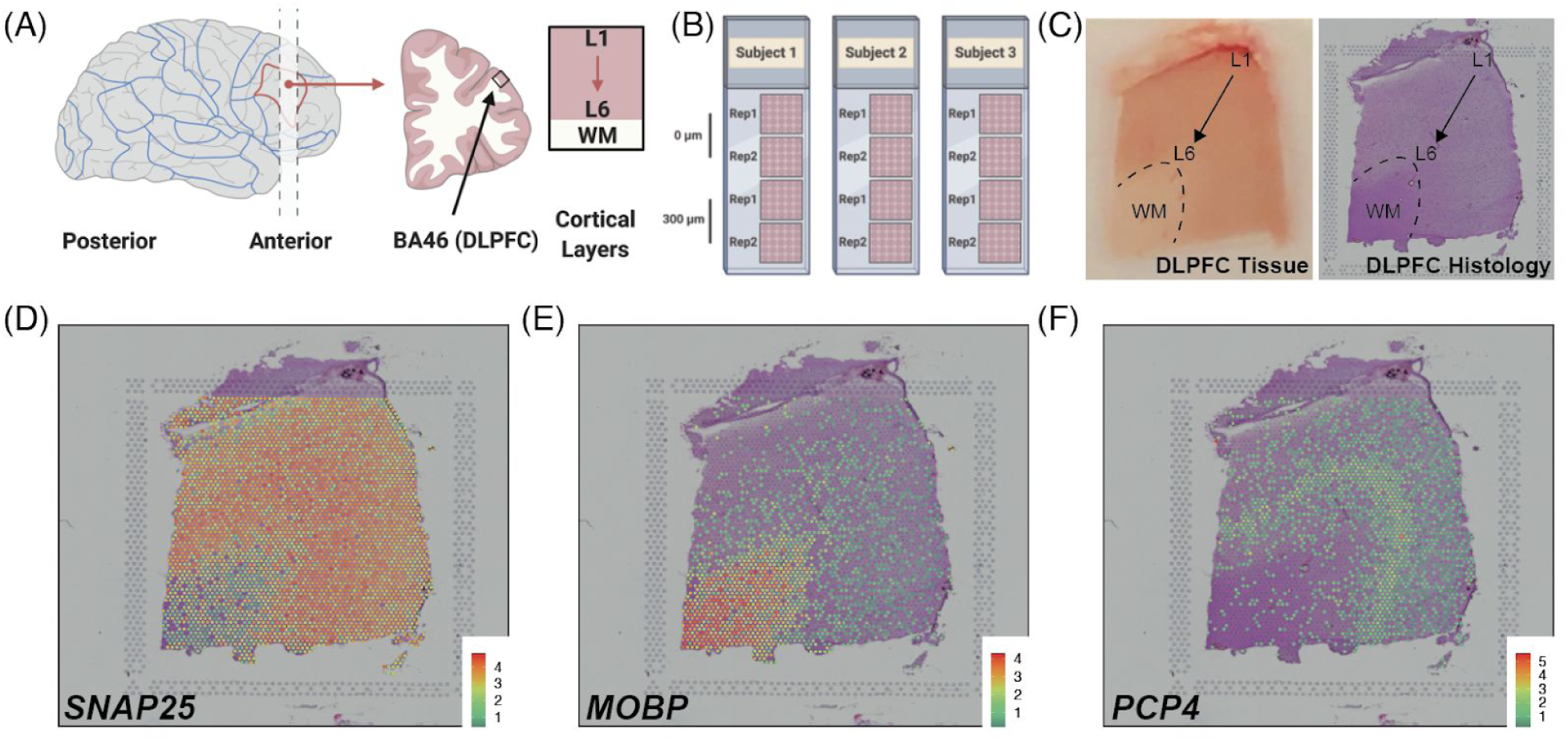
Spatial transcriptomics in DLPFC using Visium. (**A**) Tissue blocks of DLPFC were acquired in the anatomical plane perpendicular to the pial surface and extended to the gray-white matter junction. Each block spanned the 6 cortical layers and WM. (**B**) Schematic of experimental design including two pairs of ‘spatial replicates’ from three independent neurotypical adult donors. Each pair consisted of two, directly adjacent 10µm serial tissue sections with the second pair located 300µm posterior from the first, resulting in a total of 12 samples run on the Visium platform. (**C**) DLPFC tissue block and corresponding histology from sample 151673. (**D-F**) Spotplots depicting log-transformed normalized expression (logcounts) for sample 151673 for genes *SNAP25* (**D**), *MOBP* (**E**), and *PCP4* (**F**). Expression of these genes confirmed the spatial orientation of each sample by delineating the border between gray matter/neurons (*SNAP25*) and white matter/oligodendrocytes (*MOBP*) and defining L5 (*PCP4*). Spotplots of *SNAP25*, *MOBP*, and *PCP4* for all 12 samples can be found in Figure S1, Figure S2, and Figure S3. See also Table S1.

### Gene expression in the DLPFC across cortical laminae

We first generated aggregated layer-enriched expression profiles for each spatial replicate using a ‘supervised’ approach. We used cytotectonic architecture (Rajkowska and Goldman-Rakic, 1995a, 1995b) and robustly expressed region/layer-enriched markers (*MBP*-WM, *PCP4*-L5) combined with a dimensionality reduction method, specifically *t*-Distributed Stochastic Neighbor Embedding (*t*-SNE) (van der Maaten and Hinton, 2008), to assign individual spots to each of the six neocortical layers or the WM (Figure S4, Method Details: Spot-level data processing). Then, we performed ‘pseudo-bulking’ (Crowell et al., 2019; Kang et al., 2018; Lun and Marioni, 2017) by summing the UMI counts for each gene within each layer across each spatial replicate to generate layer-enriched expression profiles (**Figure 2A**, Method Details: Layer-level data processing). The pseudo-bulking approach, summarizing 47,681 spots to 76 layer-aggregated profiles across the 12 samples, removed sparsity and greatly increased UMI coverage of genes (**Figure 2A**). Unsupervised clustering of these layer-enriched expression profiles revealed the top component of variation in the data related to laminar differences, particularly between the white and gray matter (**Figure 2B**), with high concordance between the pairs of spatial replicates (Figure S5). Segmentation of histological images confirmed sparser cell densities in layer 1 (L1), a molecular layer enriched in synaptic processes, with 33.4% and 21.7% of spots containing 0 and 1 cell body, respectively. We observed increased cell densities in the oligodendrocyte-enriched WM, with 3.9% and 5.9% of spots containing 0 and 1 cell body, respectively (Table S2). We hypothesized that these ‘neuropil spots’ with 0 cell bodies may be enriched with neuronal processes (i.e. axons and dendrites; Table S3), and as predicted we identified significant enrichment of genes that are preferentially expressed in the transcriptome of synaptic terminals (Hafner et al., 2019) (⍴=0.38, *p*=1.9e-30, Figure S6) (Method Details: Neuropil enrichment analyses). Together, these analyses demonstrate the power of concurrently acquiring histology and gene expression data and highlight the ability of the Visium platform to achieve high resolution spatial expression profiling within the human DLPFC.

**Figure 2:**
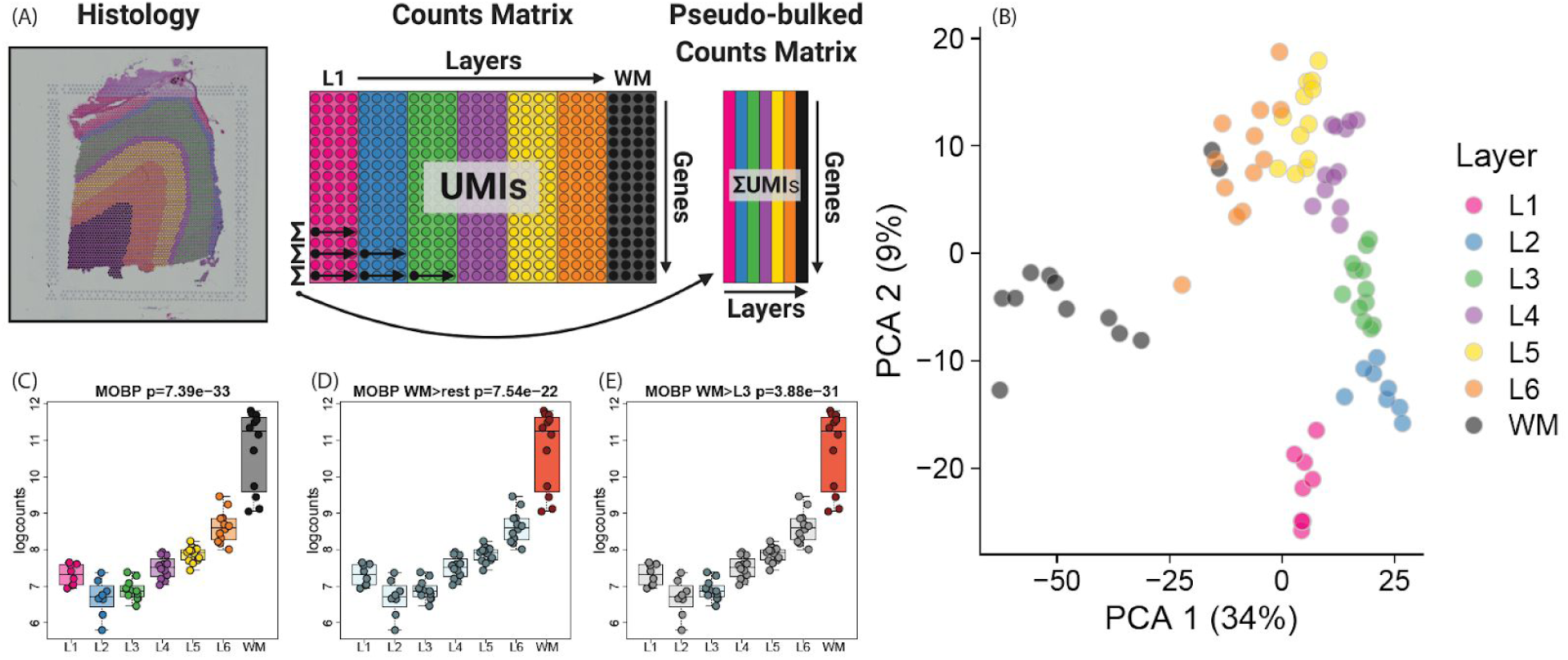
Layer-enriched gene expression in the DLPFC. (**A**) Visual description of the ‘pseudo-bulking’ statistical procedure, which collapses the spatial transcriptomics data from spot-level (∼4000 spots) to layer-level (6 layers + WM) data within each tissue section. (**B**) Principal component analysis (PCA) of layer-level (‘pseudo-bulked’) expression profiles across all sections and subjects. The first principal component separates the white and gray matter, and the second principal component associates with laminae. Visual depictions of the three statistical models employed to assess laminar enrichment, using *MOBP* as an example, including (**C**) “ANOVA” model, which tests whether the means of the seven layers are different, (**D**) ‘enrichment’ model, which tests whether each layer differs from all other layers - shown is WM (orange) vs other 6 layers (light blue), and (**E**) ‘pairwise’ model, which tests each layer versus each other layer - shown in WM (orange) versus L3 (light blue), which other layers in gray. See also Figure S4, Figure S5, Figure S7, and Table S4.

We used three strategies to perform differential expression (DE) analyses using the layer-enriched expression profiles generated above with linear mixed-effects modeling (Figure S7, Method Details: Layer-level gene modeling). The first strategy involved testing for differences in mean expression across the six layers plus WM (we also tested for differences in mean expression with only six layers, excluding WM), termed the ‘ANOVA’ model (**Figure 2C**), which estimates an F-statistic for each gene. This strategy revealed extensive differential expression across the laminar organization of the DLPFC, with 10,633 (47.6%) DE genes (DEGs) across the six gray matter layers plus WM (at FDR < 0.05) and 8,581 (38.4%) DEGs across the six gray matter layers excluding WM (FDR < 0.05). As expected, these results suggested extensive differences in gene expression between the layers of the DLPFC beyond broad white versus gray matter comparisons. The second strategy identified layer-enriched genes by testing for differences in expression between one layer versus all other layers, termed the ‘enrichment’ model (**Figure 2D**), which resulted in a *t*-statistic (termed ‘layer-enriched statistics’ hereafter) and *p*-value (and corresponding FDR adjusted *q*-value) for each expressed gene and layer (Method Details: Layer-level gene modeling). The largest expression differences were between WM and the neocortical layers, with 9,124 DEGs (FDR < 0.05), and the smallest differences were between L3 and all other layers with 183 DEGs genes (Table S4). In the third strategy, we tested for genes differentially expressed between each pair of layers (21 pairs), termed the ‘pairwise’ model (**Figure 2E**, Method Details: Layer-level gene modeling), which produced significant DEGs ranging from 8,500 for WM versus L3 to 292 for L4 versus L5 (Table S4). Together, these analyses highlight the extensive gene expression differences between the different layers of the human adult DLPFC.

### Identifying novel layer-enriched genes in human cortex

Several resources have compiled genes that exhibit laminar-specific expression across both rodent (Molyneaux et al., 2007) and human cortex (Zeng et al., 2012). While both overlapping and unique marker genes have been identified, these studies used different technologies, examined different developmental stages, and queried different regions of cortex. Therefore, we systematically assessed the robustness of these previously identified marker genes in our human adult DLPFC layer-enriched gene expression dataset. First, we tested for enrichment of previously published layer-enriched genes - as a set - among our layer-enriched DEGs, and found strong enrichment (*p*=1.22e-41). Since many of these marker genes were previously annotated to multiple layers (i.e. *CCK* and *ENC1*, **Figure 3**), rather than a single layer as queried in our DE analyses, we fit the ‘optimal’ statistical model for each gene using our layer-enriched expression profiles (Method Details: Known marker genes optimal modeling, Table S5). For example, *CCK* was annotated to L2, L3, L5 and L6, which were together tested against combining L1, L4, and WM in this optimal model. Only a subset of previously-associated layer-enriched genes showed high ranks and significant differential expression in our human DLPFC data (Figure S8), which were largely driven by markers identified by Zeng *et al*. (Zeng et al., 2012).

**Figure 3:**
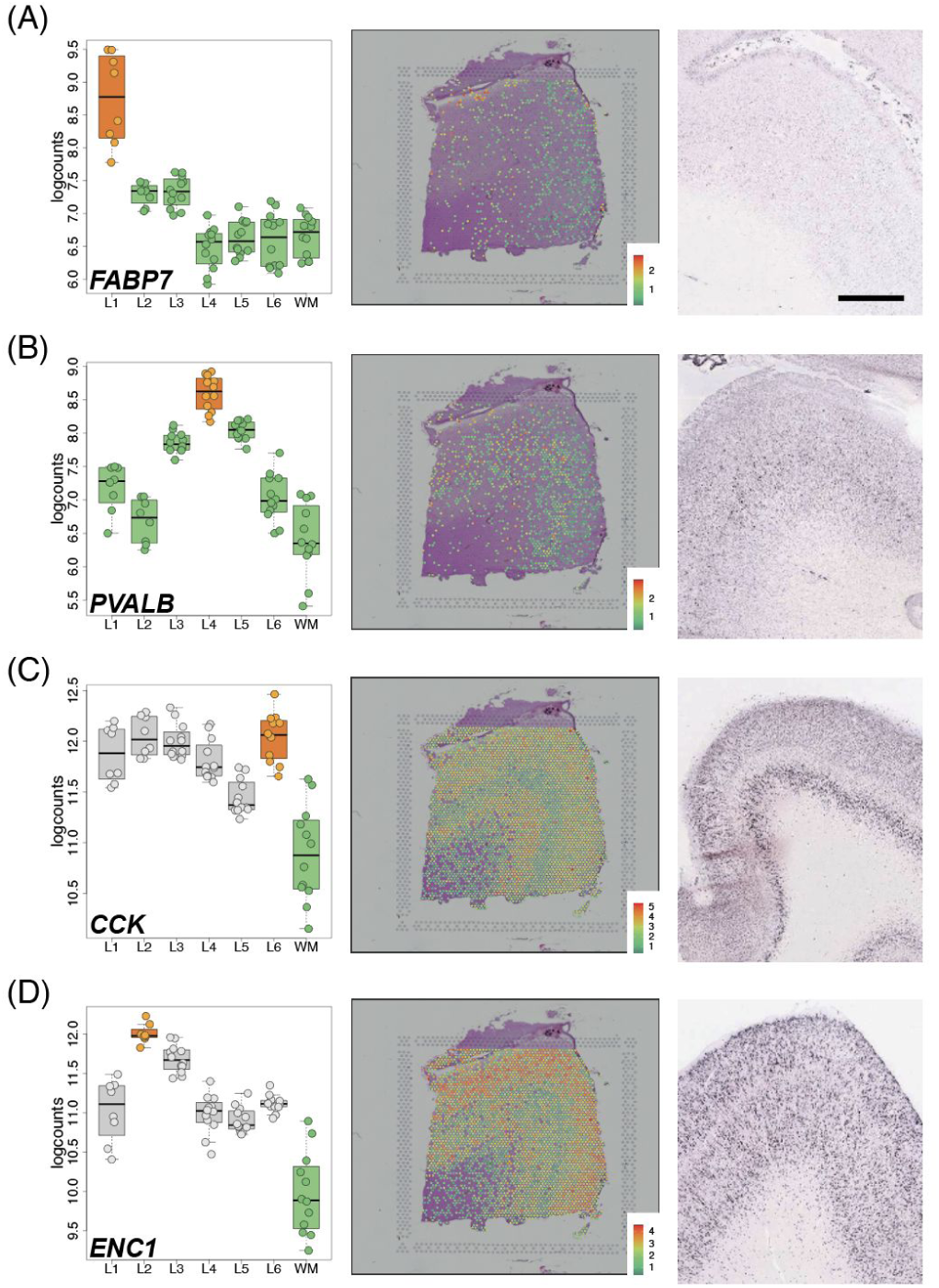
Visium replicates layer-enrichment of previously identified layer marker genes. (A-D) Left panels: Boxplots of log-transformed normalized expression (logcounts) for genes *FABP7* (**A,** L1>rest, *p*=5.01e-19), *PVALB* (**B,** L4>rest, *p*=1.74e-09), *CCK* (**C,** L6>WM, *p*=4.48e19), and *ENC1* (**D,** L2>WM, *p*=4.61e-25). Middle panels: Spotplots of log-transformed normalized expression (logcounts) for sample 151673 for genes *FABP7* (**A**), *PVALB* (**B**), *CCK* (**C**), and *ENC1* (**D**). Right panels: *in situ* hybridization (ISH) images from temporal cortex (**A, D**), DLPFC (**B**), or visual cortex (**C**) of adult human brain from Allen Human Brain Atlas: http://human.brain-map.org/ (Hawrylycz et al., 2012). Box and spot plots can be reproduced using our web application at: http://spatial.libd.org/spatialLIBD. Scale bar for Allen Brain Atlas ISH images=1.6mm. See also Figure S9 and Table S5.

We further confirmed laminar enrichment of a number of canonical marker genes, including *CCK*, *ENC1*, *CUX2*, *RORB*, and *NTNG2*, and validated these findings against publicly available singleplex *in situ* hybridization data from the Allen Brain Institute’s Human Brain Atlas (Hawrylycz et al., 2012) (**Figure 3** and Figure S9). Interestingly, while many of these genes (*FABP7*, *ADCYAP1*, *PVALB*) showed layer-enriched expression in our data, they were not classified by the Allen Brain Institute resources as being layer markers, demonstrating the utility of quantitative transcriptome-scale spatial approaches. Although we confirmed several canonical layer-enriched/specific genes, we found that only 59.5% of previously identified marker genes were significant DEGs (FDR < 0.05) in human DLPFC (Table S5). Indeed, we identified several genes previously underappreciated as laminar markers in human DLPFC, including *AQP4* (L1), *HPCAL1* (L2), *FREM3* (L3), *TRABD2A* (L5) and KRT17 (L6) (**Figure 4** and Figure S10). We validated these novel layer-enriched DEGs using multiplex single molecule fluorescent *in situ* hybridization (**Figure 4** and Figure S11, Methods Details: RNAscope smFISH). Novel layer-enriched DEGs were also validated by multiplexing with previously identified layer markers in the literature, many of which were also replicated in our Visium data (Figure S12).

**Figure 4:**
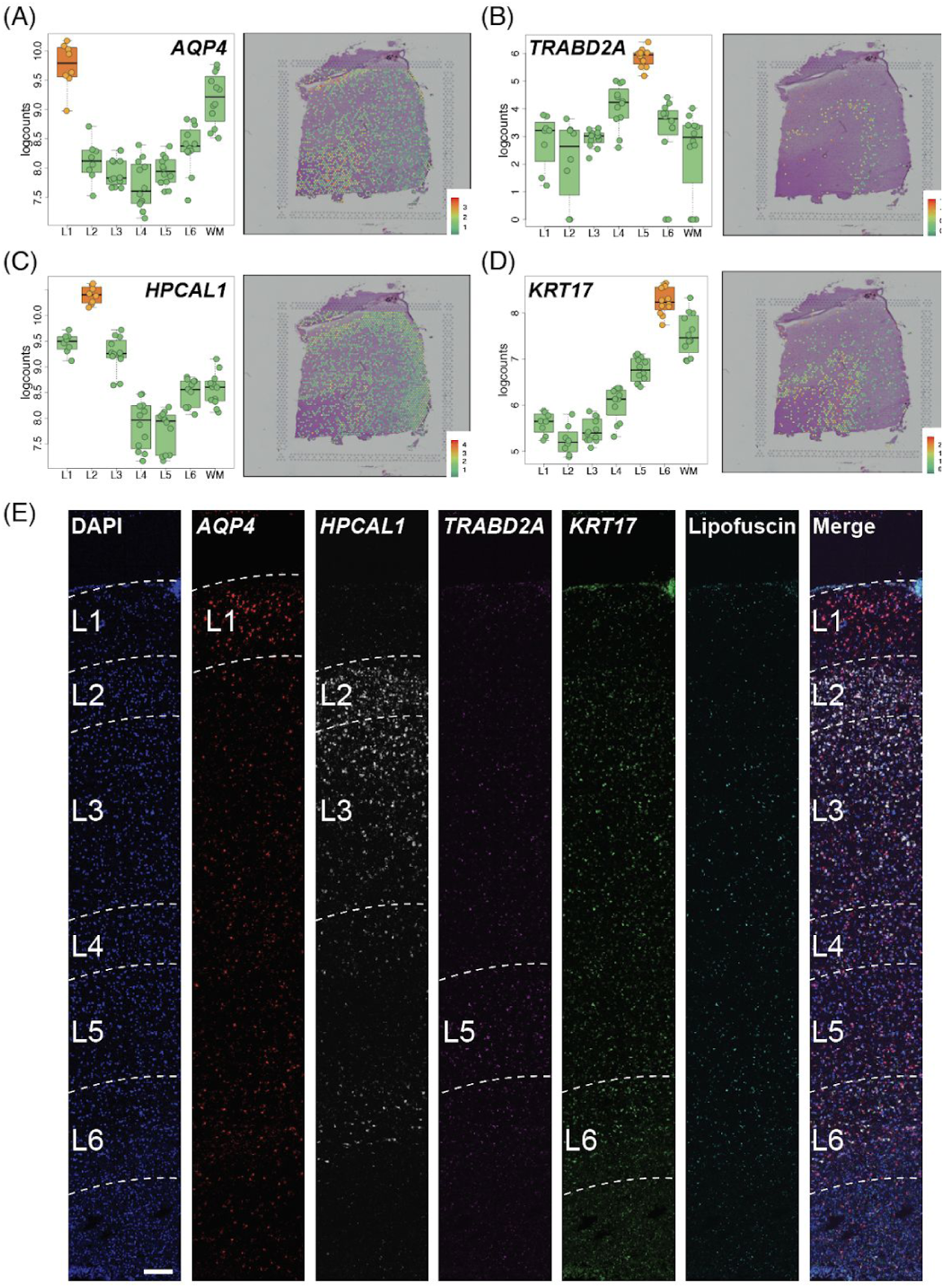
Discovery and smFISH validation of novel layer-enriched genes. (A-D) Left panels: Boxplots of log-transformed normalized expression (logcounts) for genes *AQP4* (**A,** L1>rest, *p*=1.47e-10), *TRABD2A* (**B,** L5>rest, *p*=4.33e-12), *HPCAL1* (**C**, L2>rest, *p*=9.73e-11), and *KRT17* (**D**, L6>rest, *p*=5.05e-12). Middle panels: Spotplots of log-transformed normalized expression (logcounts) for sample 151673 for genes *AQP4* (**A**), *TRABD2A* (**B**), *HPCAL1* (**C**) *and KRT17*(**D**). (**E**) Multiplex single molecule fluorescent in situ hybridization (smFISH) in a cortical strip of DLPFC. Maximum intensity confocal projections depicting expression of DAPI (nuclei), *AQP4*, *HPCAL1*, *TRABD2A, KRT17,* and lipofuscin autofluorescence. Merged image without lipofuscin autofluorescence. Scale bar=200μm. See also Figure S10, Figure S11, and Figure S12.

### Spatial registration of single nuclei RNA sequencing (snRNA-seq)

Adding spatial resolution to snRNA-seq datasets generated from human brain tissue has the potential to provide further insights about the function of molecularly-defined cell types. Specifically, layer-enriched expression profiles and differential expression statistics derived from the ‘enrichment model’ in our Visium data can be used to spatially “register” snRNA-seq datasets and add layer-enriched information to data-driven expression clusters that do not contain inherent anatomical information (**Figure 5A**, Methods Details: snRNA-seq spatial registration). We first used snRNA-seq data from Hodge *et al*. (Hodge et al., 2019) to confirm our layer-enriched expression profiles and validate this spatial registration strategy. While the snRNA-seq data in that study was obtained predominantly from NeuN+ sorted neuronal nuclei that were isolated from manually-dissected layers of the human postmortem middle temporal gyrus cortex, our layer-enriched DEGs from spatially-barcoded bulk tissue sections were in agreement with the laminar assignments from which these nuclei were derived (**Figure 5B**). We further validated this strategy on bulk RNA-seq data that was generated from manually-dissected laminar serial sections of the human cortex from four donors (He et al., 2017). This data however lacked corresponding histology data to definitively annotate specific cortical layers, and assignment of sections to layers likely underestimated the amount of WM present (∼5 sections/sample instead of just one predicted section), and missed L1 in one of their four subjects (H1) (Figure S13).

**Figure 5:**
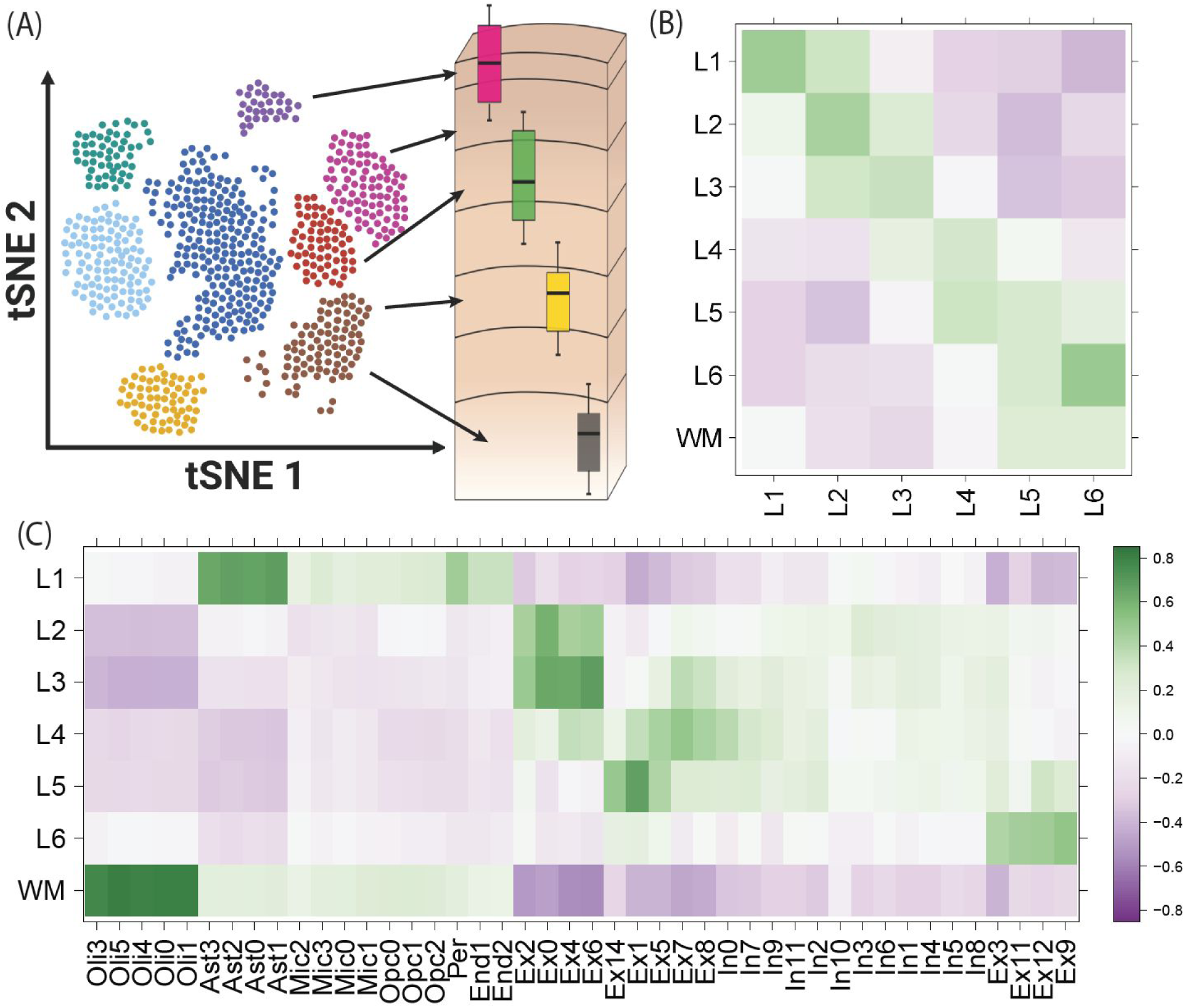
Spatial registration of snRNA-seq data. (**A**) Overview of the spatial registration approach. Heatmap of Pearson correlation values evaluating the relationship between our derived layer-enriched statistics (y-axis) for 700 genes and (**B**) layer-enriched statistics from snRNA-seq data in human medial temporal cortex produced by Hodge et al. (Hodge et al., 2019) (these data only profiled layers 1-6 in the gray matter, x-axis) and (**C**) cell type-specific statistics for cellular subtypes that were annotated by Mathys *et al*. from snRNA-seq data in human prefrontal cortex (Mathys et al., 2019)(x-axis). Oli = oligodendrocyte, Ast = astrocyte, Mic = microglia, Opc = oligodendrocyte precursor cell, Per = pericyte, End = endothelial, Ex = excitatory neurons, In = inhibitory neurons. See also Figure S13, Figure S14, and Figure S15.

We then used our layer-enriched statistics to perform spatial registration across three independent snRNA-seq datasets from human cortex. First, we generated our own snRNA-seq data from DLPFC using 5,231 nuclei from two donors, and performed data driven clustering to generate 30 preliminary cell clusters across 7 broad cell types (Figure S14, Method Details: DLPFC snRNA-seq data generation). Integration of our layer-enriched statistics refined excitatory and inhibitory neuronal subclasses into upper and deep layer subgroups beyond expected enrichments of glial cells in the WM (Figure S15 **A**). We further assessed the robustness of this approach by re-analyzing processed snRNA-seq from 48 donors across 70,634 nuclei obtained from the human prefrontal cortex (BA10) across 44 broad clusters in a study of Alzheimer’s disease (Mathys et al., 2019). Glial cell subpopulations showed expected enrichments, with preferential expression of oligodendrocyte subtypes in the WM, astrocyte subtypes in L1, and microglia, oligodendrocyte precursor (OPC), pericytes, and endothelial subtypes in both L1 and WM (**Figure 5C**). Neuronal cell subtypes showed greater laminar diversity, with multiple excitatory and inhibitory neuronal cell types associating with L2/L3, L4, L5, and L6 preferential expression, with generally more layer-enriched expression within excitatory cells (**Figure 5C**). Interestingly, our analysis showed that the excitatory neuronal subclasses (Ex2, Ex4, Ex6) identified by Mathys *et al*. that were most associated with clinical traits of Alzheimer’s disease were preferentially localized to the upper layers (L2/L3) of DLPFC in our data. This finding contrasts the inferences that were drawn by Mathys *et al*., which made layer assignments based on data obtained from the serial sections in He *et al*. described above (He et al., 2017). Specifically, they concluded that excitatory neuronal subclass Ex4 and Ex6 were preferentially expressed in the deeper layers while excitatory neuronal subclass Ex2 showed no laminar enrichment.

We lastly applied our spatial registration analysis to a study of autism spectrum disorder (ASD) (Velmeshev et al., 2019) including snRNA-seq data from 104,559 nuclei isolated from the human prefrontal cortex and anterior cingulate cortex that were obtained from 41 samples across 31 donors, which were annotated to 17 clusters in a study of ASD (Velmeshev et al., 2019) (Figure S15 **B**). As expected, we confirmed expected spatial contexts; for example, the highest enrichment of oligodendrocytes was again found in our histologically-defined WM. Our spatial registration framework was also able to refine the laminar predictions of cell-types in these previous studies. For example, integration of layer-enriched genes defined by Visium with snRNA-seq data from Velmeshev *et al*. indicated that astrocyte populations were most enriched in L1, while excitatory neurons annotated to L4 were more likely to be found in L5. These analyses demonstrate how this ‘spatial registration’ framework can be readily applied to any existing snRNA-seq or scRNA-seq datasets from dissociated cells to add back anatomical information.

### Clinical relevance of layer-enriched gene expression profiling

Given that several studies have identified associations between different brain disorders and molecularly-defined cell types, we assessed the clinical relevance of spatial gene expression using several different brain disorder-associated gene sets. We assessed the laminar enrichment of (1) gene sets derived from genes linked to different disorders via DNA profiling, (2) genes differentially expressed in postmortem brains of patients with a variety of brain disorders and neurotypical controls, and (3) genes associated with genetic risk via transcriptome-wide association studies (TWAS) (Gusev et al., 2016). We first used broad gene sets for different brain disorders compiled by Birnbaum *et al*. (Birnbaum et al., 2014), which showed laminar enrichments specifically for ASD (Figure S16, Table S6, Method Details: Clinical gene set enrichment analyses). We used the latest SFARI Gene database (Abrahams et al., 2013) to refine these associations, and demonstrate enrichments of L2 (OR=2.74, *p*=6.0e-21), L5 (OR=2.1, *p*=8.7e-7) and L6 (OR=2.7, *p*=1.8e-7) with ASD risk genes (**Figure 6A**). We confirmed the L2 (OR=3.6, *p*=3.9e-6) and L5 (OR=4.0, *p*=6.7e-5) associations in a recent exome sequencing study by Satterstrom *et al*. (Satterstrom et al., 2020), which identified 102 genes with ASD-associated variants. Interestingly, stratifying these genes by their clinical symptoms refined the laminar enrichments, as the 53 genes associated with ASD-dominant traits were more enriched for L5 (OR=4.9, *p*=5.3e-4, 8 genes: *TBR1, SATB1, ANK2, RORB, MKX, CELF4, PPP5C, AP2S1*), whereas the 49 genes associated with neurodevelopmental delay were more enriched for L2 (OR=4.5, *p*=7.8e-5, 12 genes: *CACNA1E, MYT1L, SCN2A, TBL1XR1, NR3C2, SYNGAP1, GRIN2B, IRF2BPL, GABRB3, RAI1, TCF4, ADNP*), suggesting that different functional subclasses of neurons might be contributing to each clinical subgroup. These layer-enriched expression associations for risk genes were largely independent of the enrichments seen comparing genes more highly expressed (WM: *p*=1.9e-29 and L1: *p*=4.5e-61) or more lowly expressed (L3: *p*=2.9e-5, L4: *p*=1.7e-42, L5: *p*=3.2e-36, and L6: *p*=1.9e-7) in brains of ASD patients compared to neurotypical controls (Table S6).

**Figure 6:**
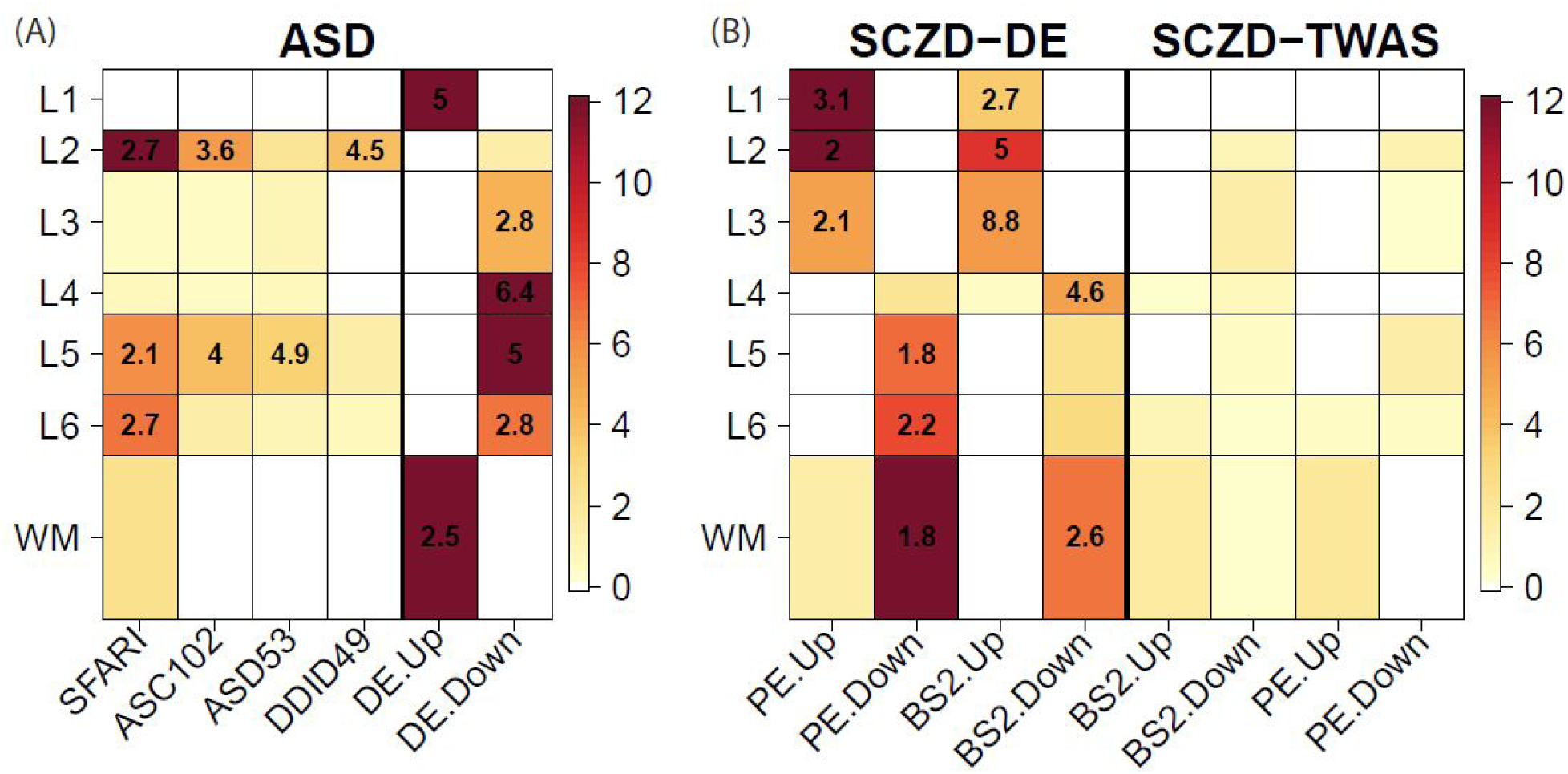
Layer-enrichment of neurodevelopmental and neuropsychiatric gene sets. We performed enrichment analyses using Fisher’s exact tests for our layer-enriched statistics versus a series of predefined gene sets related. (**A**) Autism spectrum disorder (ASD) laminar enrichments for SFARI (Abrahams et al., 2013) and Satterstrom *et al* (Satterstrom et al., 2020) for 102 overall ASD genes (ASC102), which were further stratified into 53 predominantly ASD (ASD53) and 49 predominantly developmental delay (DDID49) genes, as well as genes differentially expressed (DE) in the brains of individuals with ASD versus neurotypical controls as reported in the Gandal et al psychENCODE (PE) study (Gandal et al., 2018).(**B**) Schizophrenia disorder (SCZD) genes, including those from differential expression (DE) and transcriptome-wide association study (TWAS) analyses of RNA-seq data from brains of individuals with SCZD compared to neurotypical controls in the BrainSeq (BS) (Collado-Torres et al., 2019) and PE (Gandal et al., 2018) studies. ‘Up’ and ‘Down’ labels indicate whether genes are more highly or lowly expressed, respectively, in individuals with ASD or SCZD compared to neurotypical controls. Color scales indicate -log10(*p*-values), which were thresholded at *p*=10^-12^, and numbers within significant heatmap cells indicate odds ratios (ORs) for the enrichments. See also **Figure S16**, **Table S6**, **Table S7**, and **Table S8**.

We further assessed laminar enrichment of genes proximal to common genetic variation associated with SCZD, ASD, bipolar disorder (BPD), and major depressive disorder (MDD) (de Leeuw et al., 2015). These analyses identified significant overlap between L2-enriched and L5-enriched genes and risk for SCZD (at Bonferroni < 0.05), with additional overlap between L2-enriched genes and risk for bipolar disorder (at FDR < 0.05, Table S7). As above with ASD, there were markedly different laminar enrichments for genes associated with SCZD illness state.

Enrichment analyses of DEGs identified in two large SCZD postmortem brain datasets (Collado-Torres et al., 2019; Gandal et al., 2018), while highly convergent across studies, showed extensive enrichment across all layers, with increased expression of L1, L2, and L3 genes and decreased expression of WM, L4, L5 and L6 genes in patients compared to controls (**Figure 6B**). As secondary analyses, we performed heritability partitioning analysis (Finucane et al., 2015) for layer-enriched gene sets, which again identified significant heritability enrichment exclusively for L2 enriched-genes, specifically for SCZD, BPD, and educational attainment (Table S8, Method Details: Clinical gene set enrichment analyses). We additionally assessed TWAS statistics constructed for SCZD and BPD from single nucleotide polymorphism (SNP) weights computed from DLPFC (Gandal et al., 2018; Jaffe et al., 2020). While we did not observe strong enrichments of TWAS signal for any layer-enriched gene expression, SCZD risk genes in L2 and L5 suggested decreased expression in illness (**Figure 6B**, Table S6). Together, these analyses highlight the potential utility of these data in gleaning clinical insights by incorporating layer-enriched gene expression of the adult DLPFC into the interpretation of risk genes.

### Data-driven layer-enriched clustering in the DLPFC

Lastly, we explored the use of three alternative ‘data-driven’ approaches to classify Visium spots into laminar and non-laminar patterns, in contrast to the ‘supervised’ approach of identifying layer-enriched DEGs from manually-annotation of layers based on cytoarchitecture (**Figure 7A, B**; Figure S17), which may not be feasible in other brain regions or human tissues that lack clear or established morphological boundaries. Towards this goal, we explored the use of two gene sets: (1) genes exhibiting spatially variable expression patterns (SVGs) using the *SpatialDE* method (Svensson et al., 2018) within each of the 12 samples (Table S9), and (2) highly variable genes (HVGs) using the *scran* Bioconductor package (Lun et al., 2016). While no laminar information was used to identify SVGs and HVGs, interestingly these gene sets could identify both laminar and non-laminar spatial patterns (**Figure 7C, D**). For example, we identified several SVGs that were non-laminar, including *HBB, IGKC,* and *NPY*, which likely relate to blood cells, immune cells, and inhibitory interneuron classes (**Figure 7D**). In a completely data-driven and ‘unsupervised’ approach, we then used several implementations of unsupervised clustering methods with spot-level Visium data using these gene sets, with the possibility of further incorporating spatial coordinates of the spots, since we reasoned that adjacent spots should tend to show more similar expression levels (**Figure 7E**, Figure S18, Figure S19 and **Supplementary File 1**; Method Details: Data-driven layer-enriched clustering analysis). We compared these results to a ‘semi-supervised’ approach (unsupervised clustering guided by the layer-enriched genes identified using the DE “enrichment” models (Figure S7) and an approach using known rodent and human layer marker genes from Zeng *et al*. (Zeng et al., 2012) (**Figure 7E**, **Supplementary File 1**, and Table S10).

**Figure 7:**
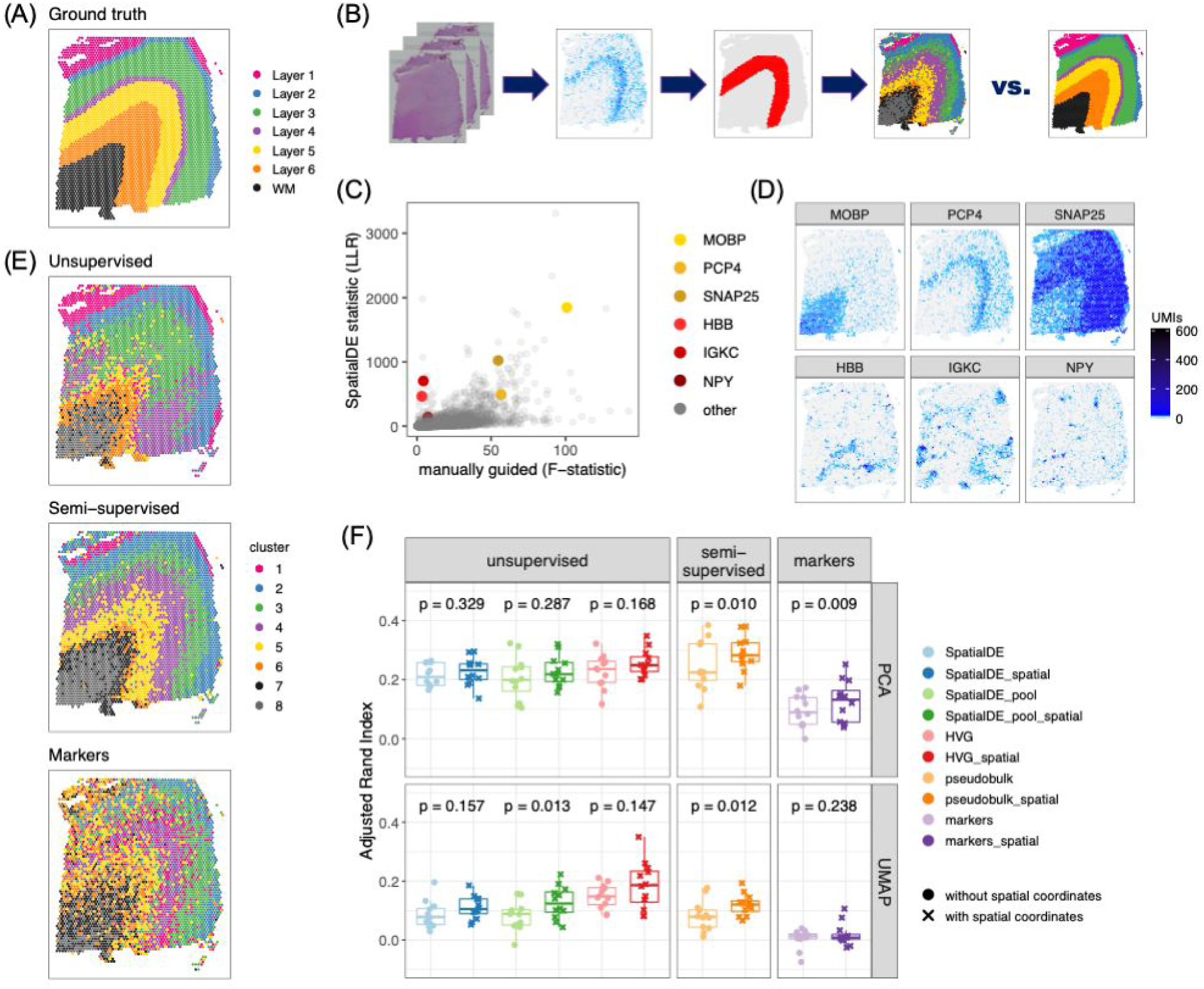
Data-driven layer-enriched clustering in the DLPFC. (**A**) Supervised annotation of DLPFC layers based on cytoarchitecture and selected gene markers (as shown in Figure 2A), used as ‘ground truth’ to evaluate the data-driven clustering results, for sample 151673. **(B)** Schematic illustrating the data-driven clustering pipeline, consisting of: (i) identifying genes (HVGs or SVGs) in an unbiased manner, (ii) clustering on these genes, and (iii) evaluation of clustering performance by comparing with ground truth. **(C)** Comparison of gene-wise test statistics for SVGs identified using *SpatialDE* (log-likelihood ratio, LLR) and genes from the DE ‘enrichment’ models (**Figure S7**) (F-statistics; WM included) for sample 151673. Colors indicate selected genes with laminar (red shades) and non-laminar (yellow shades) expression patterns. **(D)** Expression patterns for selected laminar (top row) and non-laminar (bottom row) genes identified using *SpatialDE* (corresponding to highlighted genes in (C)) in sample 151673. **(E)** Visualization of clustering results for the best-performing implementations of: (i) ‘unsupervised’ clustering (method ‘HVG_PCA_spatial’, which uses highly variable genes (HVGs) from *scran* (Lun et al., 2016), 50 principal components (PCs) for dimension reduction, and includes spatial coordinates as features for clustering); (ii) ‘semi-supervised’ clustering guided by layer-enriched genes identified using the DE enrichment models; and (iii) clustering guided by known markers from Zeng *et al*. (Zeng et al., 2012) (Method Details: Data-driven layer-enriched clustering analysis and **Table S10**). **(F)** Evaluation of clustering performance for all methods across all 12 samples, using manually annotated ground truth layers (as in (A)) and adjusted Rand index (ARI). Points represent each method and sample, with results stratified by clustering methodology (Method Details: Data-driven layer-enriched clustering analysis and **Table S10**). *P*-values represent statistical significance of the difference in ARI scores when including the two spatial coordinates as features within the clustering, using a linear model fit for each method (overall model across all methods: *p=*5.8e-6). See also **Figure S17**, **Figure S18**, **Figure S19**, **Table S9**, **Table S10**, and **Supplementary File 1**.

Using the manually-annotated layers as a ‘gold standard’ (**Figure 7A**, Figure S17), we evaluated the performance of the three approaches (‘unsupervised’, ‘semi-supervised’ and ‘markers’) using the adjusted Rand index (ARI) as the performance metric. Specifically, the ARI measures the similarity between the predicted cluster labels from our three approaches and the ‘gold standard’ cluster labels, with higher values corresponding to better performance (**Figure 7F**). First, we found consistent, but moderate, performance improvements by incorporating *x*, *y* spatial coordinates of the spots into the clustering methods across all three approaches (**Figure 7F**). Within the ‘unsupervised’ approach, we found that using the HVGs resulted in the highest ARI, but with the SVGs also comparable in performance (**Figure 7F**). However, the ‘semi-supervised’ approach resulted in the highest ARI out of all three approaches. This likely stems from the circularity of performing data-driven clustering guided by our layer-enriched DEGs on the same data, but this could be powerful in future spatial transcriptomics studies in the human cortex.

## Discussion

In this study we used the 10x Genomics Visium spatial transcriptomics platform to define the topography of gene expression in the DLPFC of the postmortem human brain. While a number of genome-scale spatial technologies have been successfully used in the mouse brain, our study is the first, to our knowledge, to implement Visium technology in human brain tissue. Based on examination of its histological organization and cytoarchitecture, the neocortex can be divided into six layers. Histological layers contain multiple cell types, including excitatory neurons, inhibitory neurons, and glia, and layers can be differentiated based on cell type composition and density, as well as morphology and connectivity of resident cell types (DeFelipe and Fariñas, 1992; Harris and Shepherd, 2015; Narayanan et al., 2017; Radnikow and Feldmeyer, 2018). Studies of postmortem brains from individuals with neuropsychiatric disorders have identified disease-associated changes in gene expression and synaptic structure that can be spatially localized to different cortical laminae (Sweet et al., 2010; Velmeshev et al., 2019). Because brain structure and function are tightly intertwined, defining the molecular landscape within the existing tissue architecture is a critical next step in understanding how brain function goes awry in neurodevelopmental, neuropsychiatric and neurodegenerative disorders. Our study takes a key step in adding new functional insights into spatially and molecularly-defined cell populations in the cortex by analyzing gene expression within the intact spatial organization of the human DLPFC.

First, we demonstrated the potential clinical translation of quantifying layer-enriched expression profiles in human brain samples. By integrating our data with clinical gene sets and genes differentially expressed in the brains of individuals with various neuropsychiatric disorders, we demonstrated preferential layer-enriched expression of genes implicated in ASD and SCZD. Genes that harbor *de novo* mutations associated with ASD (Satterstrom et al., 2020) were preferentially expressed in L2 and L5 based on Visium data. Subsets of these genes associated with specific clinical characteristics could be further partitioned into specific laminae, as genes predominantly associated with neurodevelopmental delay (NDD) were preferentially expressed in L2 and genes predominantly associated with ASD were preferentially expressed in L5. These specific laminar associations with penetrant *de novo* variants were in contrast to broad laminar enrichments of genes differentially expressed in the brains of patients with ASD (Gandal et al., 2018) and lack of laminar enrichment of genes implicated by common genetic variation (Grove et al., 2019). Interestingly these same two layers - L2 and L5 - showed preferential enrichment of genes implicated in common variation for SCZD (Pardiñas et al., 2018), and to a lesser extent, BPD (Stahl et al., 2019). These results were in contrast to differential expression analyses from postmortem studies of brain tissue from patients with SCZD compared to neurotypical controls (Collado-Torres et al., 2019; Gandal et al., 2018), which showed increased expression of upper layer genes and decreased expression of deep layer and WM genes. Further, we show that the heritability of schizophrenia is enriched for L2, a finding that implicates intracortical information processing as the focus of genetic risk mechanisms. These spatial gene expression patterns thus refine the laminar contexts of different neuropsychiatric disorders and may provide new targets for molecular interrogation.

Second, we overlaid recent large-scale snRNA-seq data from several cohorts to both confirm our layer-enriched expression signatures and further annotate gene expression-driven clusters to individual cortical layers. The shift from homogenate sequencing studies of brain tissue (Collado-Torres et al., 2019; Fromer et al., 2016; Jaffe et al., 2018) to large-scale snRNA-seq has already begun, with increasing sample sizes and numbers of nuclei (Mathys et al., 2019; Velmeshev et al., 2019), and will only continue to grow. Our strategy of “spatial registration” using individual gene-level statistics from both layer-specific versus cell type-specific expression profiles from hundreds or thousands of genes is likely more powerful than table-based enrichment analyses using small subsets of previously-defined marker genes.

Spatial registration of multiple independent datasets with our Visium data showed that layer-enriched patterns of expression can be extracted from snRNA-seq data, as subtypes of excitatory neuronal cells, and to a lesser extent, inhibitory neuronal cells, could be classified by their preferential laminar enrichment. While this strategy does not aid in constructing cell clusters in snRNA-seq data, it is a powerful tool to better annotate and interpret data-driven clusters and add spatial context to cell type-specific gene expression in the brain.

Third, in contrast to manually annotating laminar clusters based on cytoarchitecture, which is very labor-intensive, we evaluated the performance of alternative, data-driven approaches to cluster spots based on spatially variable genes (Svensson et al., 2018). We note that these unsupervised approaches can be used to identify novel spatial organizations, particularly those related to inhibitory neuronal subpopulations, brain vasculature, or immune function. Indeed, we identified variable spatial expression of 1) *NPY*, which encodes a neuropeptide highly expressed in a subpopulation of inhibitory interneurons, 2) *HBB,* which encodes a subunit of hemoglobin found in red blood cells, and 3) *IGKC*, which encodes the constant region of immunoglobulin light chains found in antibodies (**Figure 7D**). The layer-enriched genes defined here can be used to aid data-driven clustering in human cortex, and performed better than previously-defined markers (**Figure 7E, F**). Data-driven approaches identify previously unknown cellular organizations, and can also be applied to other human tissues or brain structures whose morphological patterning is not as defined as the cerebral cortex.

Microdissection techniques, including LCM approaches have been used to generate laminar-specific gene expression profiles in human cortex (Dong et al., 2018; He et al., 2017; Jaffe et al., 2019). However, because dissected regions are removed from the surrounding spatial context, boundaries cannot be definitively defined, hindering the ability to examine spatial relationships between cell populations or to define gradients of gene expression across structures. For example, several layer-enriched genes identified by Visium show striking gradients of gene expression, such as *HPCAL1* which is highly expressed in L2 but steadily decreases in expression through L4, L5, and L6. Conversely, *KRT17* is enriched in L6 and progressively decreases in expression through L5, L4, L3, and L2. Moreover, given the spatial organization of most brain regions, LCM approaches are often unable to isolate neuropil from cell bodies. In contrast, the Visium platform provides genome-wide transcriptomic information within the context of brain cytoarchitecture, which allowed us to sample regions containing only neuropil without having to perform specialized dissections. A major advantage of Visium in the human brain is the flexibility to analyze spatial gene expression from numerous angles (i.e. supervised clustering, unsupervised clustering, neuropil only) within a single experiment, which would be nearly impossible to accomplish with more labor intensive approaches such as LCM. While the current resolution of a spatially-barcoded spot in the Visium platform is 55μm, we found that 15.0% of spots contained a single cell body, highlighting an additional available level of interrogation for downstream analysis. Ongoing advances in these technologies will only improve this spatial resolution, as custom platforms can reach subcellular resolutions of 10μm and 2μm (Rodriques et al., 2019; Vickovic et al., 2019). Finally, Visium afforded several experimental advantages compared to fluorescence microscopy-based spatial transcriptomics approaches (Chen et al., 2015; Codeluppi et al., 2018) including, 1) coverage across a large area of brain tissue, 2) unbiased, transcriptome-wide analysis of gene expression (i.e. no requirement to select gene targets of interest), and 3) no confounds from lipofuscin autofluorescence. However, consistent dissections will be critical for applying Visium technology at large scale to generate equivalent clusters across tissue sections for spot aggregation approaches as performed here. As spatial transcriptomic technologies continue to develop, integration of transcriptomic and proteomic data in the same tissue section by incorporating immunohistochemical approaches will be an important future capability.

In contrast to the snRNA-seq approaches that encompass the vast majority of gene expression profiling studies in frozen postmortem human brain tissue, Visium is not limited to analysis of information in the nucleus. Indeed, on-slide cDNA synthesis methods preserve the integrity of information from both cytosol and neuronal processes, including dendrites and axons (neuropil). Cell segmentation of high-resolution histology images acquired before on-slide cDNA synthesis allowed us to determine that each spot contained an average of 3.3 cells with 9.7% of spots containing no cell bodies and only neuropil. We hypothesized that spots with no cell bodies would be enriched for transcripts highly expressed in neuronal processes and synapses. As predicted, we identified significant enrichment of genes preferentially expressed in synaptic terminals within ‘neuropil spots’ that contained no cell bodies. Given that robust evidence now supports the existence of localized mRNA expression and protein synthesis in both the pre- and post-synaptic compartments (Biever et al., 2019), directly studying neuropil-enriched transcripts in human brain has the potential to provide novel insights about expression of locally translated synaptic genes that may be missed with snRNA-seq analysis of dissociated nuclear preparations. Better understanding the regulation of synaptically localized transcripts in human cortex is important because the regulation of synaptic proteins controls neuronal homeostasis and drives synaptic plasticity (Biever et al., 2019). We further found enriched mitochondrial gene expression in sparser layers like L1 (Figure S20). This likely relates to our finding that L1 was most enriched for ‘neuropil spots’, and a higher energetic supply to axons and dendrites would be expected (Harris et al., 2012; Overly et al., 1996). Moreover, converging evidence suggests that impairments in the formation or maintenance of synapses in key circuits underlies risk for neuropsychiatric and neurodevelopmental disorders, including SCZD and ASD (Moyer et al., 2015; Sweet et al., 2010; Velmeshev et al., 2019). Supporting this notion, genes associated with increased risk for SCZD that were identified by GWAS were found to be preferentially enriched for synaptic neuropil transcripts (Skene et al., 2018).

While the laminar structure of the neocortex is largely preserved across mammalian species, several recent studies have underscored key similarities and differences in laminar gene expression between humans, primates, and rodents (He et al., 2017; Hodge et al., 2019; Zeng et al., 2012). Given the functional importance associated with laminar origin, recent snRNA-seq studies in postmortem human cortex have attempted to annotate molecularly-defined cell type clusters to the layer from which they originated (Mathys et al., 2019; Velmeshev et al., 2019) as discussed above. However, these laminar annotations are largely derived from curated gene sets that come from rodents and non-human primates, and not necessarily human studies. While we validated laminar-enrichment of some canonical layer-specific genes that were previously identified in the rodent and human cortex (**Figure 3** and Figure S9), some classical markers, such as *BCL11B* (L5), showed weak laminar patterning in DLPFC. Likewise, many genes showed no laminar patterning (Figure S8). These findings reinforce previous studies that urge caution in translating rodent and primate studies of molecularly and spatially-defined cell types into the human brain. Indeed, using a genome-wide approach such as Visium, we identified a number of previously underappreciated layer-enriched genes in human DLPFC, including *HPCAL1* (L2), *KRT17* (L6), and *TRABD2A* (L5), that may represent markers with higher fidelity for laminar annotation of snRNA-seq clusters in human brain (**Figure 4**). We also confirmed laminar enrichment of several genes identified as cell type markers in specific cortical layers by Hodges *et al*. (*LAMP5*, *AQP4, FREM3*).

In addition to these biological insights into the structure and function of the DLPFC, we have created several resources. All raw and processed data and code presented here are freely available to the scientific community through our web application (http://spatial.libd.org/spatialLIBD), to augment current neuroscience and spatial transcriptomics research. Through our application *“spatialLIBD”*, researchers can visualize the spot-level Visium data, manually annotate spots to layers, visualize the layer-level results, assess the enrichment of gene sets among layer-enriched genes, and perform spatial registration. These, and additional features, are described in detail at http://research.libd.org/spatialLIBD/.

In summary, our study demonstrates that the Visium spatial transcriptomics platform is capable of analyzing gene expression with high spatial resolution within the existing architecture of the human DLPFC. We demonstrate the ability to integrate Visium with snRNA-seq data for spatial registration, further increasing the utility in discovering patterns of gene expression within spatially defined cell populations in the normal as well as brain of individuals with neuropsychiatric disorders. Given the promise of spatial transcriptomics for linking molecular cell types with morphological, physiological and functional correlates of connectivity, we believe these approaches are the next frontier of transcriptomics in neuroscience and psychiatry. Our study represents a major advance towards this goal by providing data, resources and proof of concept examples for how this data can be used to understand human brain function and disease.

## Acknowledgements

The authors would like to express their gratitude to our colleagues whose efforts have led to the donation of postmortem tissue to advance these studies, including at the Office of the Chief Medical Examiner of the State of Maryland, Baltimore Maryland and the Office of the Chief Medical Examiner of Kalamazoo County Michigan. We also would like to acknowledge the contributions of Llewellyn B. Bigelow, MD and Amy Deep-Soboslay for their diagnostic expertise, and Daniel R. Weinberger for providing constructive commentary and editing of the manuscript. Finally, we are indebted to the generosity of the families of the decedents, who donated the brain tissue used in these studies. We would also like to thank the Accelerating Medicines Partnership - Alzheimer’s Disease (AMP-AD) Target Discovery and Preclinical Validation program and the ROSMAP study. We would like to thank William S. Ulrich for assistance with http://spatial.libd.org/spatialDE and the Department of Biostatistics at the Johns Hopkins Bloomberg School of Public Health for hosting mirrors of our web application. We thank the Johns Hopkins University Sidney Kimmel Comprehensive Cancer Center (SKCCC) Flow Cytometry Core and the Johns Hopkins University Transcriptomics and Deep Sequencing Core for supporting snRNA-seq experiments.

## Author contribution

- K.R.M. - Conceptualization, Methodology, Validation, Investigation, Writing, Visualization
- L.C-T. - Methodology, Software, Formal analysis, Data Curation, Writing, Visualization
- L.M.W. - Methodology, Software, Formal analysis, Writing, Visualization
- C.U. - Methodology, Investigation, Resources
- B.K.B. - Formal analysis, Data Curation, Visualization
- S.R.W. - Software, Data Curation
- J.L.C. - Software, Formal analysis, Visualization
- M.N.T. - Investigation, Formal analysis
- Z.B. - Software
- M.T. - Formal analysis, Visualization
- J.C. - Investigation
- Y.Y. - Investigation
- J.E.K. - Resources
- T.M.H. - Methodology, Resources
- N.R. - Resources, Supervision, Funding acquisition
- S.C.H. - Methodology, Software, Formal analysis, Writing, Visualization, Supervision
- K.M. - Conceptualization, Methodology, Writing, Supervision, Project administration, Funding acquisition
- A.E.J.- Conceptualization, Methodology, Software, Formal analysis, Writing, Visualization, Supervision, Project administration, Funding acquisition

## STAR Methods

### CONTACT FOR REAGENT AND RESOURCE SHARING

Further information and requests for resources and reagents should be directed to and will be fulfilled by the Lead Contact: Andrew E Jaffe.

### EXPERIMENTAL MODEL AND SUBJECT DETAILS

#### Post-mortem human tissue samples

Post-mortem human brain tissue from three donors of European ancestry (**Table S1**) was obtained by autopsy primarily from the Offices of the Chief Medical Examiner of the District of Columbia, and of the Commonwealth of Virginia, Northern District, all with informed consent from the legal next of kin (protocol 90-M-0142 approved by the NIMH/NIH Institutional Review Board). Clinical characterization, diagnoses, and macro- and microscopic neuropathological examinations were performed on all samples using a standardized paradigm, and subjects with evidence of macro- or microscopic neuropathology were excluded. Details of tissue acquisition, handling, processing, dissection, clinical characterization, diagnoses, neuropathological examinations, RNA extraction and quality control measures have been described previously (Lipska et al., 2006). Briefly, dorsolateral prefrontal cortex (DLPFC) was microdissected and embedded in OCT in a 10mm x 10mm cryomold. Each sample was dissected in a plane perpendicular to the pial surface in area 46 of the cortex to capture from the pial surface to the gray-white matter junction and spanned L1-6 and the WM.

### METHOD DETAILS

#### Tissue processing and Visium data generation

Frozen samples were embedded in OCT (TissueTek Sakura) and cryosectioned at −10C (Thermo Cryostar). Sections were placed on chilled Visium Tissue Optimization Slides (3000394, 10X Genomics) and Visium Spatial Gene Expression Slides (2000233, 10X Genomics), and adhered by warming the back of the slide. Tissue sections were then fixed in chilled methanol and stained according to the Visium Spatial Gene Expression User Guide (CG000239 Rev A, 10X Genomics) or Visium Spatial Tissue Optimization User Guide (CG000238 Rev A, 10X Genomics). For gene expression samples, tissue was permeabilized for 18 minutes, which was selected as the optimal time based on tissue optimization time course experiments. Brightfield histology images were taken using a 10X objective (Plan APO) on a Nikon Eclipse Ti2-E (27755 x 50783 pixels for TO, 13332 x 13332 pixels for GEX). Raw images were stitched together using NIS-Elements AR 5.11.00 (Nikon) and exported as .tiff files with low and high resolution settings. For tissue optimization experiments, fluorescent images were taken with a TRITC filter (ex/em brand) using a 10X objective and 400 ms exposure time.

Libraries were prepared according to the Visium Spatial Gene Expression User Guide (CG000239, https://assets.ctfassets.net/an68im79xiti/3pyXucRaiKWcscXy3cmRHL/a1ba41c77cbf60366202805ead8f64d7/CG000239_VisiumSpatialGeneExpression_UserGuide_Rev_A.pdf). Libraries were loaded at 300 pM and sequenced on a NovaSeq 6000 System (Illumina) using a NovaSeq S4 Reagent Kit (200 cycles, 20027466, Illumina), at a sequencing depth of approximately 250-400M read-pairs per sample. Sequencing was performed using the following read protocol: read 1, 28 cycles; i7 index read, 10 cycles; i5 index read, 10 cycles; read 2, 91 cycles.

#### Visium raw data processing

Raw FASTQ files and histology images were processed by sample with the Space Ranger software, which uses STAR v.2.5.1b (Dobin et al., 2013) for genome alignment, against the Cell Ranger hg38 reference genome “refdata-cellranger-GRCh38-3.0.0”, available at: http://cf.10xgenomics.com/supp/cell-exp/refdata-cellranger-GRCh38-3.0.0.tar.gz. Quality control metrics returned by this software are available in **Table S1**.

#### Histology image processing and segmentation

Histology images were processed and nuclei were segmented using the “Color-Based Segmentation using K-Means Clustering” in MATLAB. The MATLAB function `rgb2lab` is used to convert the image from RGB color space to CIELAB color space also called L*a*b color space (L - Luminosity layer measures lightness from black to white, a - chromaticity-layer measures color along red-green axis, b - chromaticity-layer measures color along blue-yellow axis). The CIELAB color space quantifies the visual differences caused by the different colors in the image. The a*b color space is extracted from the L*a*b converted image and is given to the K-means clustering function `imsegkmeans` along with the number of colors the user visually identifies in the image. The `imsegkmeans` outputs a binary mask for each color it identifies. Since the nuclei in the histology images have a bright color that can be easily differentiated from the background, a binary mask generated for the nuclei color is used as the nuclei segmentation.

The segmented binary mask was used to estimate the number of nuclei in each spot. For each histology image, there is a JSON file describing some properties of the image, including the spot diameter in pixels at the full-resolution image. Additionally, for each image there is a text file in tabular format that includes one row for each spot with an identification barcode, a row, a column and pixel coordinates for the center of the spot on the full-resolution image. Using this information, the following protocol was implemented. For each spot, all pixels of the binary mask were set to zero except those within the spot radius of the center of that spot. The resulting binary mask was then labeled with a unique integer for each unique contiguous cluster of pixels. The maximum of this labeled mask was stored as an estimate of the number of nuclei within that spot.

#### Spot-level data processing

The raw Visium files for each sample (Method Details: Visium raw data processing) were read into R into a custom structure using the *SummarizedExperiment* R package (Martin Morgan, 2017) to keep them paired with the low resolution histology images for visualization purposes. They were then combined into a single *SingleCellExperiment* (Aaron Lun [Aut, 2017) object (`sce`) to allow us to perform analyses using the gene expression data from all samples.

We added information, including the number of estimated cell counts (Method Details: Histology image processing and segmentation), the sum of UMIs per spot, number of expressed genes per spot, and graph-based clustering results (computed by sample) provided by 10x Genomics Space Ranger software to the `sce` object. We evaluated the per-spot quality metrics using the function `perCellQCMetrics` from the *scran* R Bioconductor package (Lun et al., 2016) and did not drop any spots given the spatial pattern they presented. We used the *scran (Lun et al., 2016)* functions `quickCluster,` blocking by the six pairs of spatially adjacent replicates, `computeSumFactors`, and *scater*’s (McCarthy et al., 2017) `logNormCounts` to compute the log-normalized gene expression counts at the spot-level. By modeling the gene mean expression and variance with the `modelGeneVar` *scran* (Lun et al., 2016) function, blocking again by the six pairs of spatially adjacent replicates, followed by `getTopHVGs` we identified the top 10% highly variable genes (HVGs): 1,942 genes. Using this subset of HVGs, we computed principal components (PCs) with *scater*’s (McCarthy et al., 2017) runPCA to produce 50 components. Using these 50 top PCs, we computed tSNE (van der Maaten and Hinton, 2008) and UMAP (McInnes et al., 2018) dimension reduction methods using `runTSNE` (perplexity 5, 20, 50, 80) and `runUMAP` (15 neighbors) from *scater* (McCarthy et al., 2017). With the top 50 PCs, we performed graph-based clustering across all samples using 50 nearest neighbors using buildSNNGraph from *scran* (Lun et al., 2016) and the Walktrap method from implemented by *igraph* (Csardi and Nepusz, 2006) resulting in 28 clusters (`snn_k50_k4` through `snn_k50_k28`). We further cut the graph to produce clusters from 4 to the 28 in increments of 1. Members of our team used an initial *spatialLIBD* (Collado-Torres, 2020) version to assign the graph-based clusters from 10x Genomic to the closest anatomical layers for each sample (`Maynard` and `Martinowich`).

All this information was combined and displayed through a *shiny (Chang et al., 2019)* web application at http://spatial.libd.org/spatialLIBD in such a way that we, and now the scientific community, can visualize the expression of a given gene, or a given set of clustering results, across all samples or each sample individually. For any chosen sample, *spatialLIBD* allows users to view gene expression and selected cluster results both in the context of spatial histology and given dimension reduction results (PCA, tSNE, UMAP) using *plotly* (Sievert, 2018). Using this web application to visualize cytoarchitecture as well as the expression patterns of MBP and *PCP4*, known WM and L5 marker genes, a single experimenter manually assigned each spot to a cortical layer for each sample for all but 352 out of the 47,681 spots across all samples. These 352 spots were located on small fragments of damaged tissue disconnected from the main tissue section. We added these supervised layer annotations to our `sce` object and the final version is available for download through the `fetch_data` function in *spatialLIBD* (Collado-Torres, 2020).

#### Layer-level data processing

For the subject with brain ID Br5595, which lacked L1 and clear cytoarchitecture for L2 and L3, we re-labelled all “L2/L3” ambiguous spots as L3 and dropped the 352 un-assigned spots. We then pseudo-bulked (Crowell et al., 2019; Kang et al., 2018; Lun and Marioni, 2017) the spots into layer-level data by summing the raw gene expression counts across all spots in a given sample and in a given layer, and repeated this procedure for each gene, sample and layer combination (Figure 2A). This resulted in 47,329 genes quantified across 76 layer-sample combinations (7 * 12 = 84, because not all layers were clearly observed in each sample as Br5595 had no distinct L1 or L2 across all four spatial replicates, **Figure S4**). This resulted in another *SingleCellExperiment* (Aaron Lun [Aut, 2017) object called `sce_layer`. We used `librarySizeFactors` and `logNormCounts` from *scater* (McCarthy et al., 2017) to compute layer-level log normalized gene expression values. We dropped all mitochondrial genes and retained genes that were expressed in at least 5% (4 / 76 layer-sample combinations) and had an average counts greater than 0.5 as computed by `calculateAverage` from *scater* (McCarthy et al., 2017), resulting in a final set of 22,331 genes. We identified 1,280 top HVGs at the layer-level and computed 20 PCs (Figure 2B) which we then used in the tSNE (perplexity = 5, 15 and 20) and UMAP (15 neighbors) computations similar to Method Details: Spot-level data processing. We then clustered the layer-level data using several graph-based approaches as well as using k-means. This is the `sce_layer` data that is available for download through the `fetch_data` function in *spatialLIBD* (Collado-Torres, 2020).

#### Neuropil enrichment analyses

We performed differential expression analysis at the spot-level in our Visium data, comparing the 4,855 spots with 0 cell bodies to the other 42,474 spots containing at least 1 cell body, adjusting for fixed effects of layer and spatial replicate. We downloaded differential expression statistics from RNA-seq of vGLUT1+ enriched synaptosomes in mouse brain from Hafner *et al*. (Hafner et al., 2019), and lined up these data at the gene-level using homologous entrez IDs between mouse and human (via http://www.informatics.jax.org/downloads/reports/HMD_HumanPhenotype.rpt). We compared the effects of spots containing 0 cells in our data to vGLUT1+ enriched cells from Hafner et al, both across the full homologous transcript and then within genes significant in the Hafner dataset at FDR < 0.05.

#### Layer-level gene modeling

Using the layer-level data we fit three types of models (Figure 2C, **Figure S7**):

1. *ANOVA*: For this model we tested for each gene whether the log normalized gene expression counts are variable between the layers by computing F-statistics. We used `lmFit` and `eBayes` from *limma* (Ritchie et al., 2015) after blocking by the six pairs of spatially adjacent replicates and taking this correlation into account as computed by `duplicateCorrelation`.
2. *Enrichment*: Using the same functions and taking into account the same correlation structure, we computed t-statistics comparing one layer against the other six using the layer-level data. This resulted in seven sets of t-statistics (one per layer) with double-sided *P*-values. We focused on genes with positive t-statistics (expressed higher in one layer against the others) as these are enriched genes instead of depleted genes.
3. *Pairwise*: Using the same functions and taking into account the same correlation structure in addition to using `contrasts.fit` from *limma* (Ritchie et al., 2015), we computed t-statistics for each pair of layers resulting in t-statistics with double-sided *P*-values.

The modeling results are available for download through the `fetch_data` function in *spatialLIBD* (Collado-Torres, 2020) as the `modeling_results` object as well as in **Table S4**.

### Known marker genes optimal modeling

Using two lists of known layer marker genes derived from previous mouse or human studies (Zeng et al., 2012) (Molyneaux et al., 2007), we identified 29 different unique optimal models for these genes. For example, L1 + L2 versus the other layers. Using the same modeling framework (Method Details: Layer-level gene modeling) we computed t-statistics for all genes at the layer-level data for each of these 29 unique models. For each of the 29 unique models, we then retained information about the statistics for the known marker genes matching the model as well as the top ranked (with a positive t-statistic gene, **Table S5**).

### RNAscope single molecule fluorescent in situ hybridization (smFISH)

Fresh frozen DLPFC from the same neurotypical control samples used for Visium were sectioned at 10μm and stored at −80°C. *In situ* hybridization assays were performed with RNAscope technology utilizing the RNAscope Fluorescent Multiplex Kit V2 and 4-plex Ancillary Kit (Cat # 323100, 323120 ACD, Hayward, California) according to the manufacturer’s instructions. Briefly, tissue sections were fixed with a 10% neutral buffered formalin solution (Cat# HT501128 Sigma-Aldrich, St. Louis, Missouri) for 30 minutes at room temperature (RT), series dehydrated in ethanol, pretreated with hydrogen peroxide for 10 minutes at RT, and treated with protease IV for 30 minutes. Sections were incubated with 4 different probe combinations: A) L1 and L5: *AQP4*, *RELN*, *TRABD2A*, *BCL11B* (Cat 482441-C4, 413051-C2, 532881, 425561-C3, ACD, Hayward, California); B) L3 and L6: *CARTPT*, *FREM3*, *NR4A2*, (506591, 829021-C4, 582621-C3); C) L2/3 and WM: *LAMP5*, *HPCAL1*, *NDRG1*, *MBP* (487691-C2, 846051-C3, 481471, 411051-C4); D) Visium-identified genes: *AQP4*, *TRABD2A*, *KRT17 (*463661-C2), *HPCAL1*. Following probe labeling, sections were stored overnight in 4x SSC (saline-sodium citrate) buffer. After amplification steps (AMP1-3), probes were fluorescently labeled with Opal Dyes (Perkin Elmer, Waltham, MA; 1:500) and stained with DAPI (4′,6-diamidino-2-phenylindole) to label the nucleus. Lambda stacks were acquired in z-series using a Zeiss LSM780 confocal microscope equipped with 20x x 1.4 NA and 63x x 1.4NA objectives, a GaAsP spectral detector, and 405, 488, 555, and 647 lasers. All lambda stacks were acquired with the same imaging settings and laser power intensities. For each subject, a cortical strip was tile imaged at 20x to capture L1 to WM. Following image acquisition, lambda stacks in *z*-series were linearly unmixed in Zen software (weighted; no autoscale) using reference emission spectral profiles previously created in Zen (Maynard et al., 2019), stitched, maximum intensity projected, and saved as Carl Zeiss Image “*.czi*” files.

### snRNA-seq spatial registration

For each snRNA-seq dataset, we utilized publicly-available processed unique molecular index (UMI) count data for each gene and nucleus, and provided annotations of cell clusters/subtypes. Within each dataset, we performed ‘pseudo-bulking’ (Crowell et al., 2019; Kang et al., 2018; Lun and Marioni, 2017) of nuclei-level UMIs into cell type-specific log-transformed normalized counts for each unique subject. We then computed cell type ‘enrichment’ statistics for each gene and dataset-provided cell type within their pseudo-bulk profiles by performing linear mixed effects modeling comparing each cell type to all other cell types, treating donor as a random intercept (Law et al., 2014), and adjusting for study-specific covariates described below. This strategy was analogous to the layer ‘enrichment’ statistics described for our Visium data (Method Details: Layer-level gene modeling). We then computed Pearson correlation coefficients between our layer-enriched ‘enrichment’ statistics and snRNA-seq cell type-specific “enrichment” statistics among the 700 most layer-enriched genes (combining the 100 most significant genes for each of the six layers and WM in the Visium data) expressed in each snRNA-seq dataset. In addition to our DLPFC snRNA-seq dataset (Method Details: DLPFC snRNA-seq data generation), we utilized these publicly-available datasets:

1. Hodge *et al*. (Hodge et al., 2019): Processed data was obtained from https://portal.brain-map.org/atlases-and-data/rnaseq. We retained total gene counts (exons plus introns) from 49,494 nuclei corresponding to postmortem human brain tissue across both neurons and non-neurons across 50,281 genes across 6 layers and 2 cell types. These data were reduced to 52 pseudo-bulk profiles, for all unique donor-layer-type combinations. We calculated ‘enrichment’ statistics for each of the six layers in their dataset, adjusting for the fixed effect of cell type (neuronal or glial) with a random intercept of donor.
2. Velmeshev *et al*. (Velmeshev et al., 2019): Processed data was obtained from https://cells.ucsc.edu/ (under the “Autism” study data download). We used the post-QC UMI counts from all 104,559 nuclei across 65,217 genes across 41 unique donor-region pairs (for 31 unique donors and 2 brain regions) and 17 cell types. These data were reduced to 691 pseudo-bulked profiles, for all unique donor-region-type combinations. We calculated ‘enrichment’ statistics for each of the 17 cell types in this dataset, adjusting for the fixed effect of brain region, age, sex, and ASD diagnosis, with a random intercept of donor.
3. Mathys *et al*. (Mathys et al., 2019): Processed data were obtained from Synapse at accession: syn18485175. We used the post-QC UMI counts from all 70,634 nuclei across 17,926 genes across 48 unique donors and 44 cell subtypes (across 8 broad cell classes). These data were reduced to 1877 pseudo-bulked profiles, for all unique donor-subtype combinations. We calculated ‘enrichment’ statistics for each of the 44 cell subtypes in this dataset, adjusting for the fixed effect of age, sex, race, and Alzheimer’s disease diagnosis, with a random intercept of donor.
4. We also downloaded and reprocessed RNA-seq data from He *et al*. (He et al., 2017) from SRA accession SRP199498 using our previously-described RNA-seq processing pipeline (Collado-Torres et al., 2019). These data consisted of homogenate RNA-seq data from 18 serial sections across 4 unique donors.

### DLPFC snRNA-seq data generation

We performed single-nucleus RNA-seq (snRNA-seq) on DLPFC tissue from two neurotypical donors using 10x Genomics Chromium Single Cell Gene Expression V3 technology. Nuclei were isolated using a “Frankenstein” nuclei isolation protocol developed by Martelotto *et al*. for frozen tissues (Habib et al., 2016, 2017; Hu et al., 2017; Lacar et al., 2016; Lake et al., 2016). Briefly, ∼40mg of frozen DLPFC tissue was homogenized in chilled Nuclei EZ Lysis Buffer (MilliporeSigma) using a glass dounce with ∼15 strokes per pestle. Homogenate was filtered using a 70μm-strainer mesh and centrifuged at 500 x g for 5 minutes at 4°C in a benchtop centrifuge. Nuclei were resuspended in the EZ lysis buffer, centrifuged again, and equilibrated to nuclei wash/resuspension buffer (1x PBS, 1% BSA, 0.2U/μL RNase Inhibitor).

Nuclei were washed and centrifuged in this nuclei wash/resuspension buffer three times, before labeling with DAPI (10μg/mL). The sample was then filtered through a 35μm-cell strainer and sorted on a BD FACS Aria II Flow Cytometer (Becton Dickinson) at the Johns Hopkins University Sidney Kimmel Comprehensive Cancer Center (SKCCC) Flow Cytometry Core into 10X Genomics reverse transcription reagents. Gating criteria hierarchically selected for whole, singlet nuclei (by forward/side scatter), then for G_0_/G_1_ nuclei (by DAPI fluorescence). A “null” sort into wash buffer was additionally performed from the same preparation for quantification of nuclei concentration and to ensure nuclei input was free of debris. Approximately 8,500 single nuclei were sorted directly into 25.1μL of reverse transcription reagents from the 10x Genomics Single Cell 3’ Reagents kit (without enzyme). Libraries were prepared according to manufacturer’s instructions (10x Genomics) and sequenced on the Next-seq (Illumina) at the Johns Hopkins University Transcriptomics and Deep Sequencing Core.

We processed the sequencing data with the 10x Genomics’ Cell Ranger pipeline, aligning to the human reference genome GRCh38, with a reconfigured GTF such that intronic alignments were additionally counted given the nuclear context, to generate UMI/feature-barcode matrices (https://support.10xgenomics.com/single-cell-gene-expression/software/pipelines/latest/advanced/references). We started with raw feature-barcode matrices for analysis with the Bioconductor suite of R packages for single-cell RNA-seq analysis (Amezquita et al., 2020). For quality control and cell calling, we first used a Monte Carlo simulation-based approach to assess and rule out empty droplets or those with random ambient transcriptional noise, such as from debris (Griffiths et al., 2018; Lun et al., 2019). This was then followed by mitochondrial rate adaptive thresholding, which, though expected to be near-zero in this nuclear context, we allowed for a 3x median absolute deviation (MAD) threshold. This allowed for flexibility in output/purity of FACS workflows. This QC pipeline yielded 5,399 high-quality nuclei from the DLPFC from two donors, which were then rescaled across all nuclear libraries, then log-transformed for determination of highly-variable genes, again with *scran*’s `modelGeneVar`, this time taking all genes (9,313) with a greater variance than the fitted trend. Principal components analysis (PCA) was then performed on these selected genes to reduce the high dimensionality of nuclear transcriptomic data. The optimal PC space was defined with iterative graph-based clustering to determine the *d* PCs where resulting *n* clusters stabilize, with the constraint that *n* clusters </= (*d* + 1) PCs (Lun et al., 2016), resulting in a chosen *d*=81 PCs. In this PCA-reduced space, graph-based clustering was performed (specifically, k-nearest neighbors with k=20 neighbors and the Walktrap method from R package *igraph* (Csardi and Nepusz, 2006) for community detection) to identify 31 preliminary clusters. We then took all feature counts for these assignments and pseudo-bulked counts across 31 preliminary nuclear clusters, rescaling for combined library size and log-transforming normalized counts, then performed hierarchical clustering to identify preliminary cluster relationships and merging with the `cutreeDynamic` function of R package *dynamicTreeCut* (Langfelder et al., 2016). These broader clusters were finally annotated with well-established cell type markers for nuclear type identity (Mathys et al., 2019). We also used Bioconductor package *scater*’s (McCarthy et al., 2017) implementation of non-linear dimensionality reduction techniques, *t*-SNE and UMAP, with default parameters and within the aforementioned optimal PC space, simply for visualization of the high-dimensional structure in the data, which complemented the clustering results (**Figure S14**).

### Clinical gene set enrichment analyses

We assessed the laminar enrichment of a series of predefined clinical gene sets for various neuropsychiatric and neurodevelopmental disorders. These gene sets consisted of data from:

1. Birnbaum *et al*. (Birnbaum et al., 2014): 10 gene sets across SCZ, ASD, neurodevelopmental disorders, intellectual disability, bipolar disorder, and neurodegenerative disorders.
2. SFARI (Abrahams et al., 2013): 3 gene sets consisting of all human genes, high confidence genes, and syndromic genes.
3. Satterstrom *et al*. (Satterstrom et al., 2020): 6 gene sets based on exome sequencing studies.
4. psychENCODE (Gandal et al., 2018): 6 gene sets based on DE analyses of patients with ASD, SCZD, and bipolar disorder (BPD), stratified by directionality in cases, and 8 gene sets based on TWAS (ASD, SCZD, BPD, SCZD-BPD, stratified by directionality), each at FDR < 0.05.
5. BrainSeq (Collado-Torres et al., 2019): 2 gene sets based on DE analyses of patients with SCZD versus controls (at FDR < 0.05), stratified by directionality.
6. Down syndrome (Olmos-Serrano et al., 2016): 2 gene sets based on DE analyses of patients with Down syndrome versus controls (at FDR < 0.05), stratified by directionality.

We collected all reported genes in each gene set, and retained the majority that were expressed in our Visium dataset - these gene set sizes are provided in **Table S6**. Enrichment for each gene set for each layer was based on a gene being significantly more highly expressed in one layer versus all other layers (at FDR < 0.1). This calculation was performed using Fisher’s exact test, which returned an odds ratio and *p*-value for each gene set and layer (**Table S6**). Pooling all *p*-values resulted in FDR control of 5% for marginal *p*-values < 0.01.

We additionally performed MAGMA (de Leeuw et al., 2015) using the subset of 24,347 Ensembl gene IDs expressed in our pseudo-bulked Visium data that were present in the provided GR37/hg19 annotation across multiple GWASs for SCZD (Pardiñas et al., 2018), BPD (Stahl et al., 2019), MDD (Wray et al., 2018) and ASD (Grove et al., 2019). We used window sizes of +35kb and −10kb around each gene to aggregate SNPs to genes using the 1000 Genomes EUR reference profile using SNP-wise stats. We then performed gene set testing using MAGMA for seven gene sets (related to the six layers and WM) for genes with positive (+) enrichment statistics at FDR < 0.1. Additionally, we performed linkage disequilibrium score regression (LDSC) and partitioned heritability analysis (Bulik-Sullivan et al., 2015; Finucane et al., 2015) using 30 GWAS traits collected by Rizzardi *et al* (Rizzardi et al., 2019). Genomic regions were created from the same enriched and FDR < 0.1 genes as above, here with +10kb and −5kb windows, and lifted over to hg19 coordinates.

### Data-driven layer-enriched clustering analysis

For the data-driven layer-enriched clustering, we first performed feature selection in two ways to identify laminar and non-laminar patterns in our data. The first method for feature selection used was *SpatialDE* (Svensson et al., 2018) to identify genes exhibiting spatially variable expression patterns (SVGs). *SpatialDE* was run in Python version 3.8.0. We ran *SpatialDE* individually on each of the 12 samples, which returned a set of statistically significant (false discovery rate < 0.05) SVGs per sample. We included an additional filtering step to remove lowly-expressed genes (less than 1,000 total UMIs summed across spots per sample), as well as removing mitochondrial genes. This left between 521 and 2,217 genes per sample (**Table S9**). In total, there were 2,775 unique genes across samples; for comparison, we also ran clustering methods using this pooled list (**Table S9** and **Table S10**). The second feature selection method used the *scran* R Bioconductor package (Lun et al., 2016) to identify (non-spatial) highly variable genes (HVGs) across all samples combined, which identified 1,942 HVGs. Due to slow runtime, it was not possible to run *SpatialDE* on pooled spots from all samples combined.

In the ‘unsupervised’ approach to define sub-groups of spots with similar expression profiles in a completely data-driven manner, we considered the possible combinations of (i) two types of methods for dimensionality reduction (top 50 principal components (PCs) with the *BiocSingular* Bioconductor package (Lun, 2019), and top 10 UMAP (McInnes et al., 2018) components with the *uwot* R package (Melville, 2019) calculated on the top 50 PCs), (ii) the gene sets defined after applying feature selection (*SpatialDE* genes for each sample, pooled *SpatialDE* gene lists across all 12 samples, and HVGs), and (iii) including (or not) the two spatial coordinates (*x* and *y* coordinates) of each spot as additional features for clustering. For the clustering algorithm, we constructed a shared nearest neighbor graph with the *scran* Bioconductor package and then applied the Walktrap method from the *igraph* R package (Csardi and Nepusz, 2006) to obtain predicted cluster labels. We set all clustering implementations to return eight final clusters (i.e. one more than the six DLPFC layers plus white matter), which gave slightly improved clustering performance (compared to seven clusters) due to additional splitting of the white matter cluster and some outlier spots. **Table S10** contains an overview of all combinations that were tried.

For comparison, we also implemented a ‘semi-supervised’ approach, where we used the layer-enriched gene sets identified using the DE “enrichment” models described previously (**Figure S7**), and a ‘markers’ approach using known marker genes from Zeng *et al*. (Zeng et al., 2012) (**Table S10**).

To evaluate the performance of the clustering approaches, we used the adjusted Rand index (ARI), which measures the similarity between the predicted cluster labels and “gold standard” cluster labels. The manually-annotated layers were used as the “gold standard” (Figure 7 and **Figure S17**). Higher ARI values correspond to better clustering performance, with a maximum value of 1 indicating perfect clustering agreement. To evaluate the improvement in ARI when including spatial coordinates within the clustering methods, we fit a linear model on the ARI scores, comparing these methods against methods without spatial coordinates across all methods and samples, and recorded the *p*-value.

### QUANTIFICATION AND STATISTICAL ANALYSIS

The different subsections of the “Method Details” further specify the statistical models and tests used as well as the versions of the specific software used. Overall, statistical tests were performed using R versions 3.6.1 and 3.6.2 with Bioconductor version 3.10 (current release version) with detailed R session information provided in the code GitHub repositories listed under “Data and Software Availability.” The threshold and method used for statistical significance is listed in the main text along the description of the results. Plots in R were created in either base R or with the *ggplot2* R package (Wickham, 2016).

### DATA AND SOFTWARE AVAILABILITY

Raw and processed data is available from *ExperimentHub* (Bioconductor Package Maintainer, 2017) as well as the Bioconductor package *spatialLIBD*. Code is available through GitHub at https://github.com/LieberInstitute/HumanPilot (Collado-Torres et al., 2020) and https://github.com/LieberInstitute/spatialLIBD (Collado-Torres, 2020), both of which are described in their README.md files.

### ADDITIONAL RESOURCES

In order to visualize the spot-level Visium data we generated, we created a *shiny* (Chang et al., 2019) interactive browser available at http://spatial.libd.org/spatialLIBD that is powered by the Bioconductor package *spatialLIBD* (Collado-Torres, 2020).

## Supplemental Information

**Table S1. Sample metrics from Space Ranger and demographic information, related to Figure 1**. A tab-separated table with the sequencing and alignment metrics produced by Space Ranger as well as the age, sex and LIBD brain ID for the samples sequenced in this project.

**Table S2. Percent of spots with zero or one cell across layers, related to Results: Gene expression in the DLPFC across cortical laminae.** This table shows the percentage of spots in each annotated layer with zero or one segmented cells.

**Table S3. Spot cell count differential expression statistics, related to Results: Gene expression in the DLPFC across cortical laminae.** Differential expression statistics comparing spots with 0 cells to >0 cells. Positive log2 fold changes indicate higher expression in spots without cells.

**Table S4. Layer level differential expression statistics, related Figure 2**. Differential expression statistics for the (**A**) ANOVA model (one model per gene), (**B**) Enrichment model (7 models per gene, 1 per layer), and (**C**) Pairwise model (21 models per gene, 1 per pair of layers).

**Table S5. Optimal model results for known layer marker genes, related to Figure 3**. Differential expression statistics for the optimal model for each known human or mouse brain marker gene as well as the top ranked gene using the layer-level data.

**Table S6. Clinical gene sets layer enrichment statistics, related to Figure 6**. Each row is a different gene set obtained from the literature. PE: psychENCODE, BS: BrainSeq, DS: Down Syndrome, DE: Differential Expression, TWAS: transcriptome-wide association study, OR: odds ratio, NumSig: number of significant layer-enriched genes in the gene set for that particular layer.

**Table S7. Clinical gene set enrichment results with MAGMA, related to Figure 6**. P-values for MAGMA gene set test for layer-enriched genes across four GWAS for SCZD, MDD, ASD and BPD. Bold indicates FDR < 0.05 significance and red indicates Bonferroni < 0.05 significance.

**Table S8. LDSC results, related to Figure 6**. Genomic enrichments of GWAS risk SNPs using partitioned heritability analysis. Prop = proportion, h2 = heritability, p = p-value, holm = Holm’s adjusted p-values.

**Table S9. Summary of *SpatialDE* genes, related to Figure 7**. Number of statistically significant spatially variable genes (SVGs) identified per sample using *SpatialDE* (Svensson et al., 2018), before and after additional filtering for lowly-expressed genes and mitochondrial genes.

**Table S10. Description of clustering methods used for the data-driven layer-enriched clustering analyses, related to Figure 7**. Summary of combinations of design choices that were implemented for the clustering methods used for the data-driven spatial clustering analyses. Columns describe: (i) method names, (ii) the type of clustering method, (iii) the type of dimension reduction used to summarize gene expression, (iv) the source of gene sets used, and (v) whether spatial coordinates were included as additional features for clustering. The names of the clustering methods correspond to those shown in **Figure S18**, **Figure S19**, and Supplementary File 1.

**Supplementary File 1.** Visualization of clustering results for the data-driven layer-enriched clustering analyses, for all samples and clustering methods; related to **Figure 7**. This supplementary file contains visualizations of clustering results for all samples and clustering methods (Table S10) (similar to **Figure S18** and **Figure S19**, which display results for sample 151673 only). A description of the clustering methods is provided in Table S10.

---

**Figure S1.**
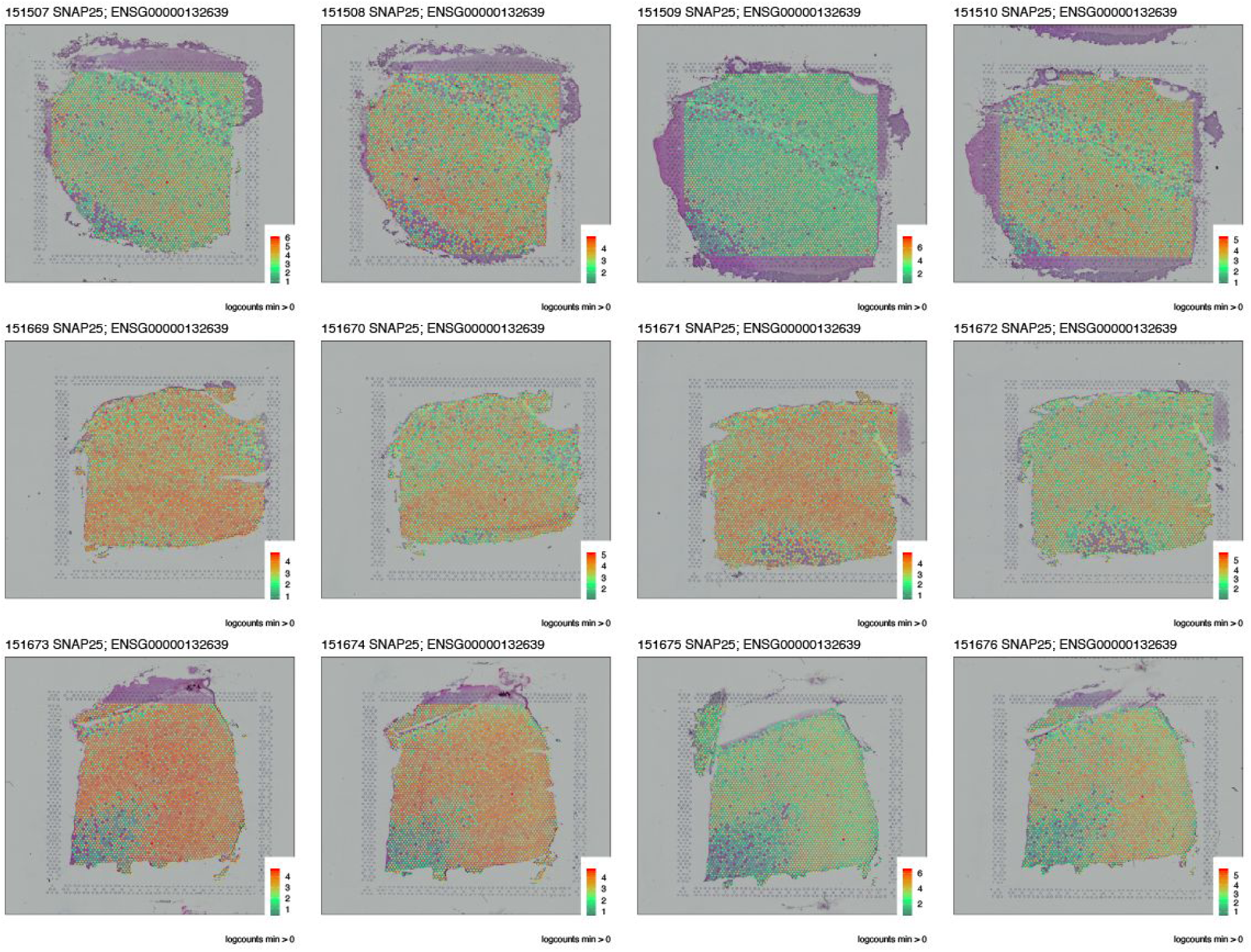
*SNAP25* expression, related to. Figure 1D. Log-transformed normalized (logcounts) for *SNAP25* gene expression across all 12 samples arranged in rows by subject.

**Figure S2.**
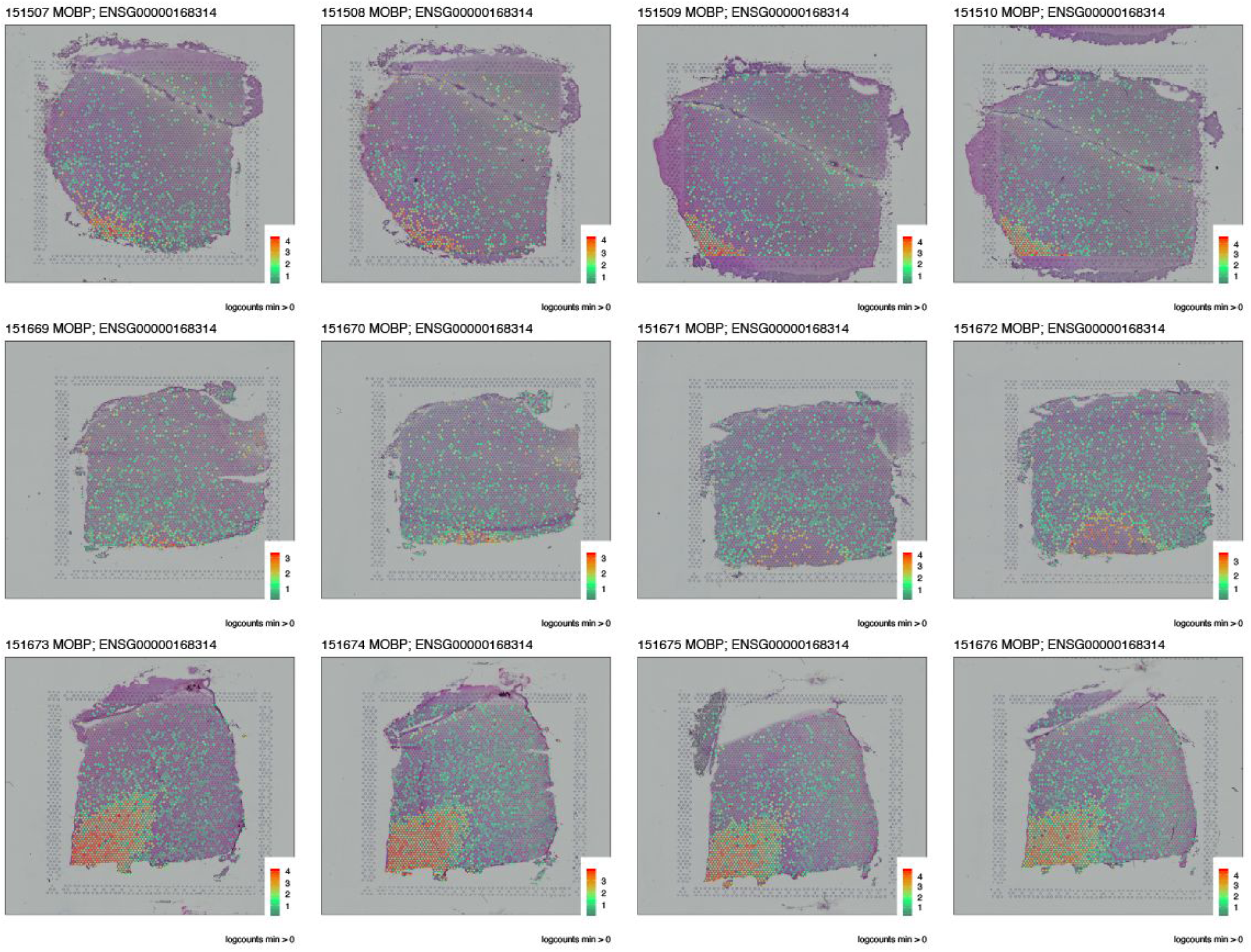
*MOBP* expression, related to. Figure 1E. Log-transformed normalized (logcounts) for *MOBP* gene expression across all 12 samples arranged in rows by subject.

**Figure S3.**
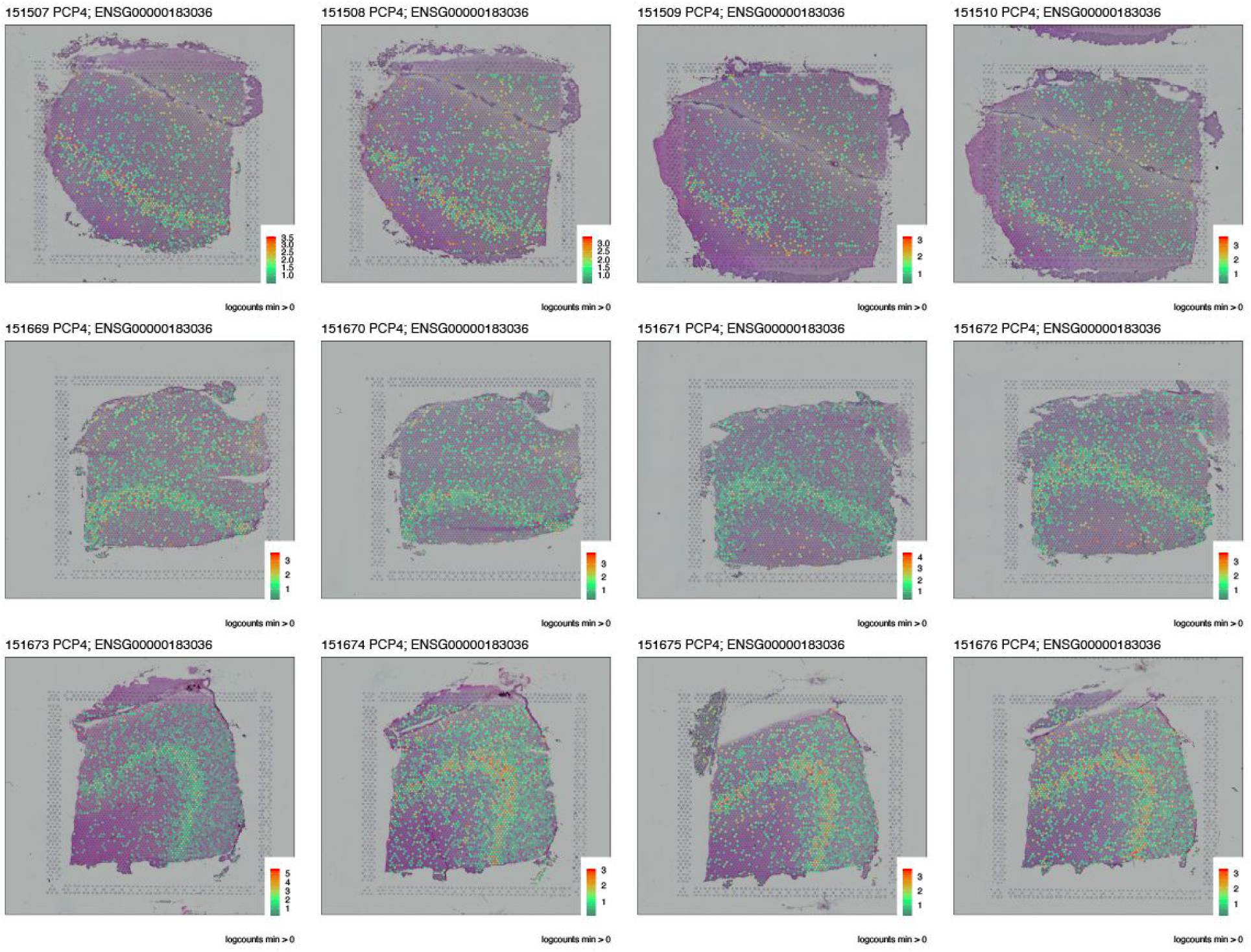
*SNAP25* expression, related to. Figure 1F. Log-transformed normalized (logcounts) for *MOBP* gene expression across all 12 samples arranged in rows by subject.

**Figure S4.**
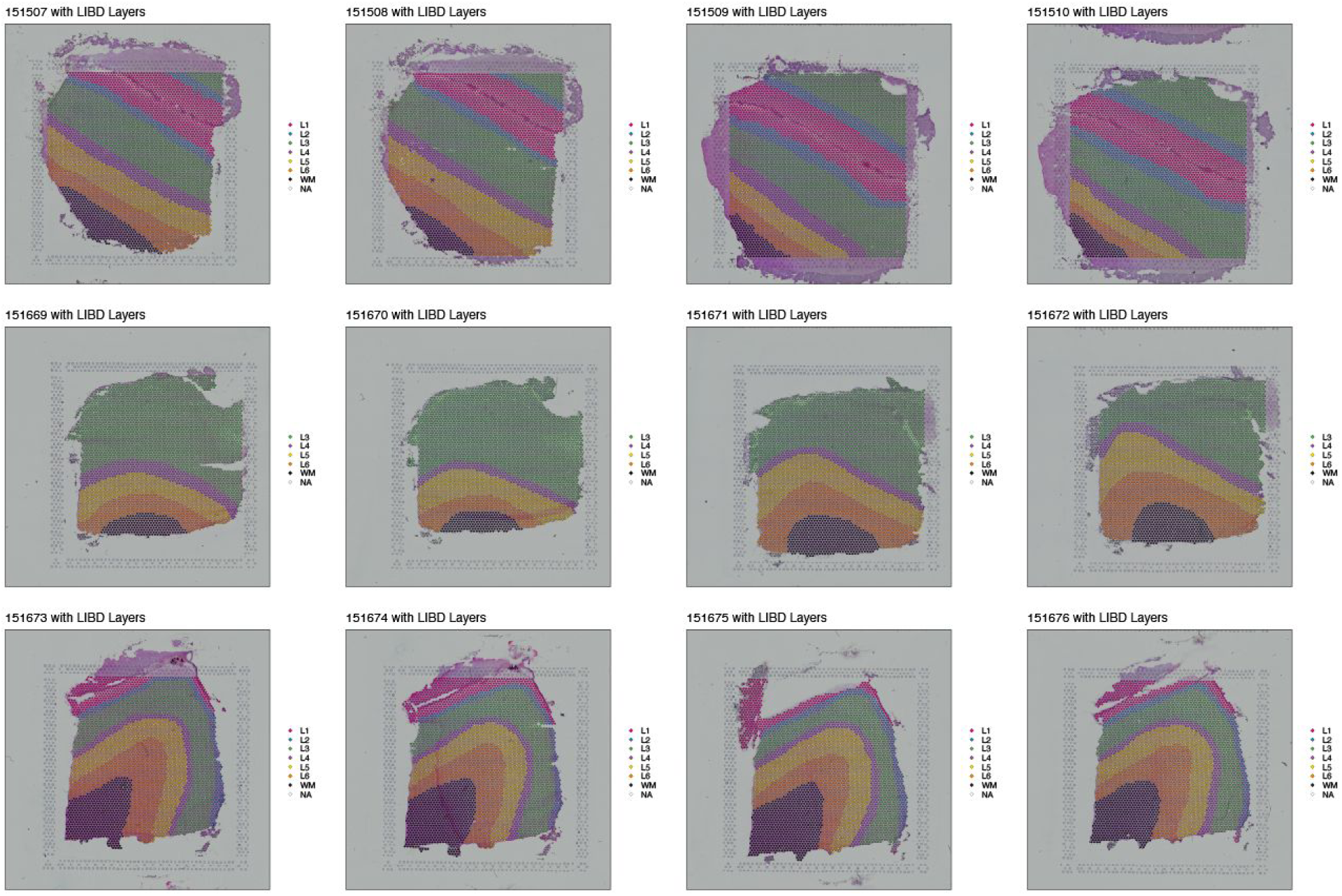
Supervised annotation of layers based on cytoarchitecture and known marker gene expression, related to. Figure 2. Manual annotation of cortical layers across all 12 samples arranged in rows by subject. See also Figure S5.

**Figure S5.**
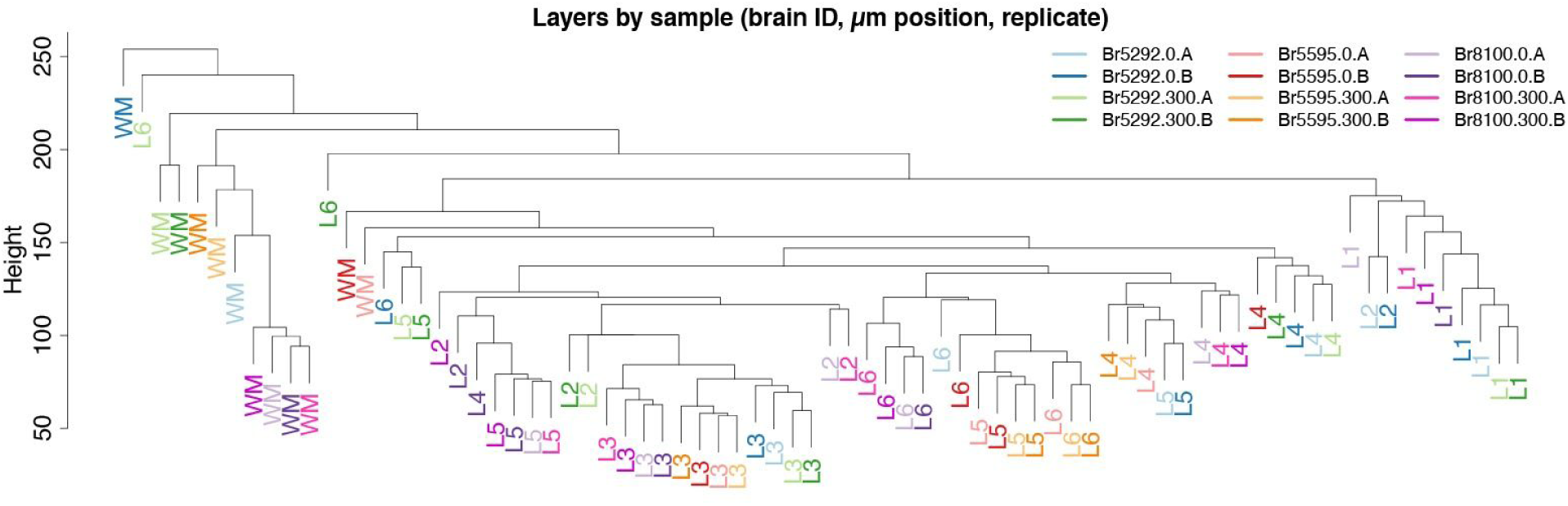
Layer-level dendrogram, related to Results: Gene expression in the DLPFC across cortical laminae and. Figure 2. Dendrogram from the hierarchical clustering performed across all 76 layer-level combinations: 6 layers plus WM across 12 samples, with two layers visually absent in one sample as shown in **Figure S4**, second row. The layer-level combinations are colored by the brain subject (BR5292, Br5595, Br8100), position (0 or 300) and adjacent spatial replicate number (A or B).

**Figure S6.**
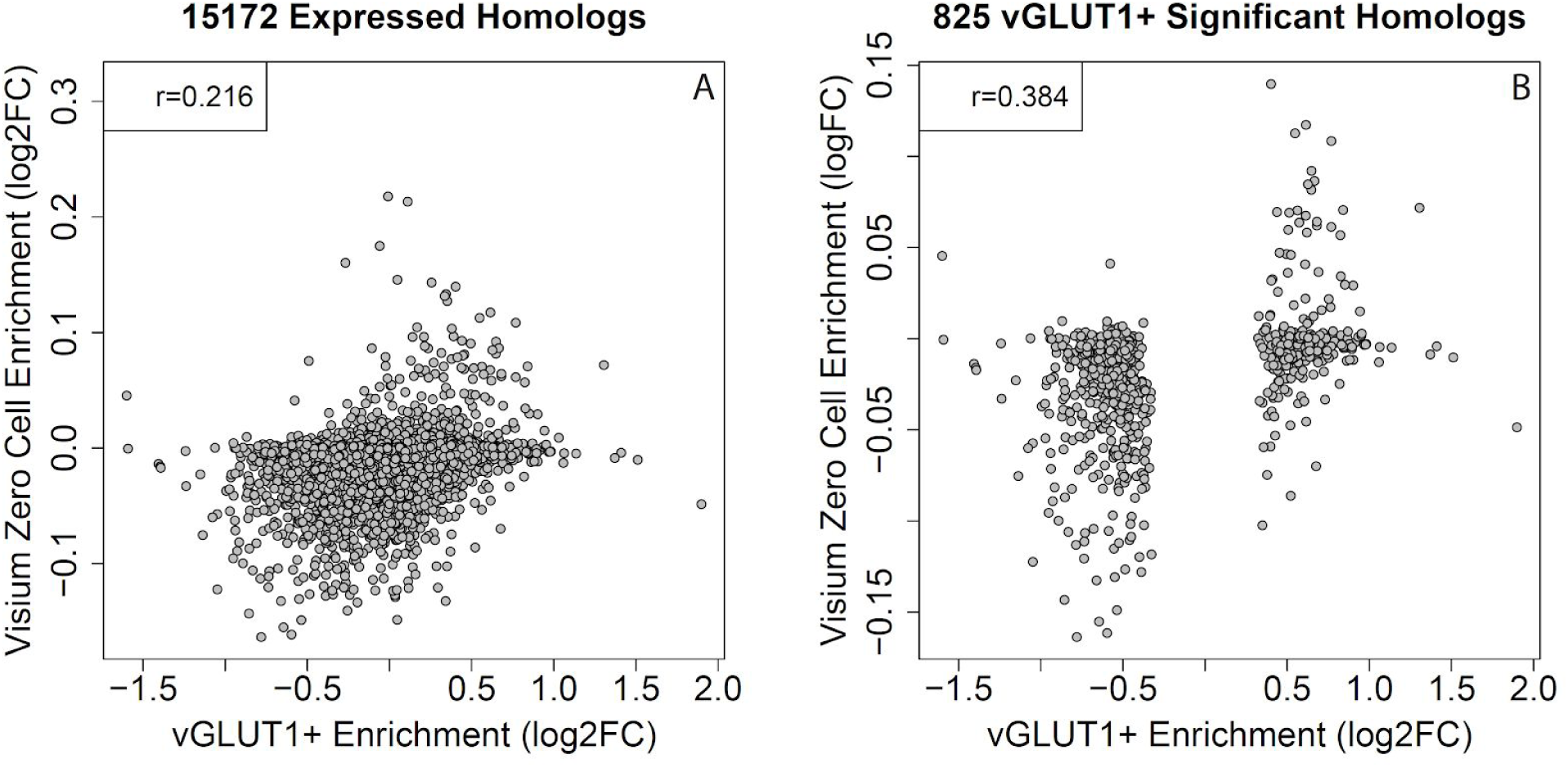
Enrichment of genes expressed in synaptic terminals among neuropil spots, related to Results: Gene expression in the DLPFC across cortical laminae. We compared DEGs from VGLUT1+ labeled synaptosomes from mouse brain from Hafner *et al* (Hafner et al., 2019) on the x-axis versus the log2 fold change comparing spot-level expression between spots with 0 cells and spots with >0 cells. Association shown between (A) all expressed homologous genes and (B) those genes that were significant in the Hafner *et al*. dataset at FDR < 0.05.

**Figure S7.**
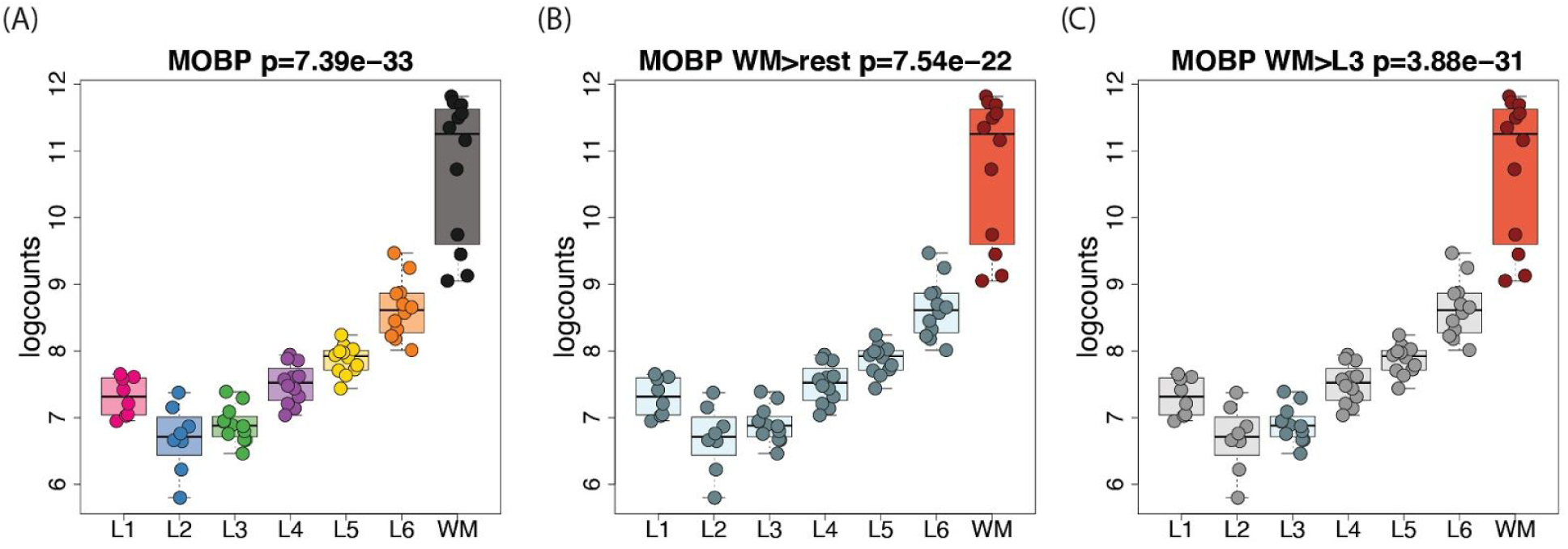
Layer-level modeling strategies illustrated with *MOBP*, related to Results: Figure 2. Overview of the different modeling strategies we performed with the layer-level pseudo-bulked expression data. (**A**) The *ANOVA* model, which evaluates whether the gene is variable in any of the layers (F-statistic); *MOBP* is the top 10th ranked of such genes. Colors represent each layer. (**B**) The *enrichment* model, which tests one layer against the rest (t-statistic); *MOBP* is the top 36th gene for white matter against other layers. Colors show the comparison being done. (**C**) The *pairwise* model where we test one layer against another (t-statistic); *MOBP* is the top ranked gene for WM > L3. Data from layers not used is shown in gray.

**Figure S8.**
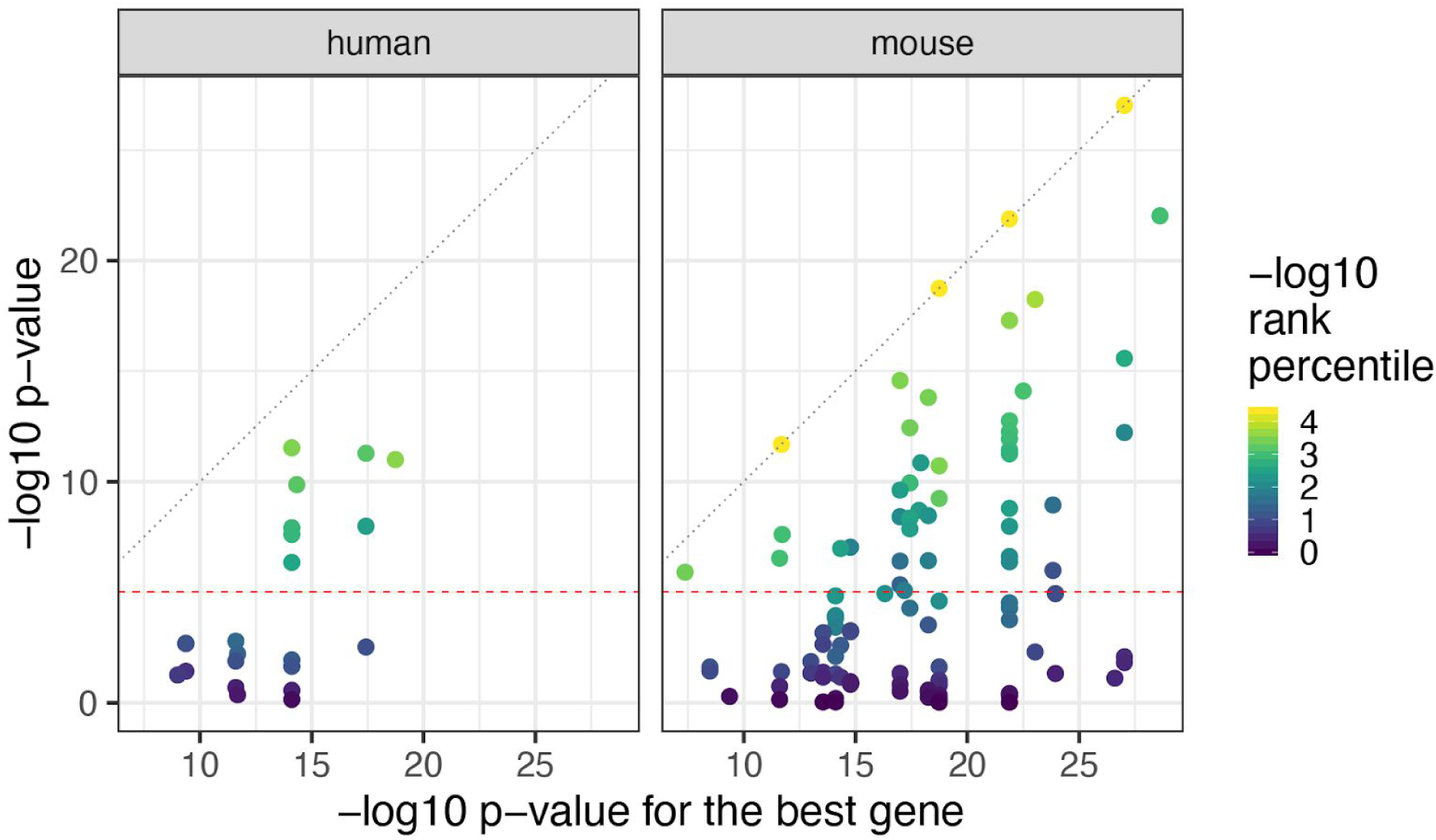
Known marker genes compared to the best gene, related to Results: Identifying novel layer-enriched genes in human cortex. Using the optimal models (Method Details: Known marker genes optimal modeling) for each known marker gene we compared the marker genes against the best gene for that given model. Results are visualized using the -log10 p-values for the marker gene (y-axis) against the best gene for that model (x-axis). Points are colored by the -log10 rank percentile of that gene in such a way that the top ranked gene is -log10(1 / 22,331) and colored in yellow.

**Figure S9.**
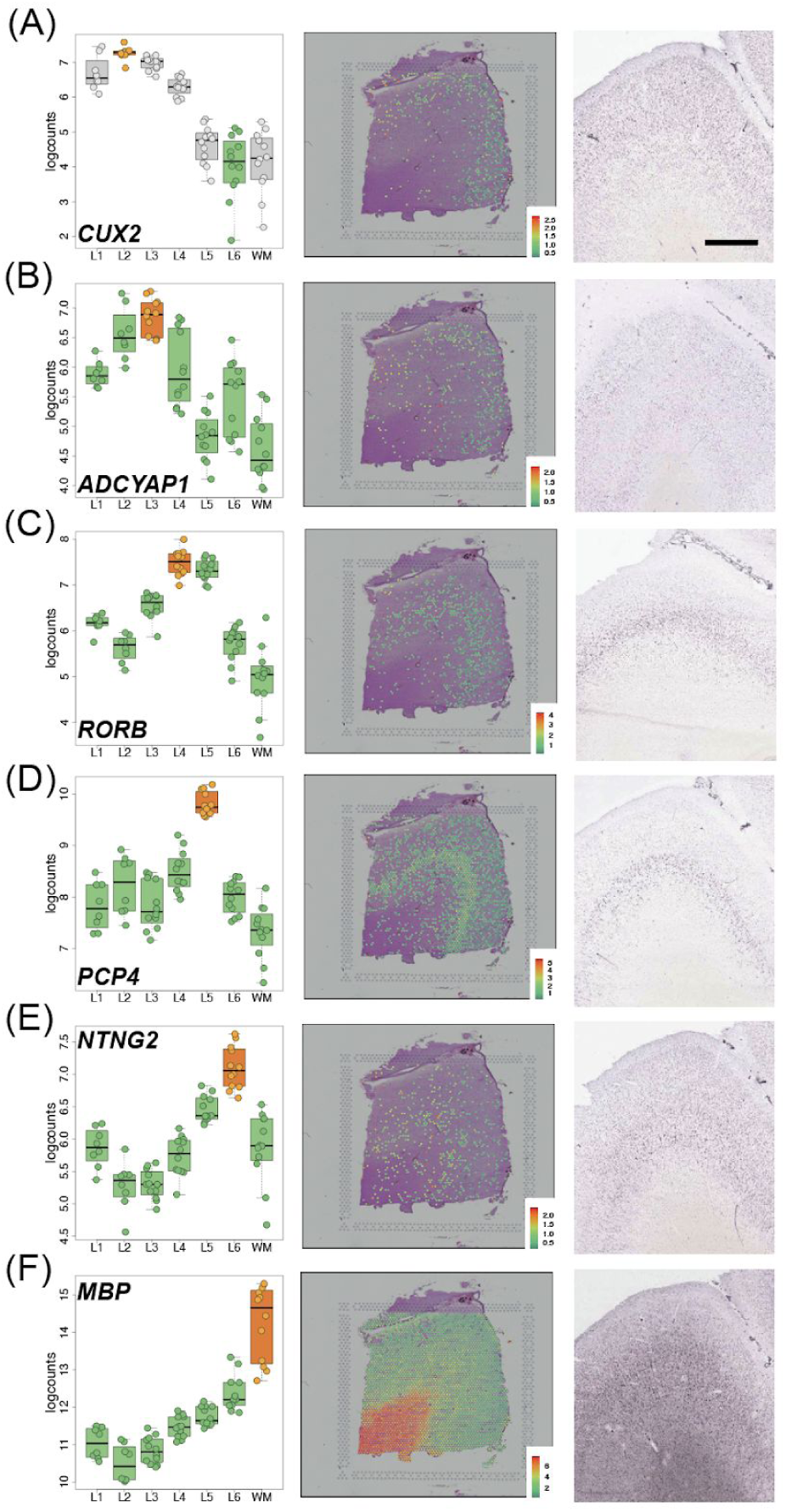
Replication of Visium layer-enriched genes by Allen Brain Atlas *in situ* hybridization (ISH) data, Related to. Figure 3. **(A-F)** Left panels: Boxplots of log-transformed normalized expression (logcounts) for genes *CUX2* (**A**, L2>L6, *p*=3.75e-19), *ADCYAP1* (**B**, L3>rest, *p*=3.57e-08), *RORB* (**C**, L4>rest, *p*=2.91e-07), *PCP4* (**D**, L5>rest, *p*=1.81e-19), *NTNG2* (**E**, L6>rest, *p*=5.22e-13), and *MBP* (**F**, WM>rest, *p*=1.71e-20). Middle panels: Spotplots of log-transformed normalized expression (logcounts) for sample 151673 for *CUX2* (**A**), *ADCYAP1* (**B**), *RORB* (**C**), PCP4 (**D**), *NTNG2* (**E**), and *MBP* (**F**). Right panels: *in situ* hybridization (ISH) images from DLPFC (**A, C, D, E, F**) or frontal cortex (**B**) of adult human brain from Allen Brain Institute’s Human Brain Atlas: http://human.brain-map.org/ (Hawrylycz et al., 2012). Scale bar for Allen Brain Atlas ISH images=1.6mm.

**Figure S10.**
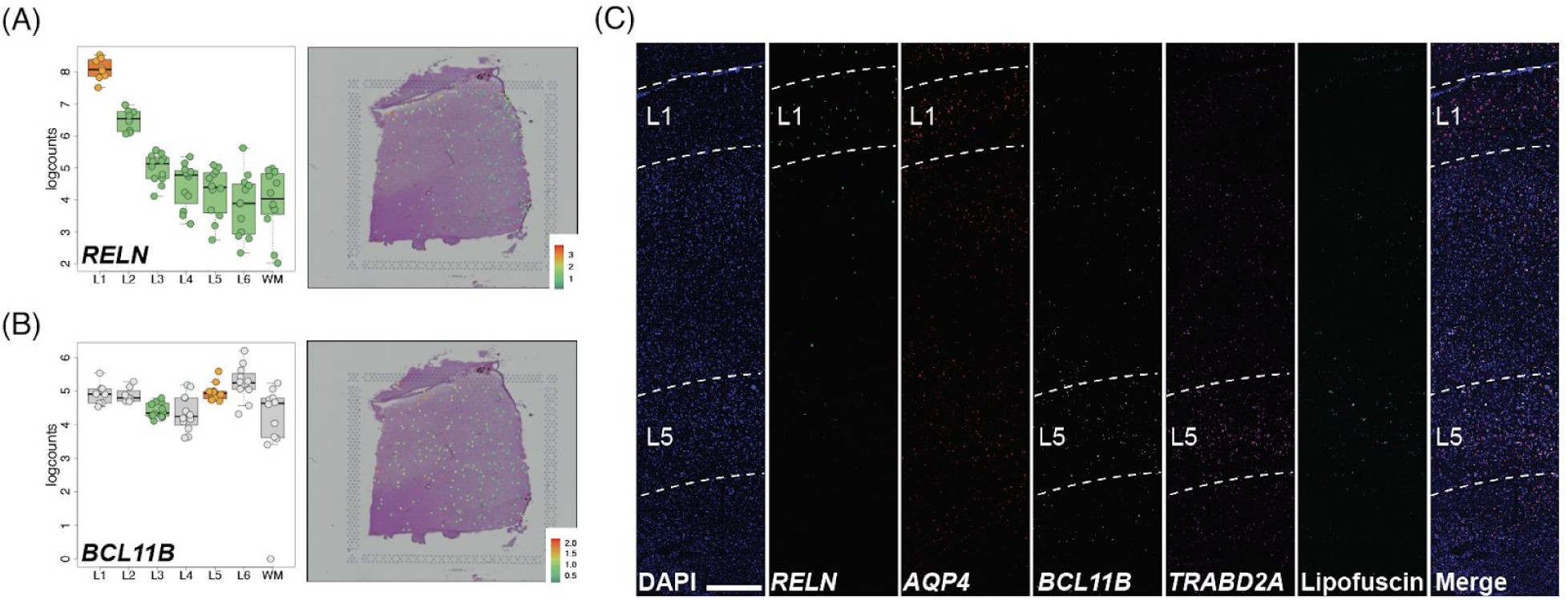
smFISH validation of L1- and L5-enriched genes, related to. Figure 4. **(A-B)** Left panels: Boxplots of log-transformed normalized expression (logcounts) for previously identified L1 and L5 marker genes RELN (**A**, L1>rest, *p*=7.94e-15,) and *BCL11B* (**B**, L5>L3, *p*=4.44e-02), respectively. Right panels: Spotplots of log-transformed normalized expression (logcounts) for sample 151673 for genes RELN (**A**) and *BCL11B* (**B**). Corresponding boxplots and spotplots for Visium-identified genes *AQP4* and *TRABD2A* in Figure 4. (**C**) Multiplex single molecule fluorescent in situ hybridization (smFISH) in a cortical strip of DLPFC. Maximum intensity confocal projections depicting expression of DAPI (nuclei), *RELN* (L1), *AQP4* (L1), *BCL11B* (L5)*, TRABD2A* (L5) and lipofuscin autofluorescence. Merged image without lipofuscin autofluorescence. Scale bar=500μm.

**Figure S11.**
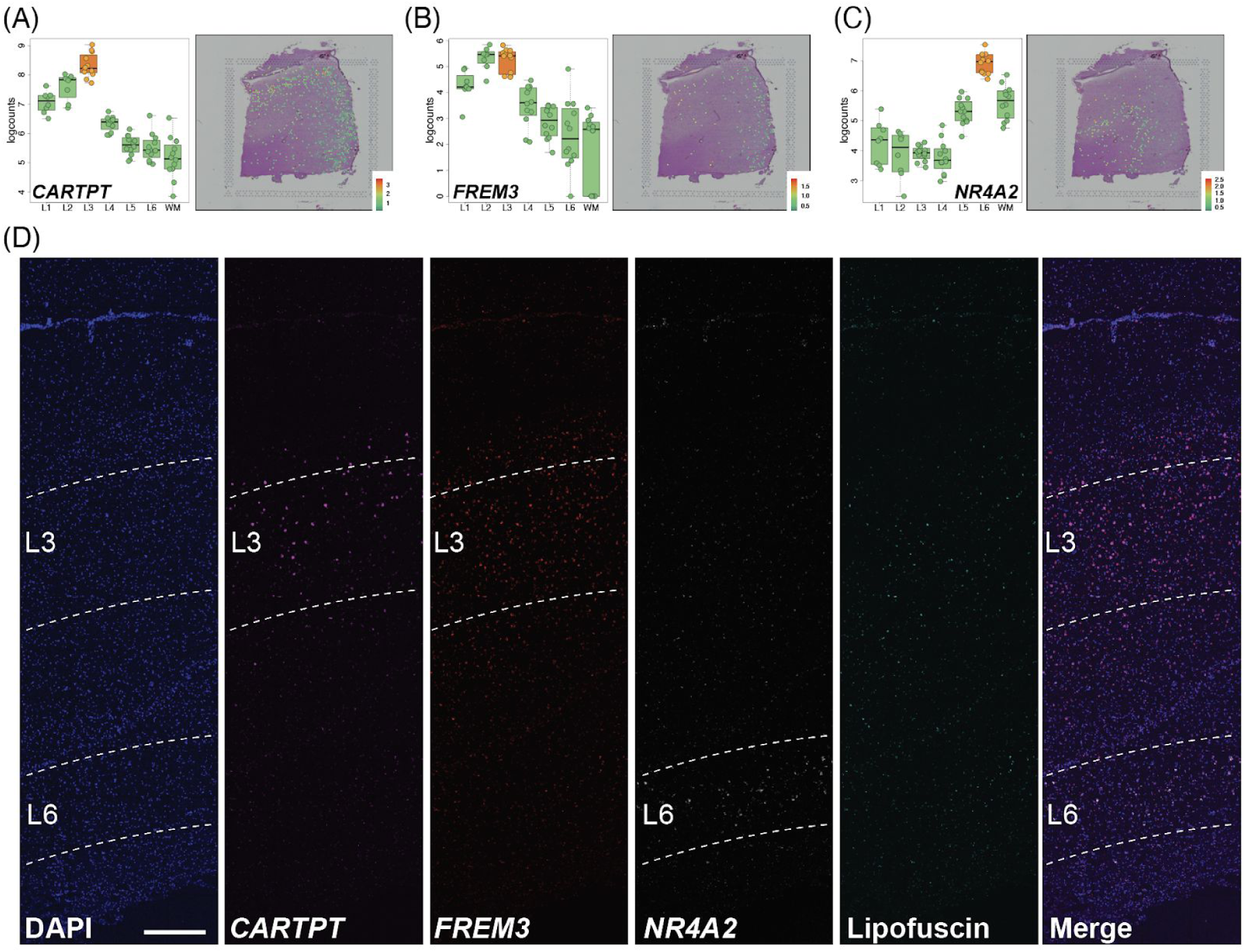
smFISH validation of L3- and L6-enriched genes, related to. Figure 4**. (A-C)** Left panels: Boxplots of log-transformed normalized expression (logcounts) for previously identified L3 and L6 marker genes CARTPT (**A**, L3>rest, *p*=2.07e-12) and *NR4A2* (**C**, L6>rest, *p*=1.15e-13), respectively, and Visium-identified gene L3 gene *FREM3* (**B**, L3>rest, *p*=8.16e-07). Right panels: Spotplots of log-transformed normalized expression (logcounts) for sample 151673 for corresponding genes. (**D**) Multiplex single molecule fluorescent in situ hybridization (smFISH) in a cortical strip of DLPFC. Maximum intensity confocal projections depicting expression of DAPI (nuclei), *CARTPT* (L3), *FREM3* (L3), *NR4A2* (L6) and lipofuscin autofluorescence. Merged image without lipofuscin autofluorescence. Scale bar=500 μm.

**Figure S12.**
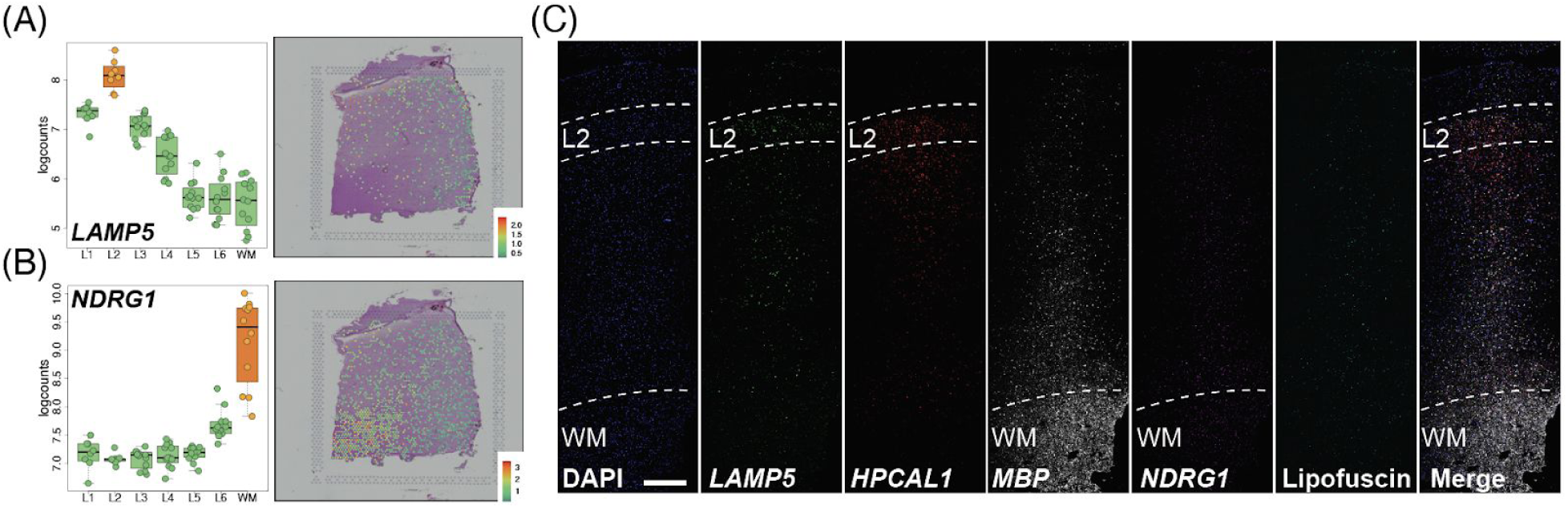
smFISH validation of L2- and WM-enriched genes, related to. Figure 4. **(A-B)** Left panels: Boxplots of log-transformed normalized expression (logcounts) for Visium-identified L2 and WM genes *LAMP5* (**A**, L2>rest, *p*=2.60e-09) and *NDRG1* (**B**, WM>rest, *p*=1.26e-26), respectively. Right panels: Spotplots of log-transformed normalized expression (logcounts) for sample 151673 for *LAMP5* (**A**) and *NDRG1* (**C**). Corresponding boxplots and spotplots for *HPCAL1* in Figure 4 and *MBP* in **Figure S9**. (**D**) Multiplex single molecule fluorescent in situ hybridization (smFISH) in a cortical strip of DLPFC. Maximum intensity confocal projections depicting expression of DAPI (nuclei), *LAMP5* (L2), *HPCAL1* (L2), *MBP* (WM)*, NDRG1* (WM) and lipofuscin autofluorescence. Merged image without lipofuscin autofluorescence. Scale bar=500 μm.

**Figure S13.**
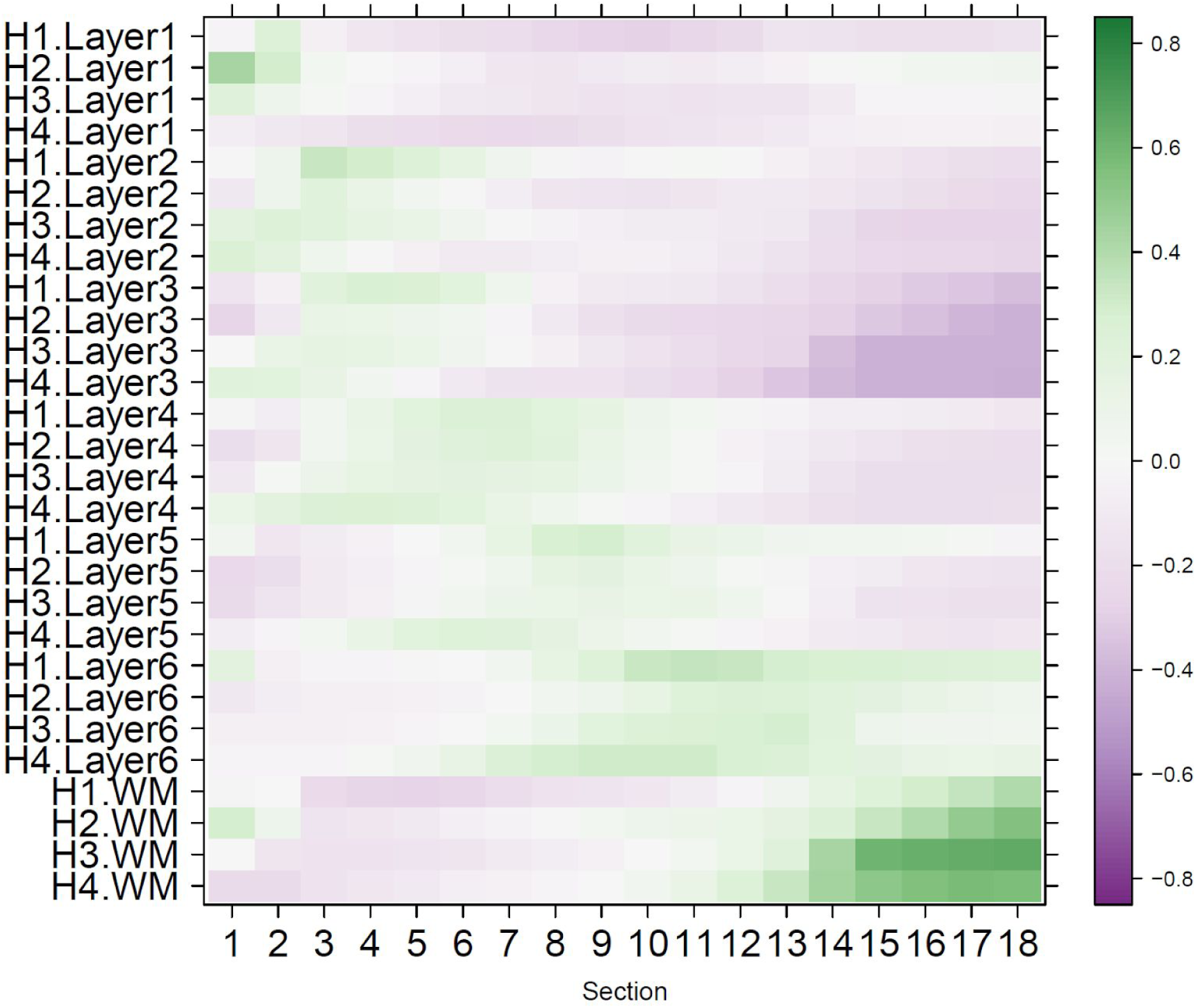
Spatial registration of bulk RNA-seq data from serial sections from He *et al*., related to. Figure 5. Heatmaps of Pearson correlation values evaluating the relationship between our Visium-derived layer-enriched statistics across 700 genes for each of the four individuals from that study (y-axis) across the 18 serial sections for each donor.

**Figure S14:**
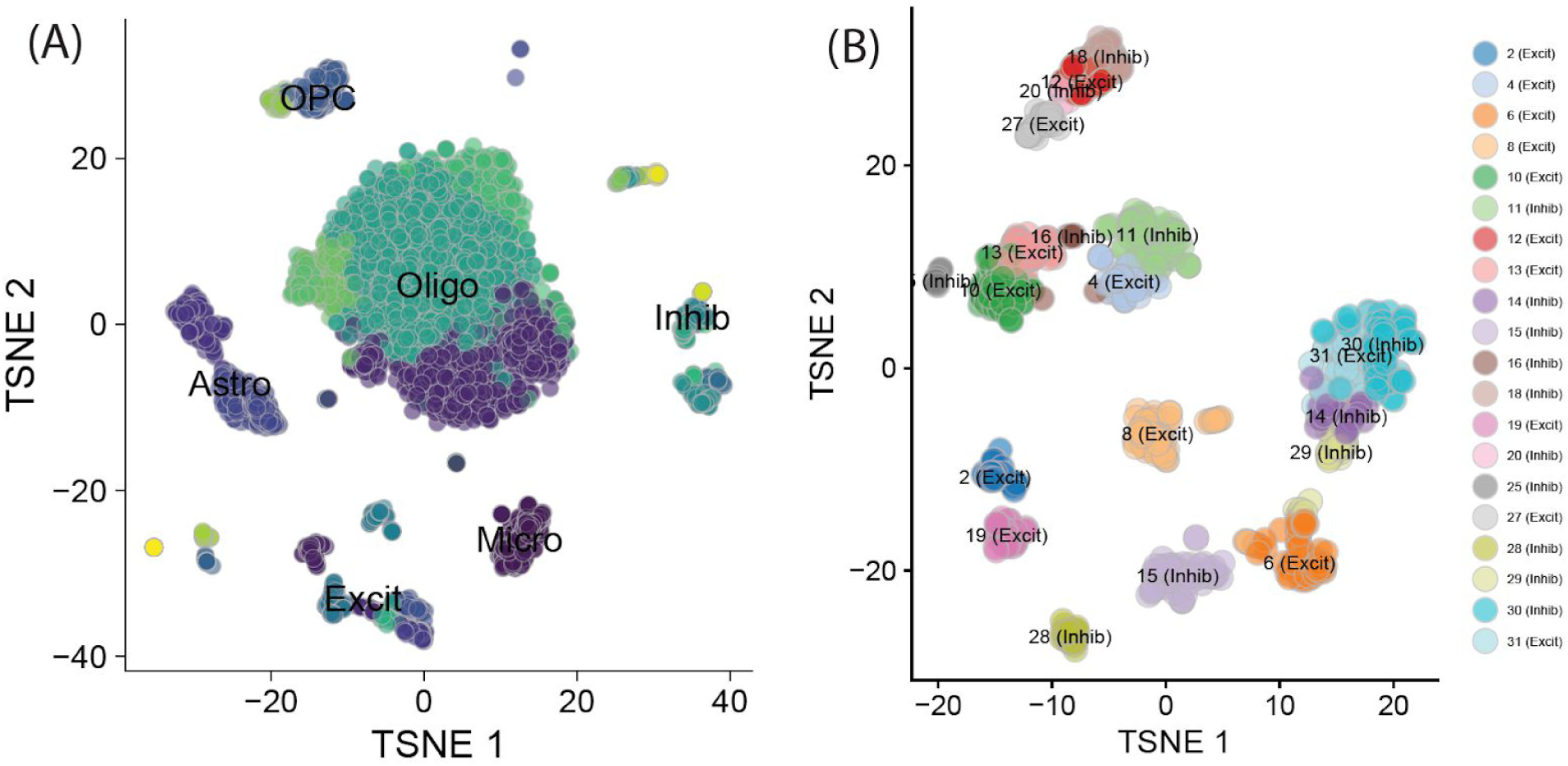
t-SNE plots of snRNA-seq data from DLPFC, related to. Figure 5. (**A**) tSNE plot of all nuclei, across 31 clusters. (**B**) tSNE plot of the subset of all neuronal nuclei.

**Figure S15.**
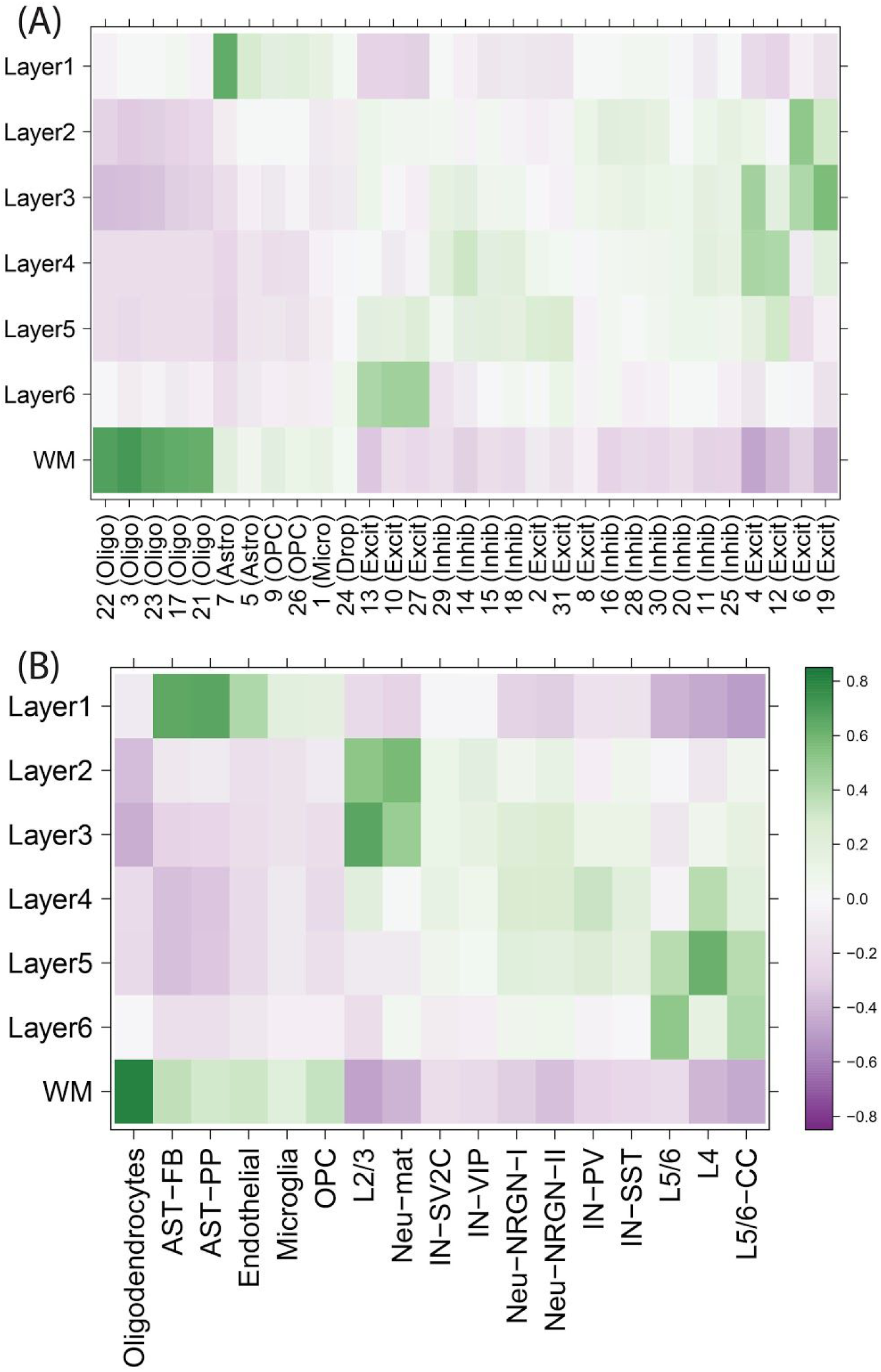
Spatial registration of snRNA-seq data, related to. Figure 5. Heatmaps of Pearson correlation values evaluating the relationship between our Visium-derived layer-enriched statistics (y-axis) for 700 genes and (**A**) Data from DLPFC from two donors, with data-driven cluster numbers and broad cell classes on the x-axis. (**B**) Data from Velmeshev *et al*. with data-driven clusters provided in their processed data.

**Figure S16.**
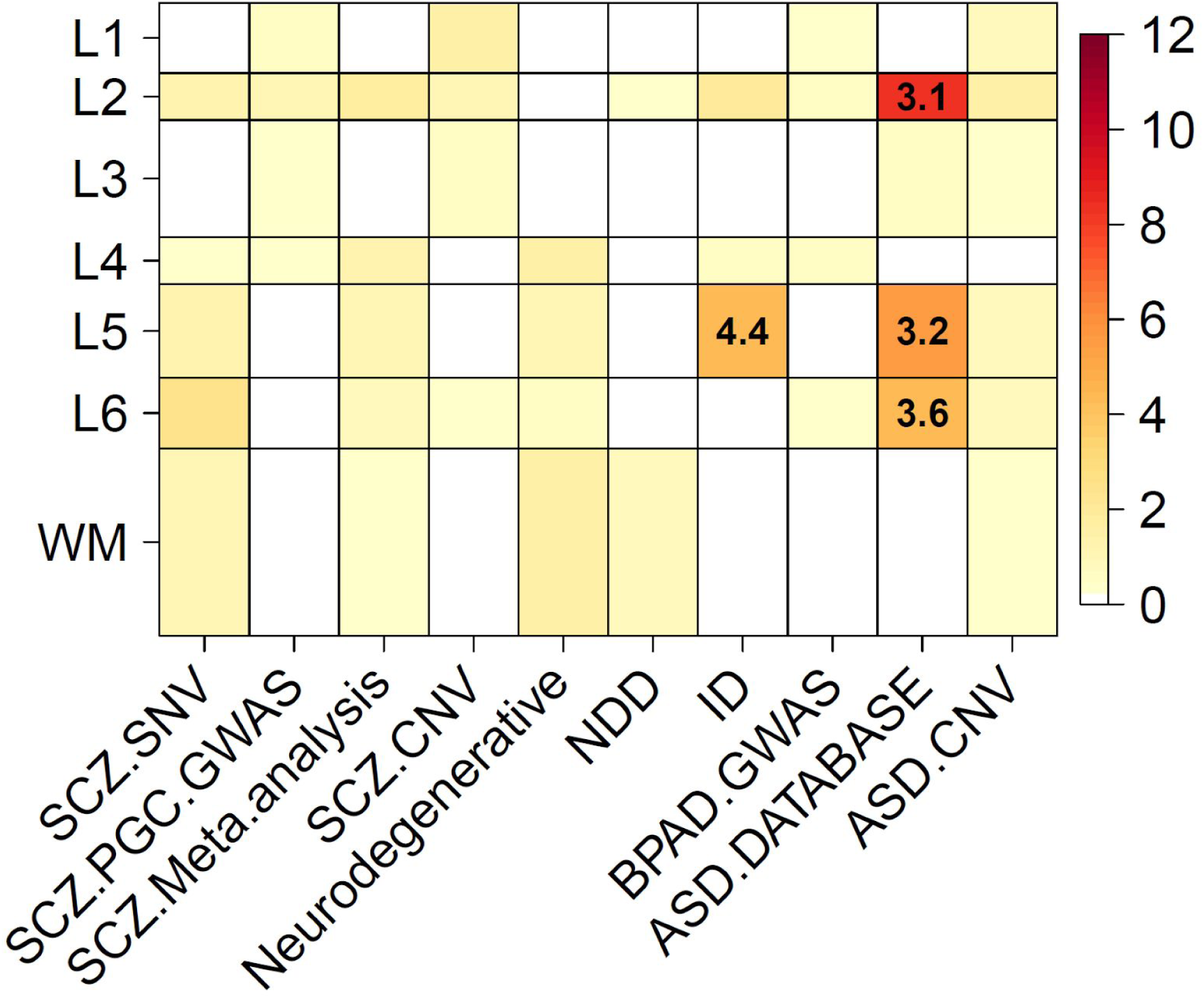
Enrichment of clinical gene sets for different neuropsychiatric and neurodevelopmental disorders, related to. Figure 6. Shown are Fisher’s exact test odds ratios and *p*-values for our Visium-derived layer-enriched statistics versus a series of predefined gene sets. Color scales indicate -log10(*p*-values), which were thresholded at *p*=10^-12^, and numbers within significant heatmap cells indicate odds ratios (ORs) for the enrichments.

**Figure S17.**
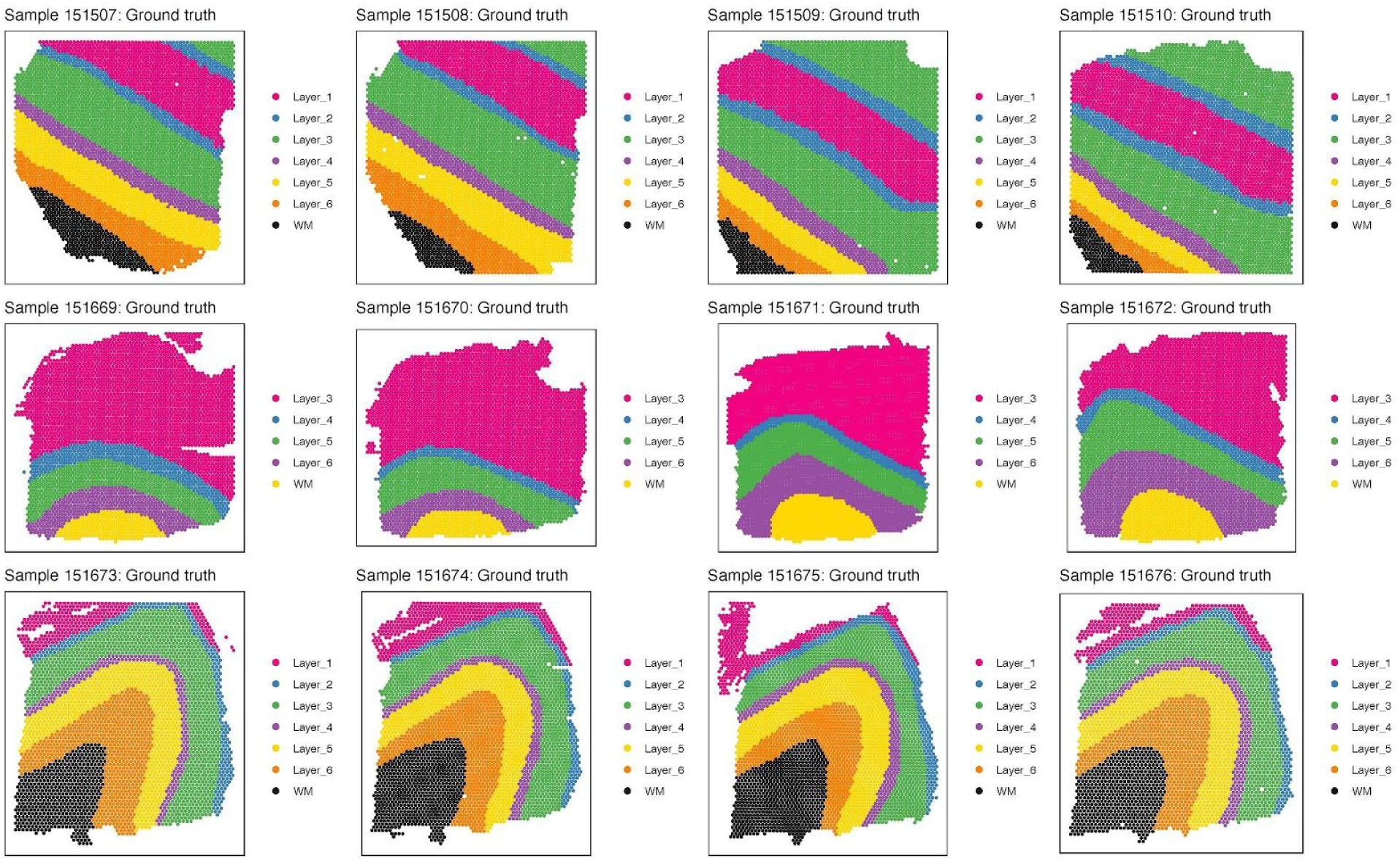
Supervised annotation of DLPFC layers across all samples, related to Figure 7. These ‘manually annotated’ layers were used as the ‘ground truth’ for evaluating the data-driven clustering results for each sample. Colors represent the six DLPFC layers and white matter (WM), and are arranged in a consistent order across samples.

**Figure S18.**
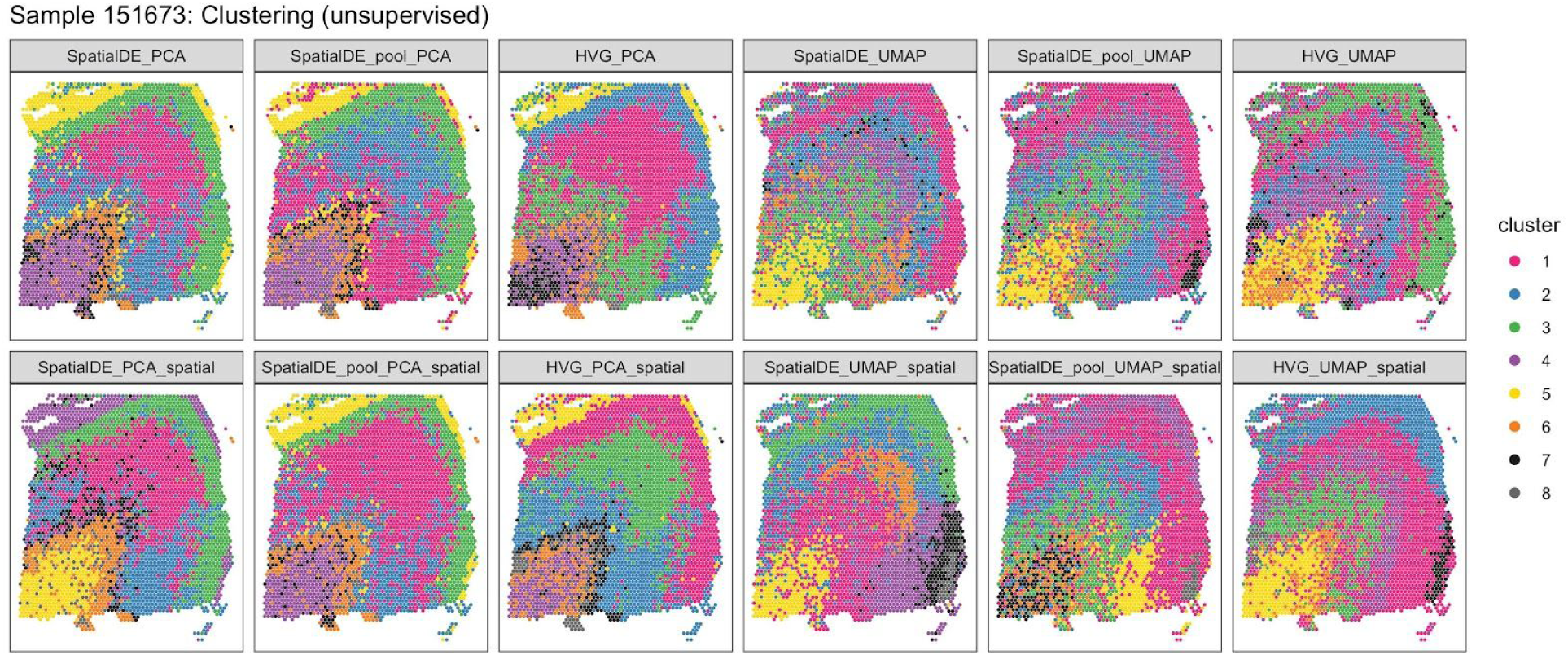
‘Unsupervised’ clustering results for sample 151673, related to. Figure 7. Visualization of clustering results for ‘unsupervised’ methods (**Table S10**) for sample 151673. Each panel displays clustering results from one clustering method. Rows display methods either without (top row) or with (bottom row) spatial coordinates included as additional features for clustering. A complete description of the different combinations of methodologies implemented in the clustering methods is provided in **Table S10**. See also **Supplementary File 1**.

**Figure S19.**
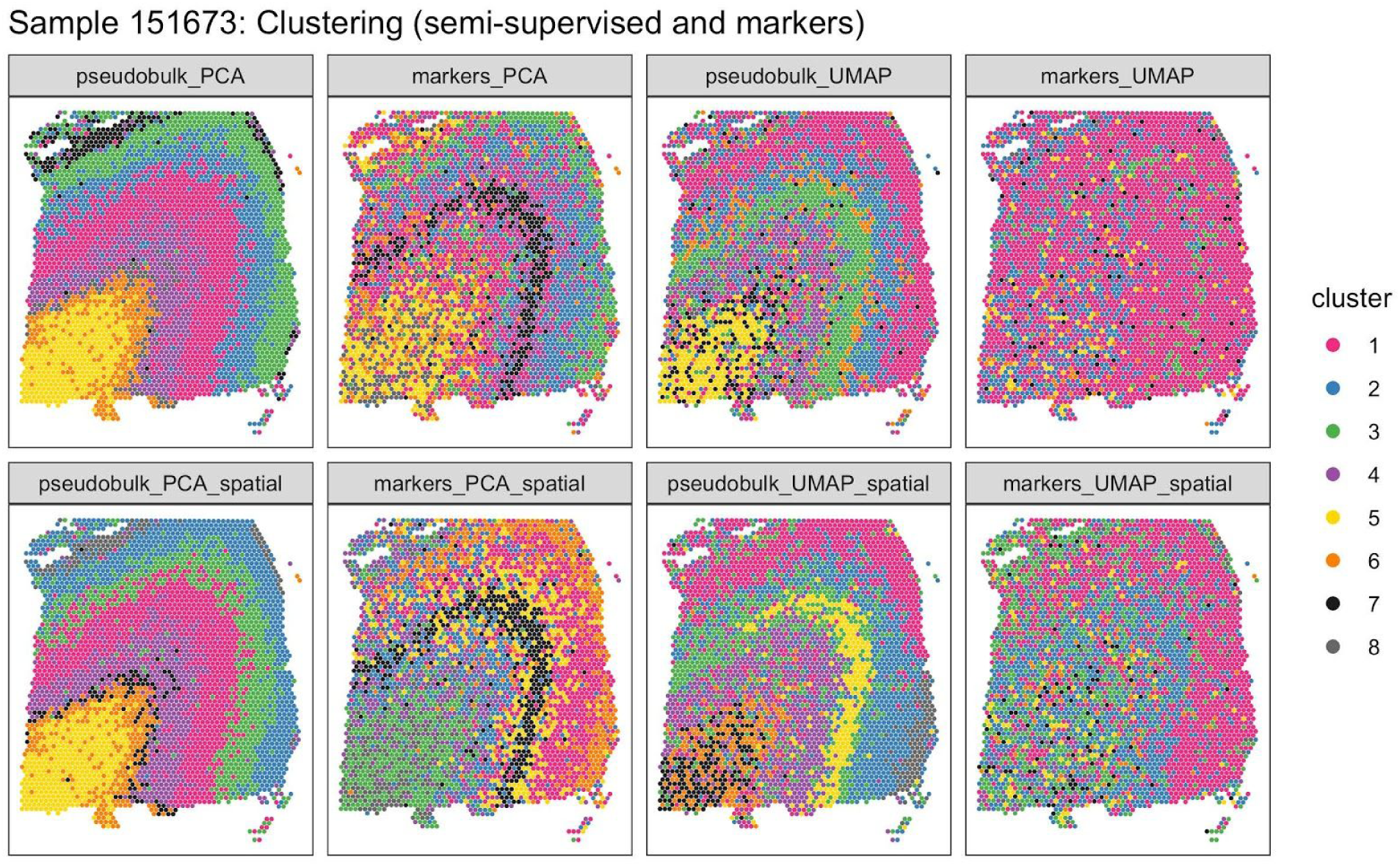
‘Semi-supervised’ and ‘markers’ clustering results for sample 151673, related to. Figure 7. Visualization of clustering results for ‘semi-supervised’ and known ‘markers’ gene set-based methods (**Table S10**) for sample 151673. Each panel displays clustering results from one clustering method. Rows display methods either without (top row) or with (bottom row) spatial coordinates included as additional features for clustering. A complete description of the different combinations of methodologies implemented in the clustering methods is provided in **Table S10**. See also **Supplementary File 1**.

**Figure S20.**
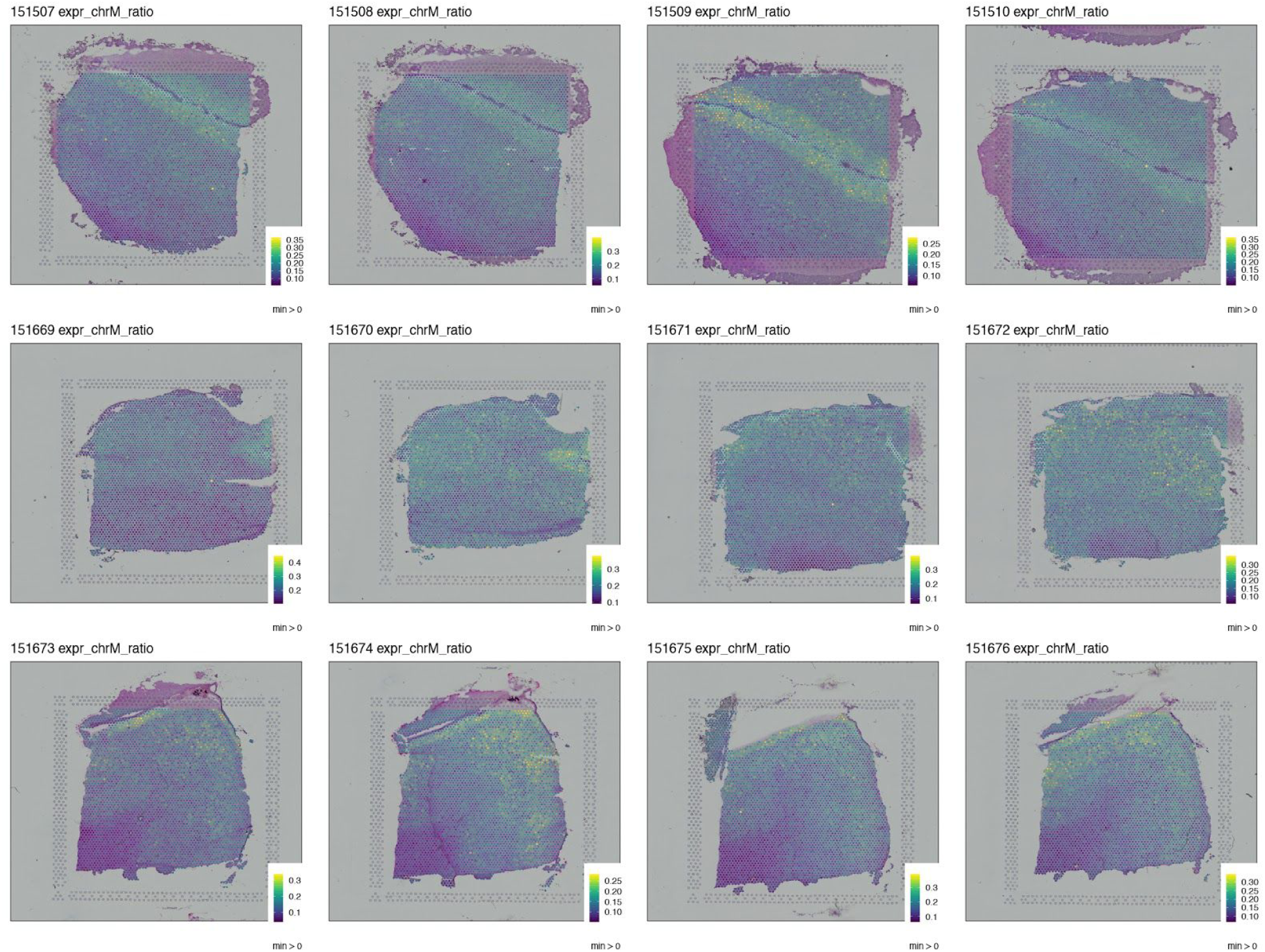
Mitochondrial proportion of expression at the spot-level, related to Discussion. Visualization of the proportion of mitochondrial gene expression compared to the total gene expression at the spot-level. Each sample has its own color scale in order for the dynamic range to be visible for each sample.

## Bibliography

1. Aaron Lun [Aut, C., Davide Risso (2017). SingleCellExperiment. Bioconductor.

2. Abrahams, B.S., Arking, D.E., Campbell, D.B., Mefford, H.C., Morrow, E.M., Weiss, L.A., Menashe, I., Wadkins, T., Banerjee-Basu, S., and Packer, A. (2013). SFARI Gene 2.0: a community-driven knowledgebase for the autism spectrum disorders (ASDs). Mol. Autism 4, 36.

3. Amezquita, R.A., Lun, A.T.L., Becht, E., Carey, V.J., Carpp, L.N., Geistlinger, L., Marini, F., Rue-Albrecht, K., Risso, D., Soneson, C., et al. (2020). Orchestrating single-cell analysis with Bioconductor. Nat. Methods 17, 137–145.

4. Asp, M., Giacomello, S., Fürth, D., Reimegård, J., Wärdell, E., Custodio, J., Salmén, F., Sundström, E., Åkesson, E., Bienko, M., et al. (2018). An Organ-Wide Gene Expression Atlas of the Developing Human Heart. SSRN Journal.

5. Bakken, T.E., Hodge, R.D., Miller, J.A., Yao, Z., Nguyen, T.N., Aevermann, B., Barkan, E., Bertagnolli, D., Casper, T., Dee, N., et al. (2018). Single-nucleus and single-cell transcriptomes compared in matched cortical cell types. PLoS ONE 13, e0209648.

6. Berglund, E., Maaskola, J., Schultz, N., Friedrich, S., Marklund, M., Bergenstråhle, J., Tarish, F., Tanoglidi, A., Vickovic, S., Larsson, L., et al. (2018). Spatial maps of prostate cancer transcriptomes reveal an unexplored landscape of heterogeneity. Nat. Commun. 9, 2419.

7. Biever, A., Donlin-Asp, P.G., and Schuman, E.M. (2019). Local translation in neuronal processes. Curr. Opin. Neurobiol. 57, 141–148.

8. Bioconductor Package Maintainer (2017). ExperimentHub. Bioconductor.

9. Birnbaum, R., Jaffe, A.E., Hyde, T.M., Kleinman, J.E., and Weinberger, D.R. (2014). Prenatal expression patterns of genes associated with neuropsychiatric disorders. Am. J. Psychiatry 171, 758–767.

10. Bulik-Sullivan, B., Loh, P.-R., Finucane, H.K., Ripke, S., Yang, J., Schizophrenia Working Group of the Psychiatric Genomics Consortium, Patterson, N., Daly, M.J., Price, A.L., and Neale, B.M. (2015). LD Score regression distinguishes confounding from polygenicity in genome-wide association studies. Nat. Genet. 47, 291–295.

11. Burgess, D.J. (2019). Spatial transcriptomics coming of age. Nat. Rev. Genet. 20, 317.

12. Chang, W., Cheng, J., Allaire, J.J., Xie, Y., and McPherson, J. (2019). shiny: Web Application Framework for R.

13. Chen, K.H., Boettiger, A.N., Moffitt, J.R., Wang, S., and Zhuang, X. (2015). RNA imaging. Spatially resolved, highly multiplexed RNA profiling in single cells. Science 348, aaa6090.

14. Codeluppi, S., Borm, L.E., Zeisel, A., La Manno, G., van Lunteren, J.A., Svensson, C.I., and Linnarsson, S. (2018). Spatial organization of the somatosensory cortex revealed by osmFISH. Nat. Methods 15, 932–935.

15. Collado-Torres, L. (2020). LieberInstitute/spatialLIBD: spatialLIBD: initial Bioconductor submission. Zenodo.

16. Collado-Torres, L., Burke, E.E., Peterson, A., Shin, J., Straub, R.E., Rajpurohit, A., Semick, S.A., Ulrich, W.S., BrainSeq Consortium, Price, A.J., et al. (2019). Regional Heterogeneity in Gene Expression, Regulation, and Coherence in the Frontal Cortex and Hippocampus across Development and Schizophrenia. Neuron 103, 203–216.e8.

17. Collado-Torres, L., Weber, L., and Hicks, S. (2020). LieberInstitute/HumanPilot: Archive the HumanPilot code for the preprint version of our project. Zenodo.

18. Crowell, H.L., Soneson, C., Germain, P.-L., Calini, D., Collin, L., Raposo, C., Malhotra, D., and Robinson, M.D. (2019). On the discovery of population-specific state transitions from multi-sample multi-condition single-cell RNA sequencing data. BioRxiv.

19. Csardi, G., and Nepusz, T. (2006). The igraph software package for complex network research. InterJournal Complex Systems, 1695.

20. Darmanis, S., Sloan, S.A., Zhang, Y., Enge, M., Caneda, C., Shuer, L.M., Hayden Gephart, M.G., Barres, B.A., and Quake, S.R. (2015). A survey of human brain transcriptome diversity at the single cell level. Proc Natl Acad Sci USA 112, 7285–7290.

21. DeFelipe, J., and Fariñas, I. (1992). The pyramidal neuron of the cerebral cortex: morphological and chemical characteristics of the synaptic inputs. Prog. Neurobiol. 39, 563–607.

22. Dobin, A., Davis, C.A., Schlesinger, F., Drenkow, J., Zaleski, C., Jha, S., Batut, P., Chaisson, M., and Gingeras, T.R. (2013). STAR: ultrafast universal RNA-seq aligner. Bioinformatics 29, 15–21.

23. Dong, X., Shi, M., Lee, M., Toro, R., Gravina, S., Han, W., Yasuda, S., Wang, T., Zhang, Z., Vijg, J., et al. (2018). Global, integrated analysis of methylomes and transcriptomes from laser capture microdissected bronchial and alveolar cells in human lung. Epigenetics 13, 264–274.

24. Finucane, H.K., Bulik-Sullivan, B., Gusev, A., Trynka, G., Reshef, Y., Loh, P.-R., Anttila, V., Xu, H., Zang, C., Farh, K., et al. (2015). Partitioning heritability by functional annotation using genome-wide association summary statistics. Nat. Genet. 47, 1228–1235.

25. Fromer, M., Roussos, P., Sieberts, S.K., Johnson, J.S., Kavanagh, D.H., Perumal, T.M., Ruderfer, D.M., Oh, E.C., Topol, A., Shah, H.R., et al. (2016). Gene expression elucidates functional impact of polygenic risk for schizophrenia. Nat. Neurosci. 19, 1442–1453.

26. Gandal, M.J., Zhang, P., Hadjimichael, E., Walker, R.L., Chen, C., Liu, S., Won, H., van Bakel, H., Varghese, M., Wang, Y., et al. (2018). Transcriptome-wide isoform-level dysregulation in ASD, schizophrenia, and bipolar disorder. Science 362.

27. Gregory, J.M., McDade, K., Livesey, M.R., Croy, I., Marion de Proce, S., Aitman, T., Chandran, S., and Smith, C. (2020). Spatial transcriptomics identifies spatially dysregulated expression of GRM3 and USP47 in amyotrophic lateral sclerosis. Neuropathol. Appl. Neurobiol.

28. Griffiths, J.A., Richard, A.C., Bach, K., Lun, A.T.L., and Marioni, J.C. (2018). Detection and removal of barcode swapping in single-cell RNA-seq data. Nat. Commun. 9, 2667.

29. Grove, J., Ripke, S., Als, T.D., Mattheisen, M., Walters, R.K., Won, H., Pallesen, J., Agerbo, E., Andreassen, O.A., Anney, R., et al. (2019). Identification of common genetic risk variants for autism spectrum disorder. Nat. Genet. 51, 431–444.

30. Gusev, A., Ko, A., Shi, H., Bhatia, G., Chung, W., Penninx, B.W.J.H., Jansen, R., de Geus, E.J.C., Boomsma, D.I., Wright, F.A., et al. (2016). Integrative approaches for large-scale transcriptome-wide association studies. Nat. Genet. 48, 245–252.

31. Habib, N., Li, Y., Heidenreich, M., Swiech, L., Avraham-Davidi, I., Trombetta, J.J., Hession, C., Zhang, F., and Regev, A. (2016). Div-Seq: Single-nucleus RNA-Seq reveals dynamics of rare adult newborn neurons. Science 353, 925–928.

32. Habib, N., Avraham-Davidi, I., Basu, A., Burks, T., Shekhar, K., Hofree, M., Choudhury, S.R., Aguet, F., Gelfand, E., Ardlie, K., et al. (2017). Massively parallel single-nucleus RNA-seq with DroNc-seq. Nat. Methods 14, 955–958.

33. Hafner, A.-S., Donlin-Asp, P.G., Leitch, B., Herzog, E., and Schuman, E.M. (2019). Local protein synthesis is a ubiquitous feature of neuronal pre- and postsynaptic compartments. Science 364.

34. Harris, K.D., and Shepherd, G.M.G. (2015). The neocortical circuit: themes and variations. Nat. Neurosci. 18, 170–181.

35. Harris, J.J., Jolivet, R., and Attwell, D. (2012). Synaptic energy use and supply. Neuron 75, 762–777.

36. Hawrylycz, M.J., Lein, E.S., Guillozet-Bongaarts, A.L., Shen, E.H., Ng, L., Miller, J.A., van de Lagemaat, L.N., Smith, K.A., Ebbert, A., Riley, Z.L., et al. (2012). An anatomically comprehensive atlas of the adult human brain transcriptome. Nature 489, 391–399.

37. He, Z., Han, D., Efimova, O., Guijarro, P., Yu, Q., Oleksiak, A., Jiang, S., Anokhin, K., Velichkovsky, B., Grünewald, S., et al. (2017). Comprehensive transcriptome analysis of neocortical layers in humans, chimpanzees and macaques. Nat. Neurosci. 20, 886–895.

38. Hodge, R.D., Bakken, T.E., Miller, J.A., Smith, K.A., Barkan, E.R., Graybuck, L.T., Close, J.L., Long, B., Johansen, N., Penn, O., et al. (2019). Conserved cell types with divergent features in human versus mouse cortex. Nature 573, 61–68.

39. Hu, P., Fabyanic, E., Kwon, D.Y., Tang, S., Zhou, Z., and Wu, H. (2017). Dissecting Cell-Type Composition and Activity-Dependent Transcriptional State in Mammalian Brains by Massively Parallel Single-Nucleus RNA-Seq. Mol. Cell 68, 1006–1015.e7.

40. Jaffe, A.E., Straub, R.E., Shin, J.H., Tao, R., Gao, Y., Collado-Torres, L., Kam-Thong, T., Xi, H.S., Quan, J., Chen, Q., et al. (2018). Developmental and genetic regulation of the human cortex transcriptome illuminate schizophrenia pathogenesis. Nat. Neurosci. 21, 1117–1125.

41. Jaffe, A.E., Hoeppner, D.J., Saito, T., Blanpain, L., Ukaigwe, J., Burke, E.E., Tao, R., Tajinda, K., Deep-Soboslay, A., Shin, J.H., et al. (2019). Cell type-specific genetic regulation of expression in the granule cell layer of the human dentate gyrus. BioRxiv.

42. Jaffe, A.E., Hoeppner, D.J., Saito, T., Blanpain, L., Ukaigwe, J., Burke, E.E., Tao, R., Tajinda, K., Deep-Soboslay, A., Shin, J.H., et al. (2020). Genetic regulation of expression in the granule cell layer of the human dentate gyrus. Nat Neurosci.

43. Kang, H.M., Subramaniam, M., Targ, S., Nguyen, M., Maliskova, L., McCarthy, E., Wan, E., Wong, S., Byrnes, L., Lanata, C.M., et al. (2018). Multiplexed droplet single-cell RNA-sequencing using natural genetic variation. Nat. Biotechnol. 36, 89–94.

44. Lacar, B., Linker, S.B., Jaeger, B.N., Krishnaswami, S.R., Barron, J.J., Kelder, M.J.E., Parylak, S.L., Paquola, A.C.M., Venepally, P., Novotny, M., et al. (2016). Nuclear RNA-seq of single neurons reveals molecular signatures of activation. Nat. Commun. 7, 11022.

45. Lake, B.B., Ai, R., Kaeser, G.E., Salathia, N.S., Yung, Y.C., Liu, R., Wildberg, A., Gao, D., Fung, H.-L., Chen, S., et al. (2016). Neuronal subtypes and diversity revealed by single-nucleus RNA sequencing of the human brain. Science 352, 1586–1590.

46. Lake, B.B., Chen, S., Sos, B.C., Fan, J., Kaeser, G.E., Yung, Y.C., Duong, T.E., Gao, D., Chun, J., Kharchenko, P.V., et al. (2018). Integrative single-cell analysis of transcriptional and epigenetic states in the human adult brain. Nat. Biotechnol. 36, 70–80.

47. Langfelder, P., Zhang, B., and with contributions from Steve Horvath (2016). dynamicTreeCut: Methods for Detection of Clusters in Hierarchical Clustering Dendrograms.

48. Law, C.W., Chen, Y., Shi, W., and Smyth, G.K. (2014). voom: Precision weights unlock linear model analysis tools for RNA-seq read counts. Genome Biol. 15, R29.

49. de Leeuw, C.A., Mooij, J.M., Heskes, T., and Posthuma, D. (2015). MAGMA: generalized gene-set analysis of GWAS data. PLoS Comput. Biol. 11, e1004219.

50. Lein, E., Borm, L.E., and Linnarsson, S. (2017). The promise of spatial transcriptomics for neuroscience in the era of molecular cell typing. Science 358, 64–69.

51. Lipska, B.K., Deep-Soboslay, A., Weickert, C.S., Hyde, T.M., Martin, C.E., Herman, M.M., and Kleinman, J.E. (2006). Critical factors in gene expression in postmortem human brain: Focus on studies in schizophrenia. Biol. Psychiatry 60, 650–658.

52. Lun, A. (2019). BiocSingular. Bioconductor.

53. Lun, A.T.L., and Marioni, J.C. (2017). Overcoming confounding plate effects in differential expression analyses of single-cell RNA-seq data. Biostatistics 18, 451–464.

54. Lun, A.T.L., McCarthy, D.J., and Marioni, J.C. (2016). A step-by-step workflow for low-level analysis of single-cell RNA-seq data with Bioconductor. [version 2; peer review: 3 approved, 2 approved with reservations]. F1000Res. 5, 2122.

55. Lun, A.T.L., Riesenfeld, S., Andrews, T., Dao, T.P., Gomes, T., participants in the 1st Human Cell Atlas Jamboree, and Marioni, J.C. (2019). EmptyDrops: distinguishing cells from empty droplets in droplet-based single-cell RNA sequencing data. Genome Biol. 20, 63.

56. van der Maaten, L., and Hinton, G. (2008). Visualizing Data using t-SNE. Journal of Machine Learning Research 9, 2579–2605.

57. Maniatis, S., Äijö, T., Vickovic, S., Braine, C., Kang, K., Mollbrink, A., Fagegaltier, D., Andrusivová, Ž., Saarenpää, S., Saiz-Castro, G., et al. (2019). Spatiotemporal dynamics of molecular pathology in amyotrophic lateral sclerosis. Science 364, 89–93.

58. Martin Morgan, V.O. (2017). SummarizedExperiment. Bioconductor.

59. Mathys, H., Davila-Velderrain, J., Peng, Z., Gao, F., Mohammadi, S., Young, J.Z., Menon, M., He, L., Abdurrob, F., Jiang, X., et al. (2019). Single-cell transcriptomic analysis of Alzheimer’s disease. Nature 570, 332–337.

60. Maynard, K.R., Tippani, M., Takahashi, Y., Phan, B.N., Hyde, T.M., Jaffe, A.E., and Martinowich, K. (2019). dotdotdot: an automated approach to quantify multiplex single molecule fluorescent in situ hybridization (smFISH) images in complex tissues. BioRxiv.

61. McCarthy, D.J., Campbell, K.R., Lun, A.T.L., and Wills, Q.F. (2017). Scater: pre-processing, quality control, normalization and visualization of single-cell RNA-seq data in R. Bioinformatics 33, 1179–1186.

62. McInnes, L., Healy, J., Saul, N., and Großberger, L. (2018). UMAP: uniform manifold approximation and projection. JOSS 3, 861.

63. Melville, J. (2019). uwot: R package.

64. Molyneaux, B.J., Arlotta, P., Menezes, J.R.L., and Macklis, J.D. (2007). Neuronal subtype specification in the cerebral cortex. Nat. Rev. Neurosci. 8, 427–437.

65. Moncada, R., Chiodin, M., Devlin, J.C., Baron, M., Hajdu, C.H., Simeone, D., and Yanai, I. (2018). Building a tumor atlas: integrating single-cell RNA-Seq data with spatial transcriptomics in pancreatic ductal adenocarcinoma. BioRxiv [Preprint]. accessed 1.12.2019.

66. Moyer, C.E., Shelton, M.A., and Sweet, R.A. (2015). Dendritic spine alterations in schizophrenia. Neurosci. Lett. 601, 46–53.

67. Narayanan, R.T., Udvary, D., and Oberlaender, M. (2017). Cell Type-Specific Structural Organization of the Six Layers in Rat Barrel Cortex. Front. Neuroanat. 11, 91.

68. Nowakowski, T.J., Bhaduri, A., Pollen, A.A., Alvarado, B., Mostajo-Radji, M.A., Di Lullo, E., Haeussler, M., Sandoval-Espinosa, C., Liu, S.J., Velmeshev, D., et al. (2017). Spatiotemporal gene expression trajectories reveal developmental hierarchies of the human cortex. Science 358, 1318–1323.

69. Olmos-Serrano, J.L., Kang, H.J., Tyler, W.A., Silbereis, J.C., Cheng, F., Zhu, Y., Pletikos, M., Jankovic-Rapan, L., Cramer, N.P., Galdzicki, Z., et al. (2016). Down syndrome developmental brain transcriptome reveals defective oligodendrocyte differentiation and myelination. Neuron 89, 1208–1222.

70. Overly, C.C., Rieff, H.I., and Hollenbeck, P.J. (1996). Organelle motility and metabolism in axons vs dendrites of cultured hippocampal neurons. J. Cell Sci. 109 ( *Pt 5*), 971–980.

71. Pardiñas, A.F., Holmans, P., Pocklington, A.J., Escott-Price, V., Ripke, S., Carrera, N., Legge, S.E., Bishop, S., Cameron, D., Hamshere, M.L., et al. (2018). Common schizophrenia alleles are enriched in mutation-intolerant genes and in regions under strong background selection. Nat. Genet. 50, 381–389.

72. Radnikow, G., and Feldmeyer, D. (2018). Layer- and Cell Type-Specific Modulation of Excitatory Neuronal Activity in the Neocortex. Front. Neuroanat. 12, 1.

73. Rajkowska, G., and Goldman-Rakic, P.S. (1995a). Cytoarchitectonic definition of prefrontal areas in the normal human cortex: I. Remapping of areas 9 and 46 using quantitative criteria. Cereb. Cortex 5, 307–322.

74. Rajkowska, G., and Goldman-Rakic, P.S. (1995b). Cytoarchitectonic definition of prefrontal areas in the normal human cortex: II. Variability in locations of areas 9 and 46 and relationship to the Talairach Coordinate System. Cereb. Cortex 5, 323–337.

75. Ritchie, M.E., Phipson, B., Wu, D., Hu, Y., Law, C.W., Shi, W., and Smyth, G.K. (2015). limma powers differential expression analyses for RNA-sequencing and microarray studies. Nucleic Acids Res. 43, e47.

76. Rizzardi, L.F., Hickey, P.F., Rodriguez DiBlasi, V., Tryggvadóttir, R., Callahan, C.M., Idrizi, A., Hansen, K.D., and Feinberg, A.P. (2019). Neuronal brain-region-specific DNA methylation and chromatin accessibility are associated with neuropsychiatric trait heritability. Nat. Neurosci. 22, 307–316.

77. Rodriques, S.G., Stickels, R.R., Goeva, A., Martin, C.A., Murray, E., Vanderburg, C.R., Welch, J., Chen, L.M., Chen, F., and Macosko, E.Z. (2019). Slide-seq: A scalable technology for measuring genome-wide expression at high spatial resolution. Science 363, 1463–1467.

78. Satterstrom, F.K., Kosmicki, J.A., Wang, J., Breen, M.S., De Rubeis, S., An, J.-Y., Peng, M., Collins, R., Grove, J., Klei, L., et al. (2020). Large-Scale Exome Sequencing Study Implicates Both Developmental and Functional Changes in the Neurobiology of Autism. Cell 180, 568–584.e23.

79. Sievert, C. (2018). plotly for R.

80. Skene, N.G., Bryois, J., Bakken, T.E., Breen, G., Crowley, J.J., Gaspar, H.A., Giusti-Rodriguez, P., Hodge, R.D., Miller, J.A., Muñoz-Manchado, A.B., et al. (2018). Genetic identification of brain cell types underlying schizophrenia. Nat. Genet. 50, 825–833.

81. Stahl, E.A., Breen, G., Forstner, A.J., McQuillin, A., Ripke, S., Trubetskoy, V., Mattheisen, M., Wang, Y., Coleman, J.R.I., Gaspar, H.A., et al. (2019). Genome-wide association study identifies 30 loci associated with bipolar disorder. Nat. Genet. 51, 793–803.

82. Ståhl, P.L., Salmén, F., Vickovic, S., Lundmark, A., Navarro, J.F., Magnusson, J., Giacomello, S., Asp, M., Westholm, J.O., Huss, M., et al. (2016). Visualization and analysis of gene expression in tissue sections by spatial transcriptomics. Science 353, 78–82.

83. Svensson, V., Teichmann, S.A., and Stegle, O. (2018). SpatialDE: identification of spatially variable genes. Nat. Methods 15, 343–346.

84. Sweet, R.A., Fish, K.N., and Lewis, D.A. (2010). Mapping Synaptic Pathology within Cerebral Cortical Circuits in Subjects with Schizophrenia. Front. Hum. Neurosci. 4, 44.

85. Velmeshev, D., Schirmer, L., Jung, D., Haeussler, M., Perez, Y., Mayer, S., Bhaduri, A., Goyal, N., Rowitch, D.H., and Kriegstein, A.R. (2019). Single-cell genomics identifies cell type-specific molecular changes in autism. Science 364, 685–689.

86. Vickovic, S., Eraslan, G., Salmén, F., Klughammer, J., Stenbeck, L., Schapiro, D., Äijö, T., Bonneau, R., Bergenstråhle, L., Navarro, J.F., et al. (2019). High-definition spatial transcriptomics for in situ tissue profiling. Nat. Methods 16, 987–990.

87. Wickham, H. (2016). ggplot2 - Elegant Graphics for Data Analysis (Cham: Springer International Publishing).

88. Wray, N.R., Ripke, S., Mattheisen, M., Trzaskowski, M., Byrne, E.M., Abdellaoui, A., Adams, M.J., Agerbo, E., Air, T.M., Andlauer, T.M.F., et al. (2018). Genome-wide association analyses identify 44 risk variants and refine the genetic architecture of major depression. Nat. Genet. 50, 668–681.

89. Zeng, H., Shen, E.H., Hohmann, J.G., Oh, S.W., Bernard, A., Royall, J.J., Glattfelder, K.J., Sunkin, S.M., Morris, J.A., Guillozet-Bongaarts, A.L., et al. (2012). Large-scale cellular-resolution gene profiling in human neocortex reveals species-specific molecular signatures. Cell 149, 483–496.

