## Supplementary File 1 for "Transcriptome-scale spatial gene expression in the human dorsolateral prefrontal cortex"

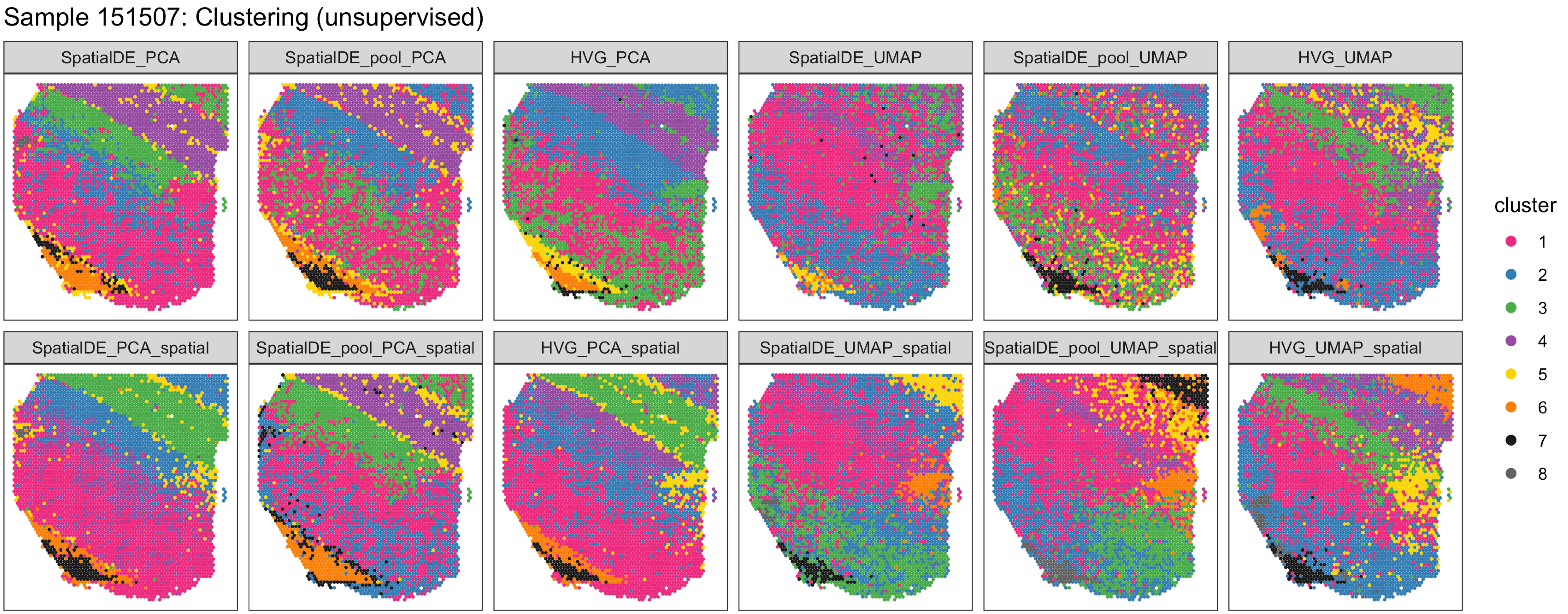

Sample 151507: Clustering (unsupervised)
SpatialDE_PCA
SpatialDE_pool_PCA
HVG_PCA
SpatialDE_UMAP
SpatialDE_pool_UMAP
HVG_UMAP
SpatialDE_PCA_spatial
SpatialDE_pool_PCA_spatial
HVG_PCA_spatial
SpatialDE_UMAP_spatial
SpatialDE_pool_UMAP_spatial
HVG_UMAP_spatial
cluster
1
2
3
4
5
6
7
8

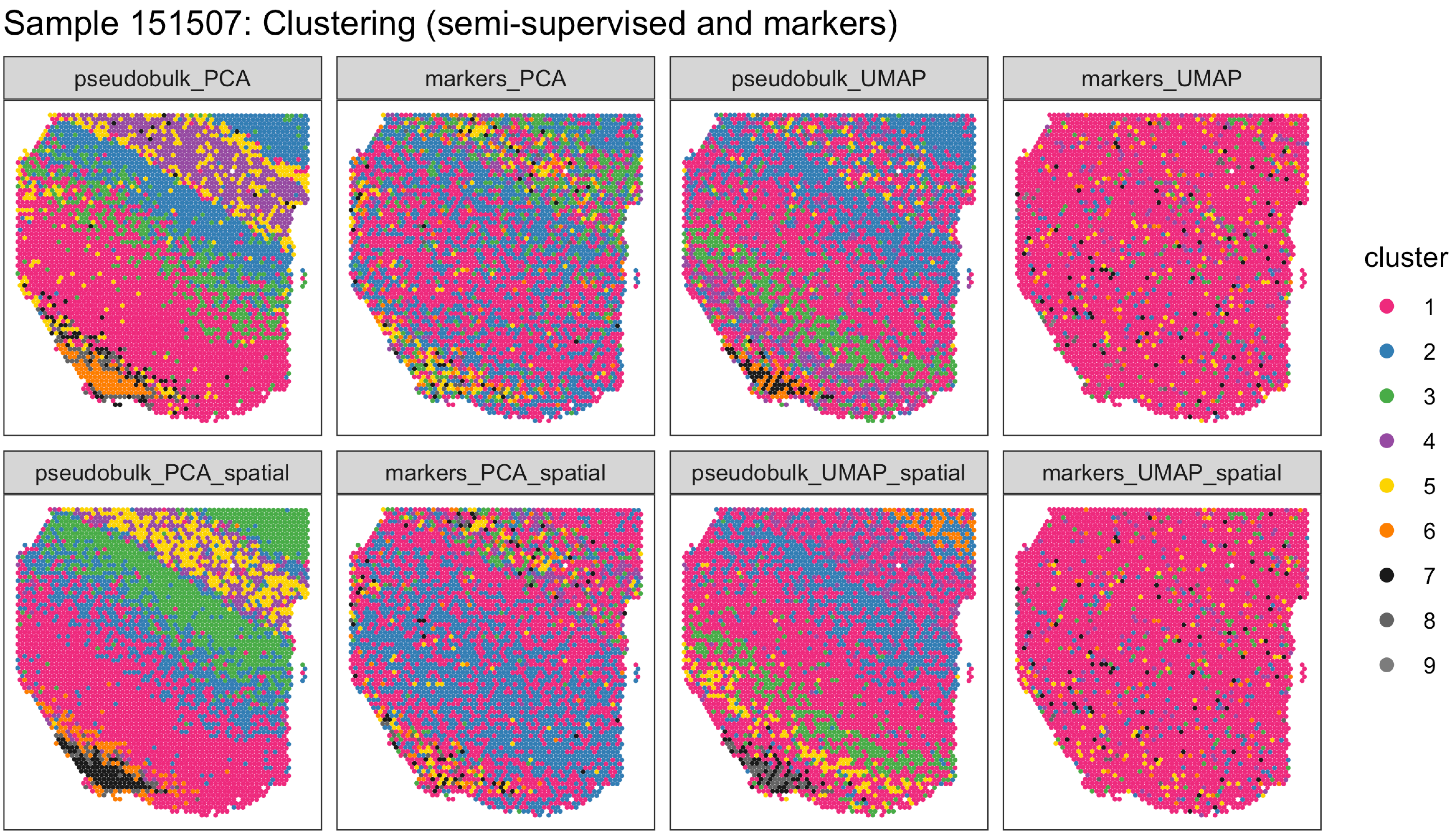
Sample 151507: Clustering (semi-supervised and markers)
pseudobulk_PCA
markers_PCA
pseudobulk_UMAP
markers_UMAP
pseudobulk_PCA_spatial
markers_PCA_spatial
pseudobulk_UMAP_spatial
markers_UMAP_spatial
cluster
1
2
3
4
5
6
7
8
9

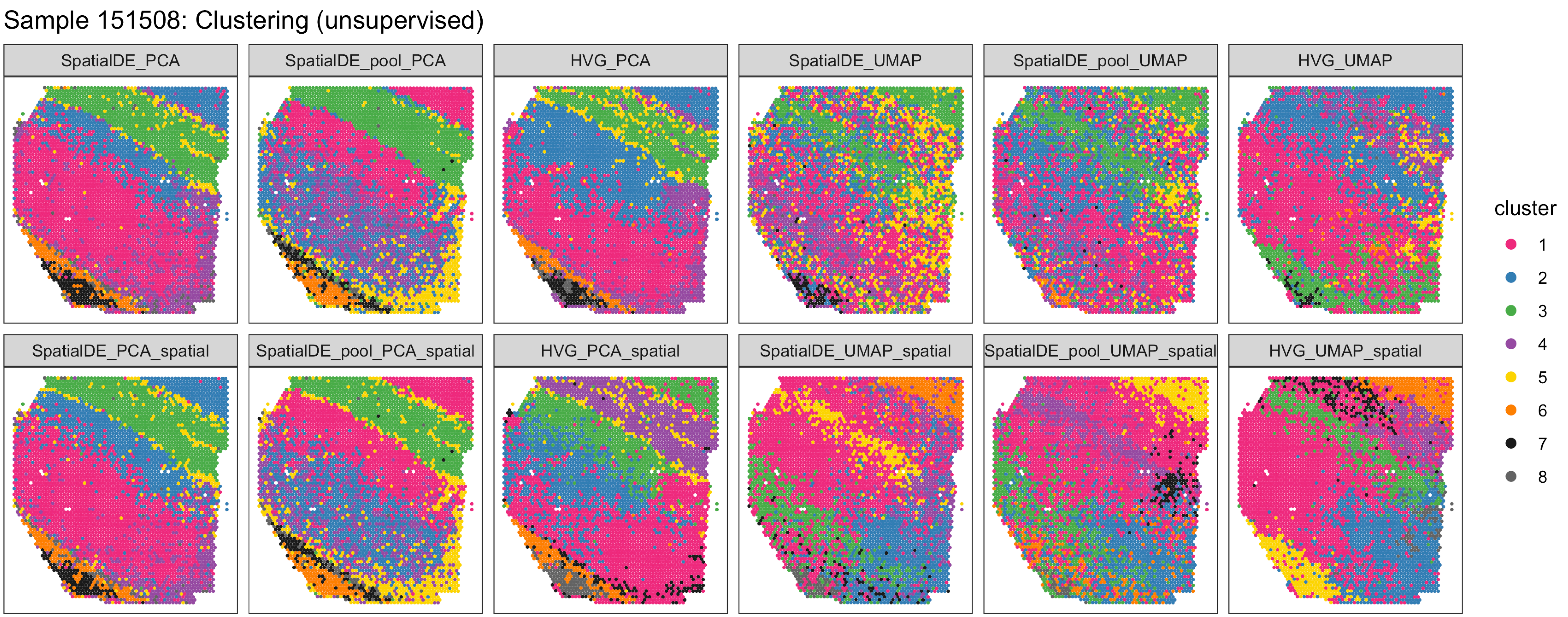
Sample 151508: Clustering (unsupervised)
SpatialDE_PCA
SpatialDE_pool_PCA
HVG_PCA
SpatialDE_UMAP
SpatialDE_pool_UMAP
HVG_UMAP
SpatialDE_PCA_spatial
SpatialDE_pool_PCA_spatial
HVG_PCA_spatial
SpatialDE_UMAP_spatial
SpatialDE_pool_UMAP_spatial
HVG_UMAP_spatial
cluster
1
2
3
4
5
6
7
8

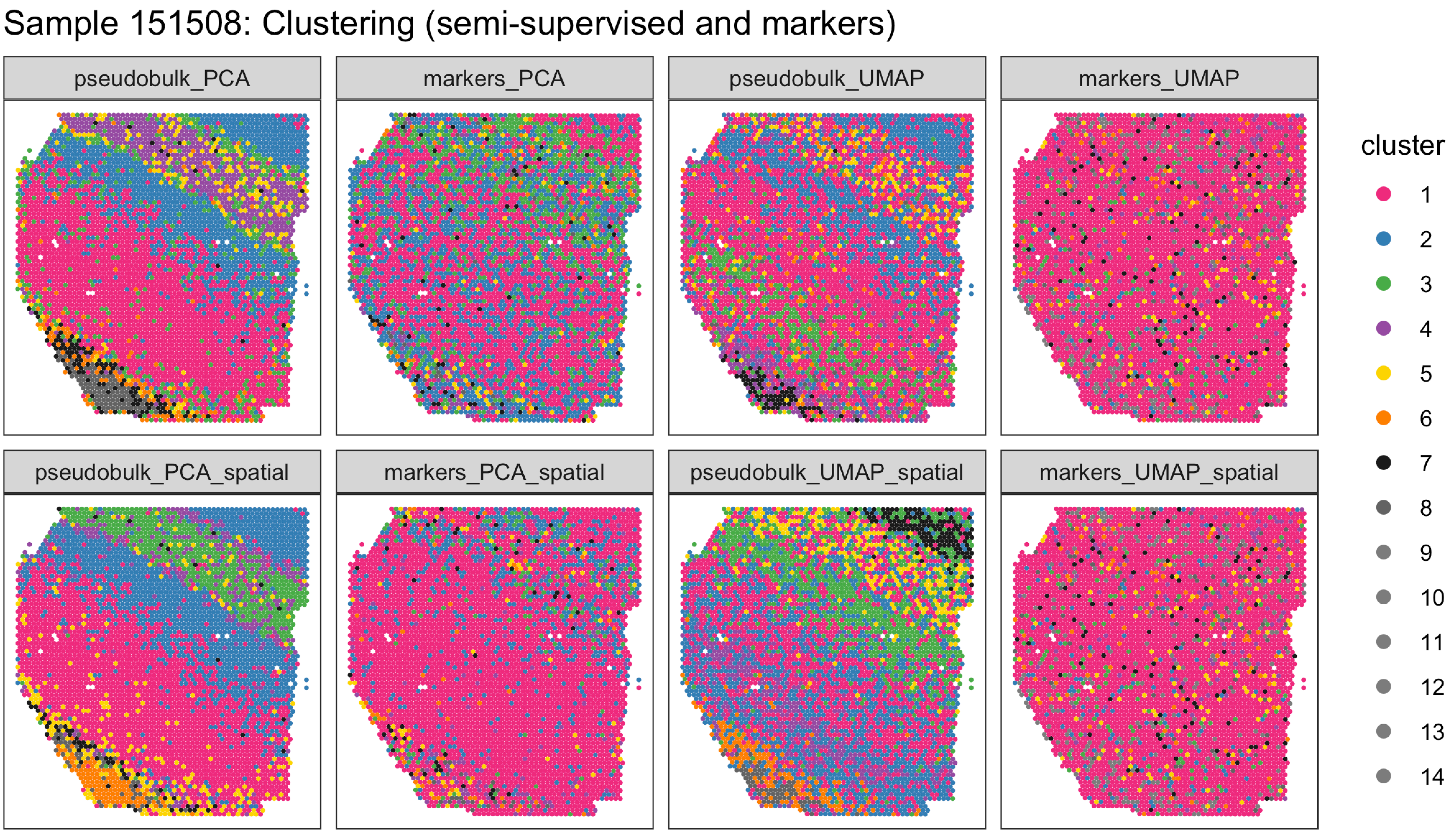

Sample 151508: Clustering (semi-supervised and markers)
pseudobulk_PCA
markers_PCA
pseudobulk_UMAP
markers_UMAP
pseudobulk_PCA_spatial
markers_PCA_spatial
pseudobulk_UMAP_spatial
markers_UMAP_spatial
cluster
1
2
3
4
5
6
7
8
9
10
11
12
13
14

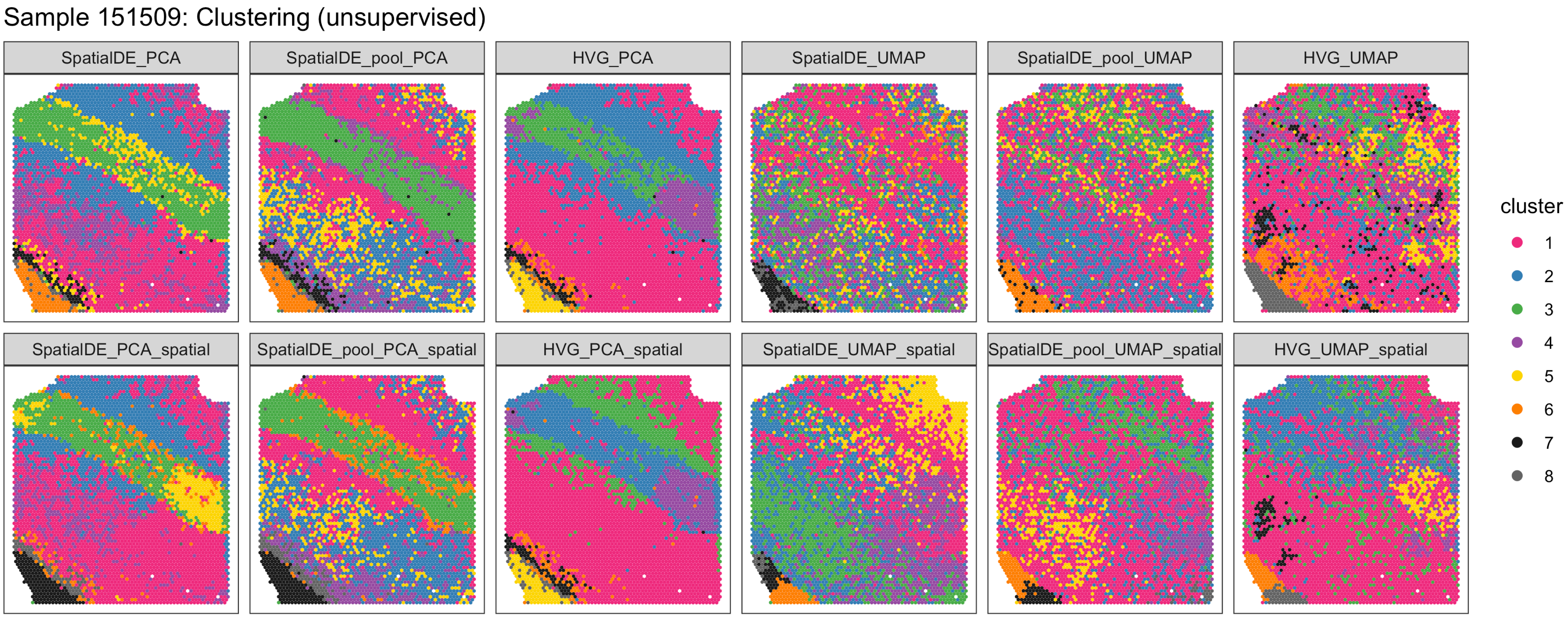

Sample 151509: Clustering (unsupervised)
SpatialDE_PCA
SpatialDE_pool_PCA
HVG_PCA
SpatialDE_UMAP
SpatialDE_pool_UMAP
HVG_UMAP
SpatialDE_PCA_spatial
SpatialDE_pool_PCA_spatial
HVG_PCA_spatial
SpatialDE_UMAP_spatial
SpatialDE_pool_UMAP_spatial
HVG_UMAP_spatial
cluster
1
2
3
4
5
6
7
8

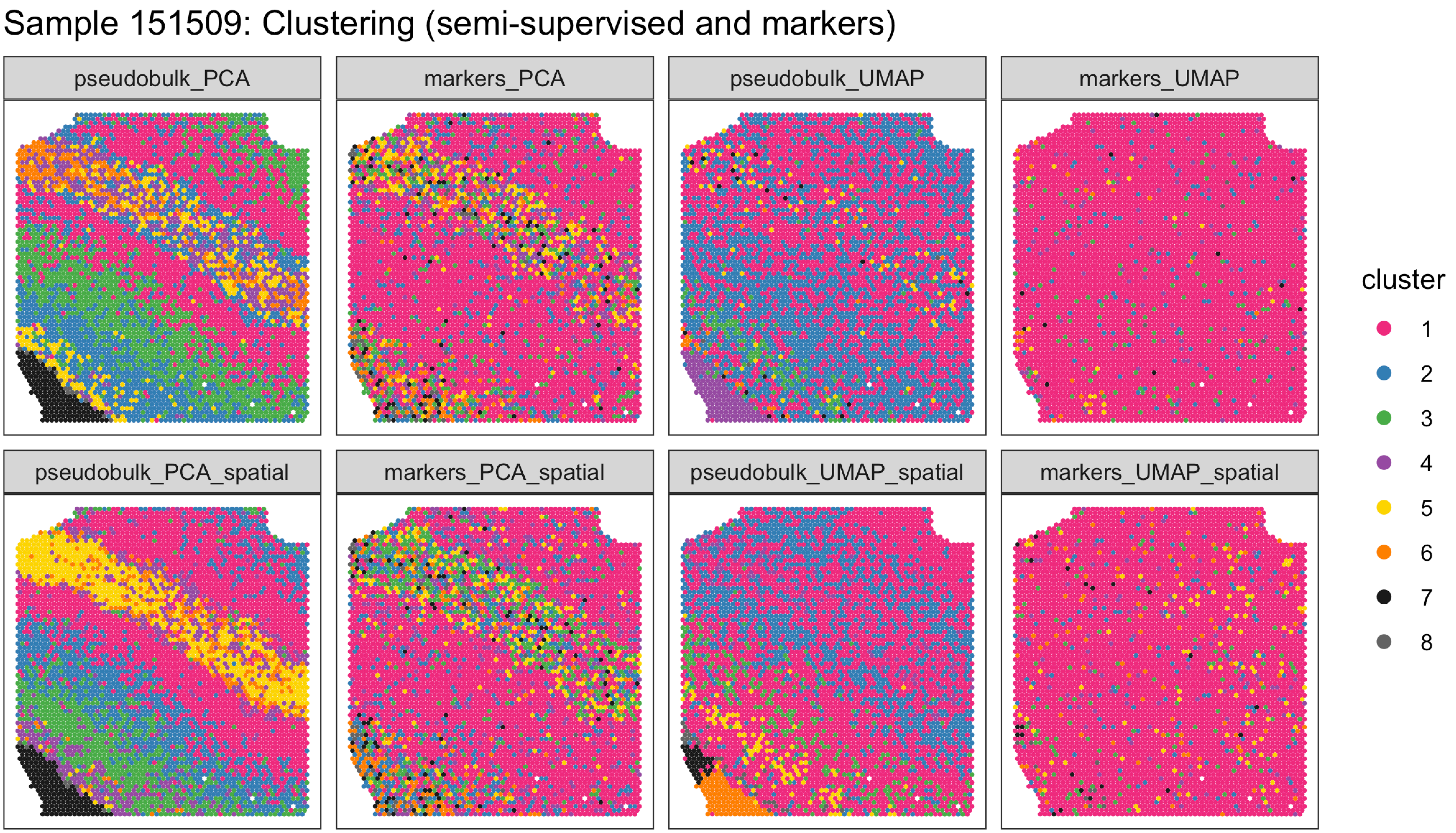

Sample 151509: Clustering (semi-supervised and markers)
pseudobulk_PCA
markers_PCA
pseudobulk_UMAP
markers_UMAP
pseudobulk_PCA_spatial
markers_PCA_spatial
pseudobulk_UMAP_spatial
markers_UMAP_spatial
cluster
1
2
3
4
5
6
7
8

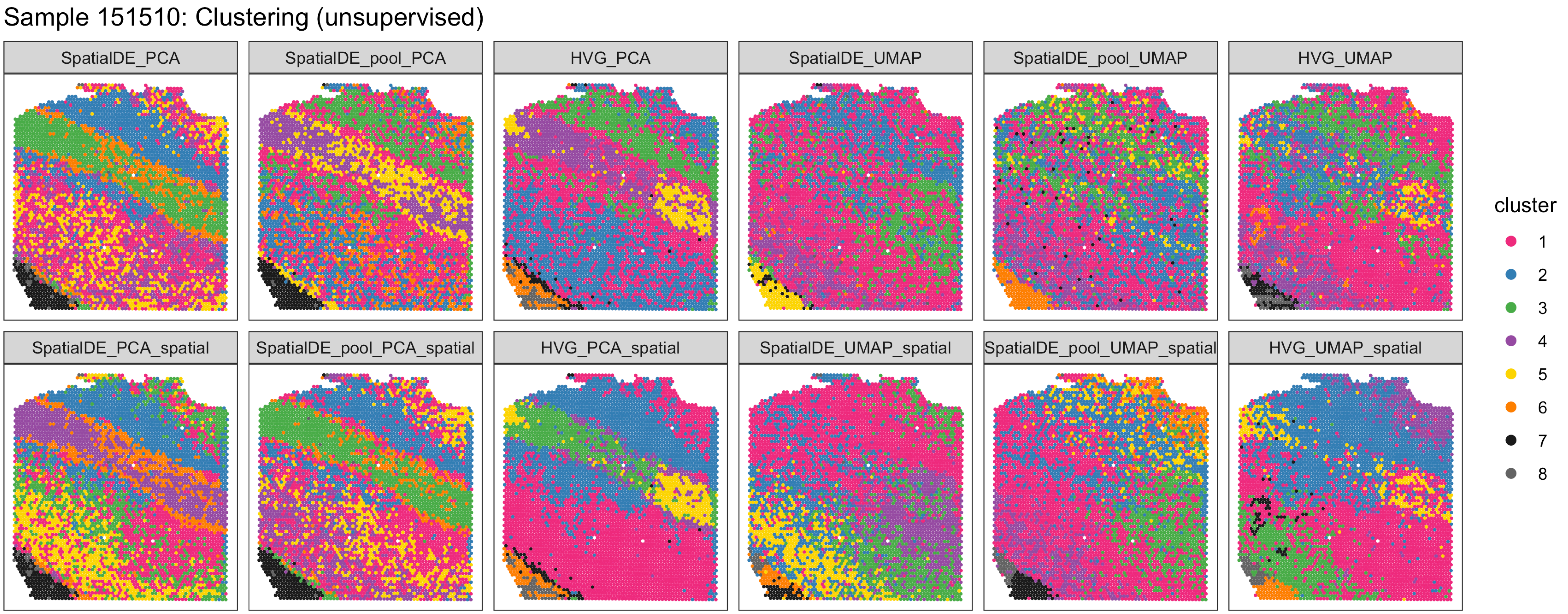
Sample 151510: Clustering (unsupervised)
SpatialDE_PCA
SpatialDE_pool_PCA
HVG_PCA
SpatialDE_UMAP
SpatialDE_pool_UMAP
HVG_UMAP
SpatialDE_PCA_spatial
SpatialDE_pool_PCA_spatial
HVG_PCA_spatial
SpatialDE_UMAP_spatial
SpatialDE_pool_UMAP_spatial
HVG_UMAP_spatial
cluster
1
2
3
4
5
6
7
8

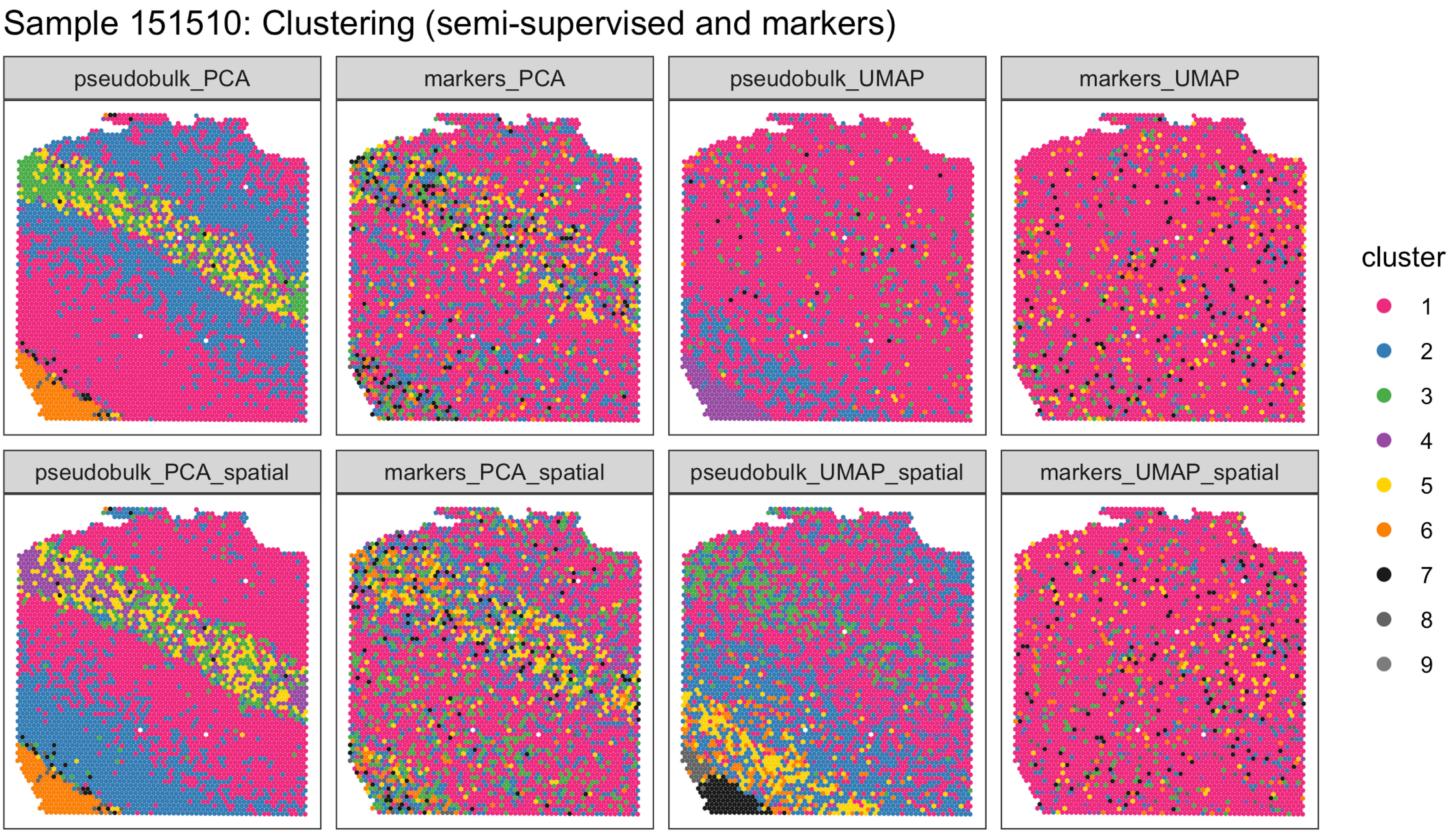
Sample 151510: Clustering (semi-supervised and markers)
pseudobulk_PCA
markers_PCA
pseudobulk_UMAP
markers_UMAP
pseudobulk_PCA_spatial
markers_PCA_spatial
pseudobulk_UMAP_spatial
markers_UMAP_spatial
cluster
1
2
3
4
5
6
7
8
9

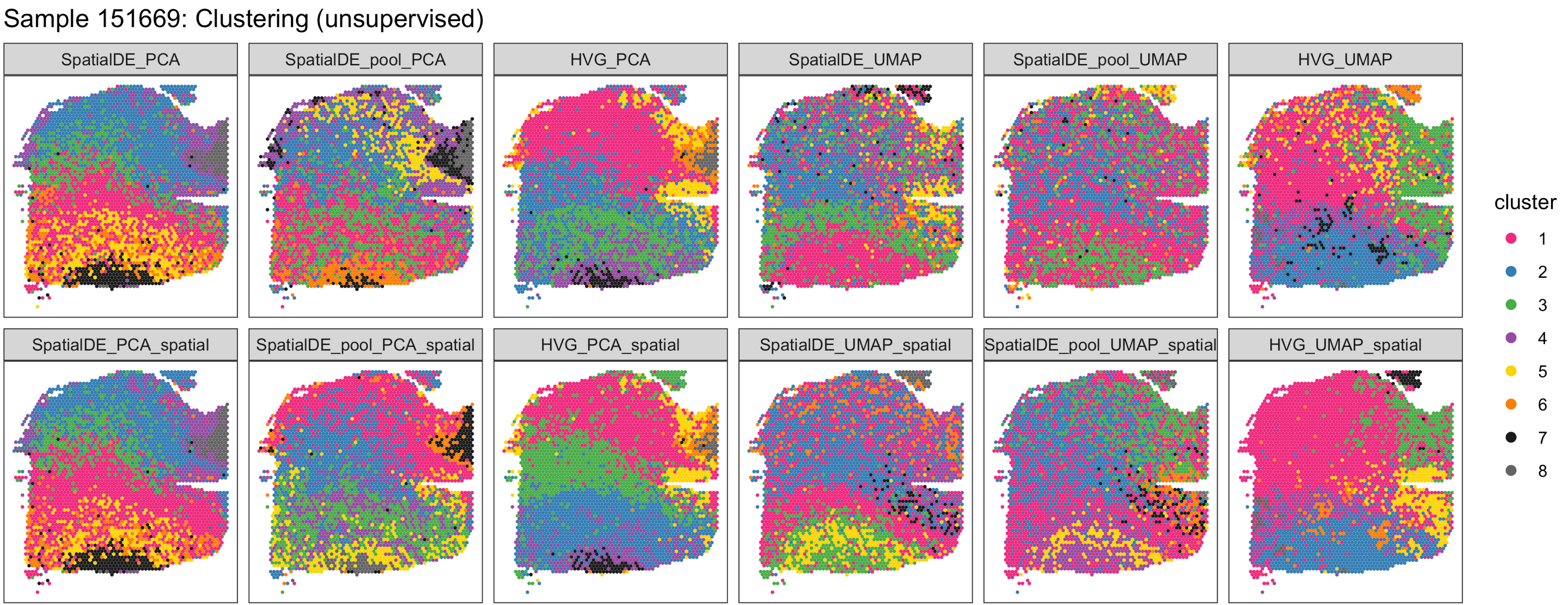
Sample 151669: Clustering (unsupervised)
SpatialDE_PCA
SpatialDE_pool_PCA
HVG_PCA
SpatialDE_UMAP
SpatialDE_pool_UMAP
HVG_UMAP
SpatialDE_PCA_spatial
SpatialDE_pool_PCA_spatial
HVG_PCA_spatial
SpatialDE_UMAP_spatial
SpatialDE_pool_UMAP_spatial
HVG_UMAP_spatial
cluster
1
2
3
4
5
6
7
8

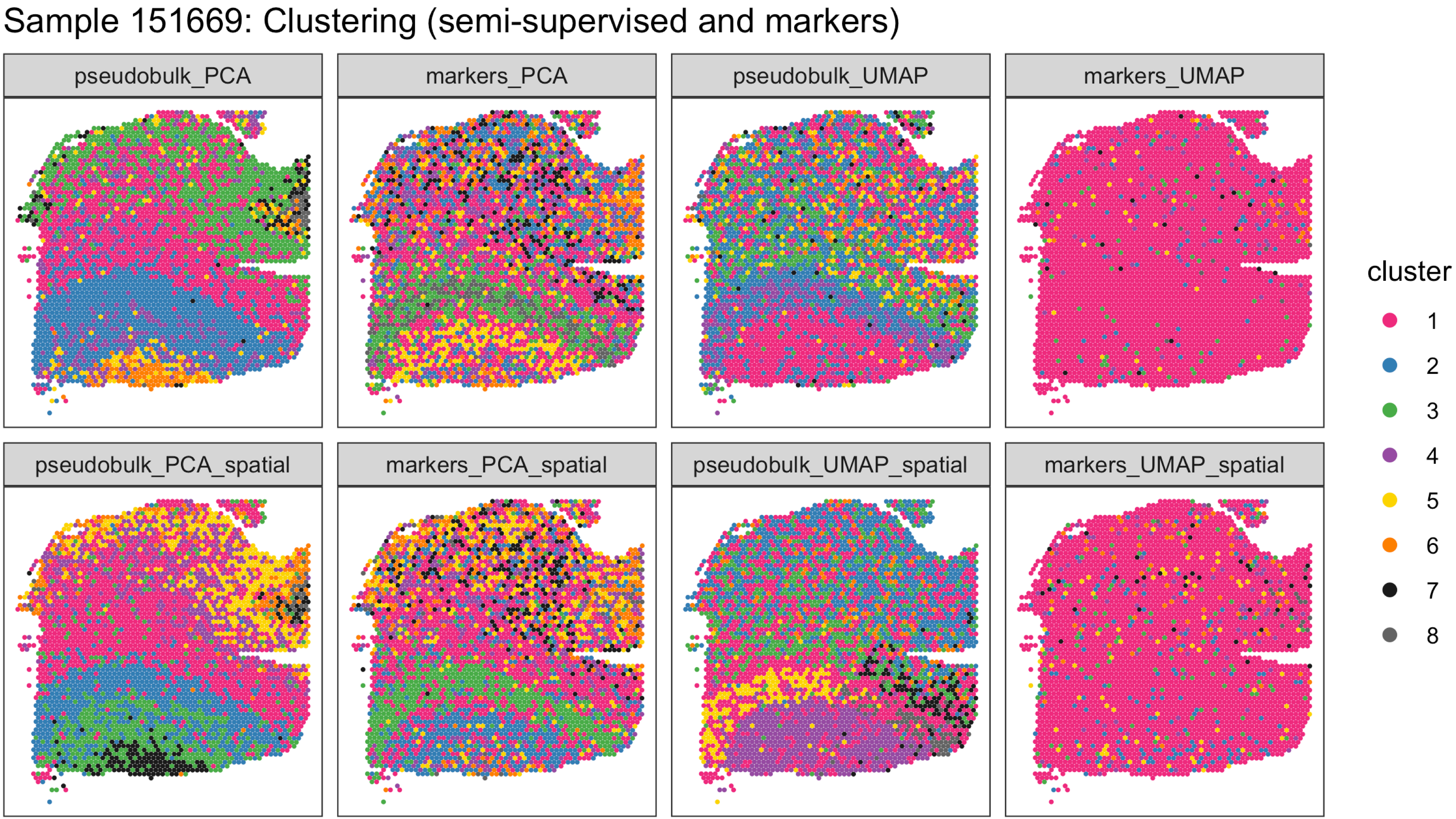
Sample 151669: Clustering (semi-supervised and markers)
pseudobulk_PCA
markers_PCA
pseudobulk_UMAP
markers_UMAP
pseudobulk_PCA_spatial
markers_PCA_spatial
pseudobulk_UMAP_spatial
markers_UMAP_spatial
cluster
1
2
3
4
5
6
7
8

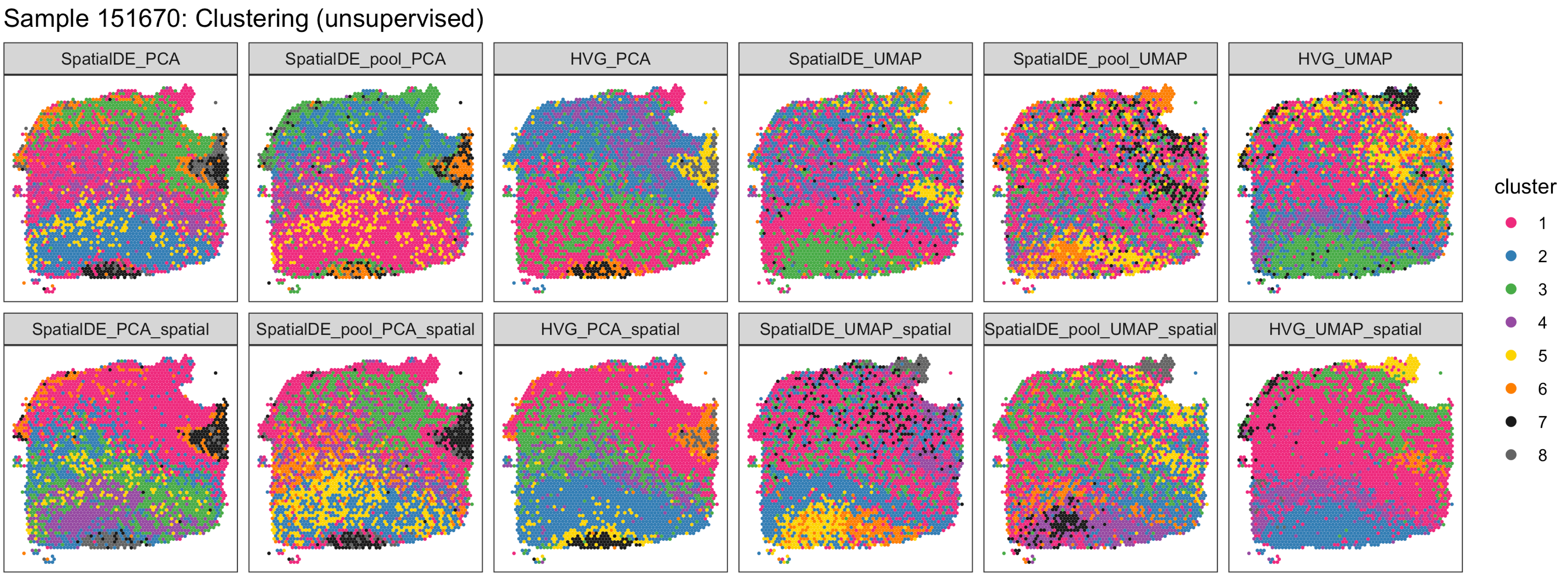
Sample 151670: Clustering (unsupervised)
SpatialDE_PCA
SpatialDE_pool_PCA
HVG_PCA
SpatialDE_UMAP
SpatialDE_pool_UMAP
HVG_UMAP
SpatialDE_PCA_spatial
SpatialDE_pool_PCA_spatial
HVG_PCA_spatial
SpatialDE_UMAP_spatial
SpatialDE_pool_UMAP_spatial
HVG_UMAP_spatial
cluster
1
2
3
4
5
6
7
8

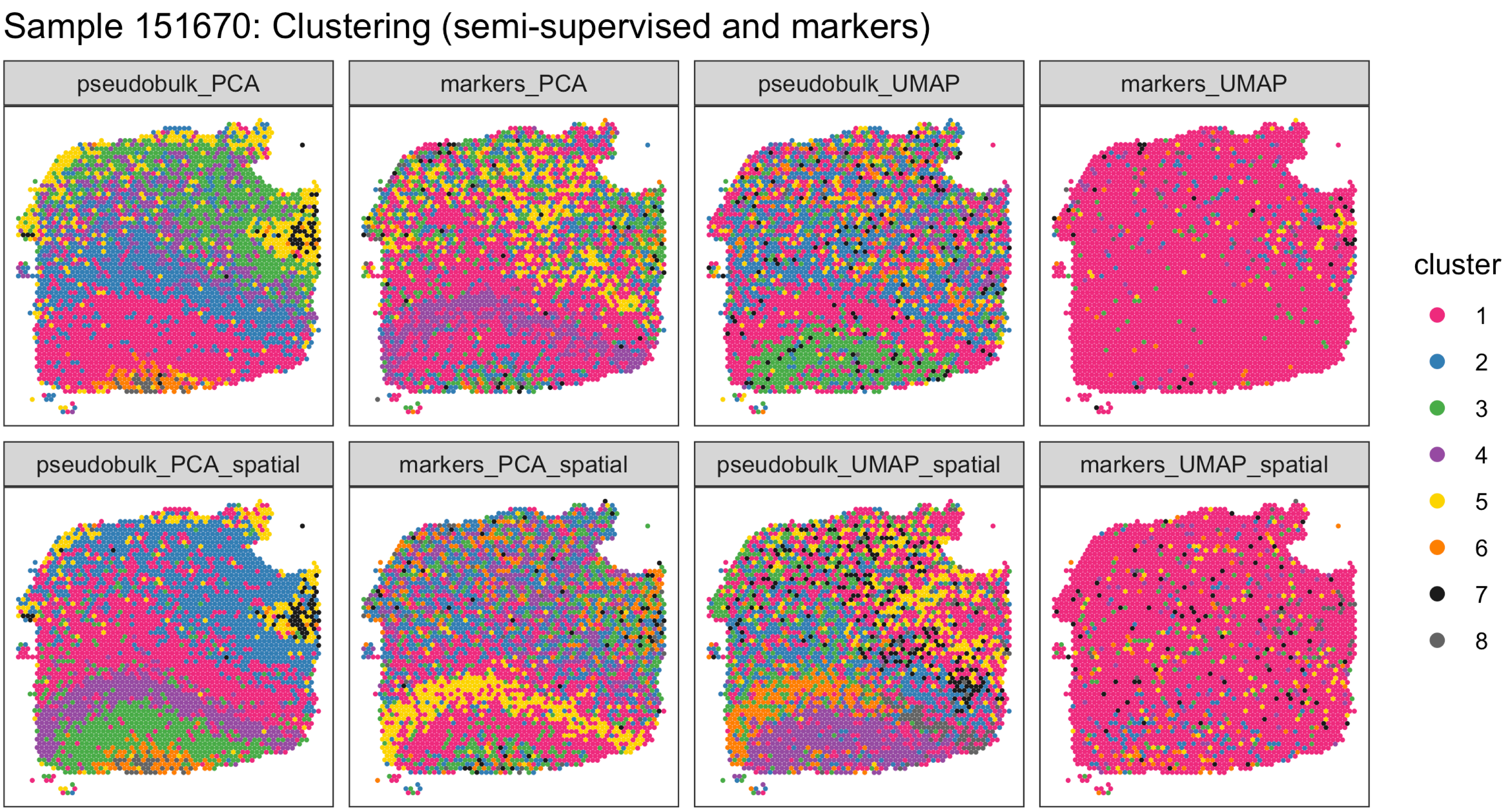

Sample 151670: Clustering (semi-supervised and markers)
pseudobulk_PCA
markers_PCA
pseudobulk_UMAP
markers_UMAP
pseudobulk_PCA_spatial
markers_PCA_spatial
pseudobulk_UMAP_spatial
markers_UMAP_spatial
cluster
1
2
3
4
5
6
7
8

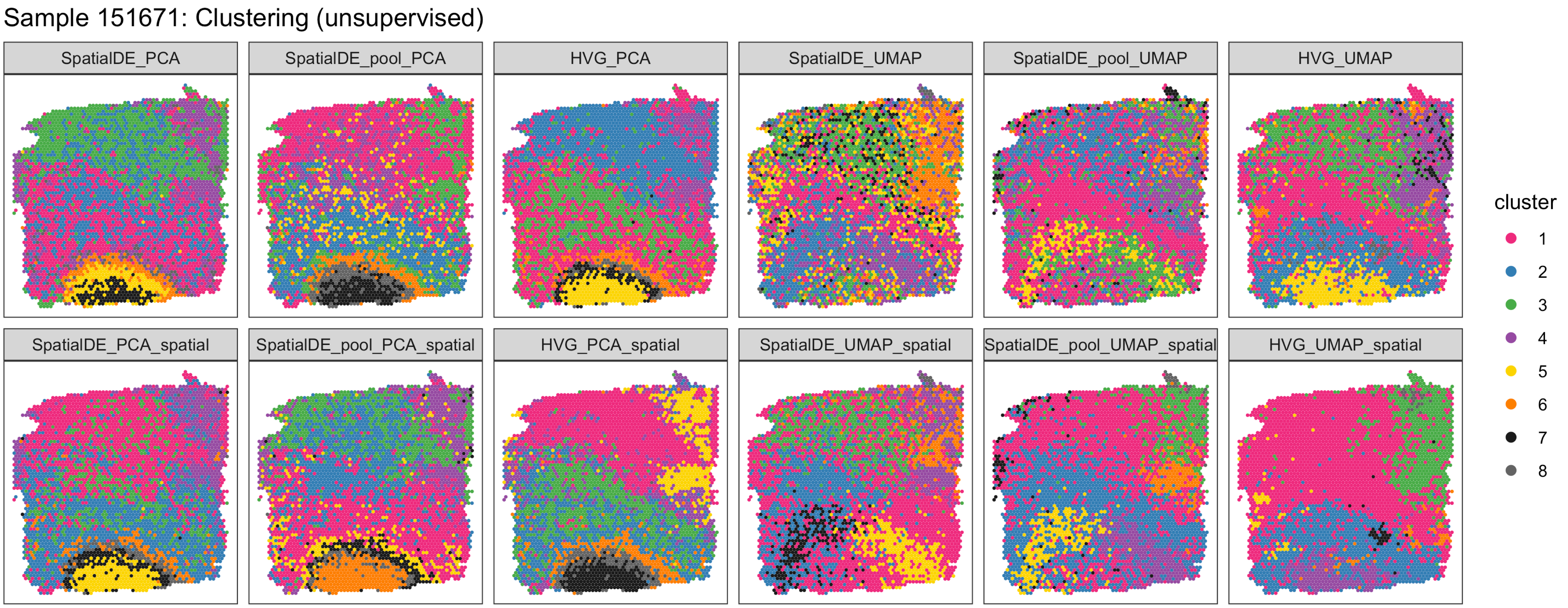
Sample 151671: Clustering (unsupervised)
SpatialDE_PCA
SpatialDE_pool_PCA
HVG_PCA
SpatialDE_UMAP
SpatialDE_pool_UMAP
HVG_UMAP
SpatialDE_PCA_spatial
SpatialDE_pool_PCA_spatial
HVG_PCA_spatial
SpatialDE_UMAP_spatial
SpatialDE_pool_UMAP_spatial
HVG_UMAP_spatial
cluster
1
2
3
4
5
6
7
8

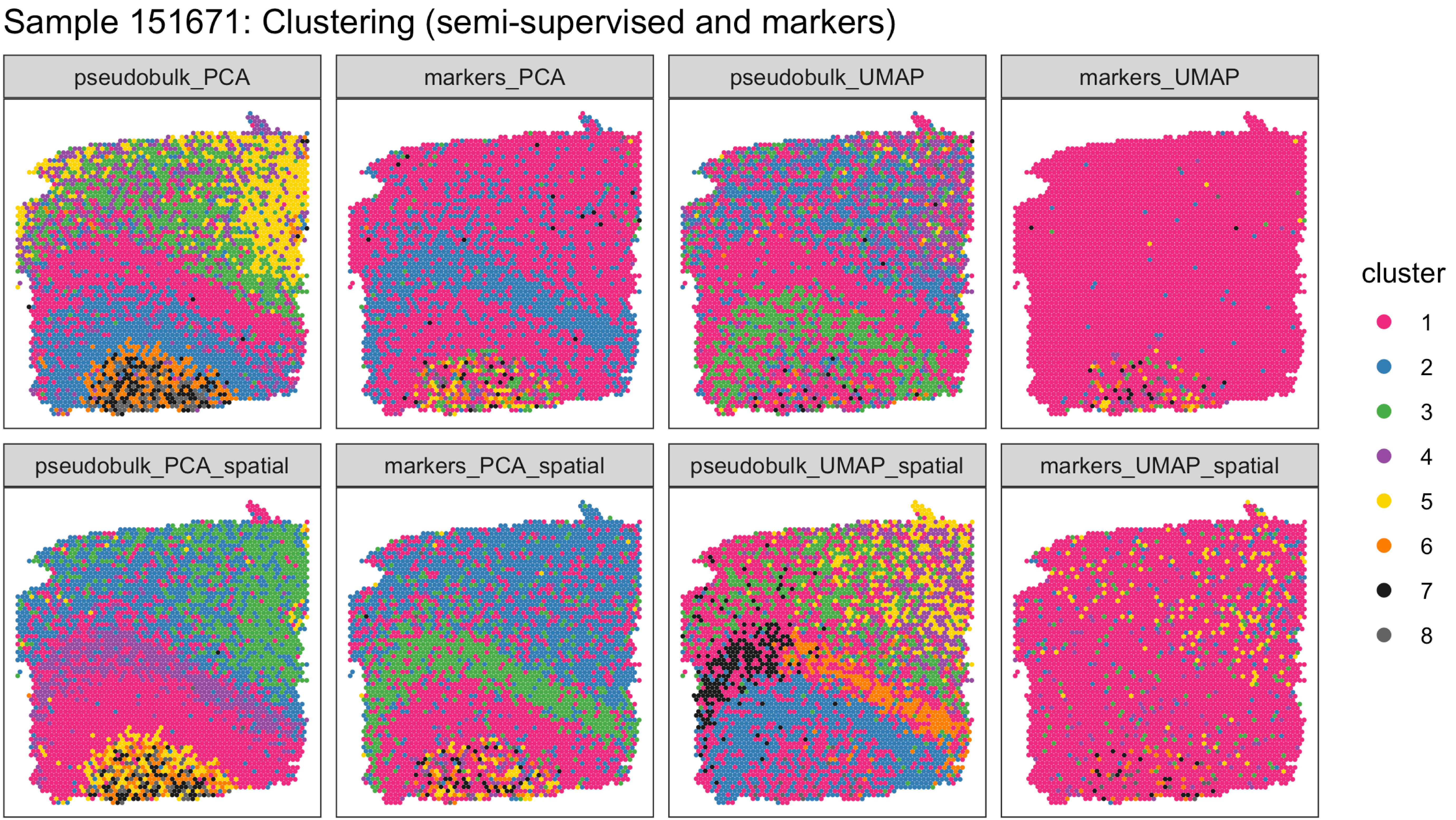

Sample 151671: Clustering (semi-supervised and markers)
pseudobulk_PCA
markers_PCA
pseudobulk_UMAP
markers_UMAP
pseudobulk_PCA_spatial
markers_PCA_spatial
pseudobulk_UMAP_spatial
markers_UMAP_spatial
cluster
1
2
3
4
5
6
7
8

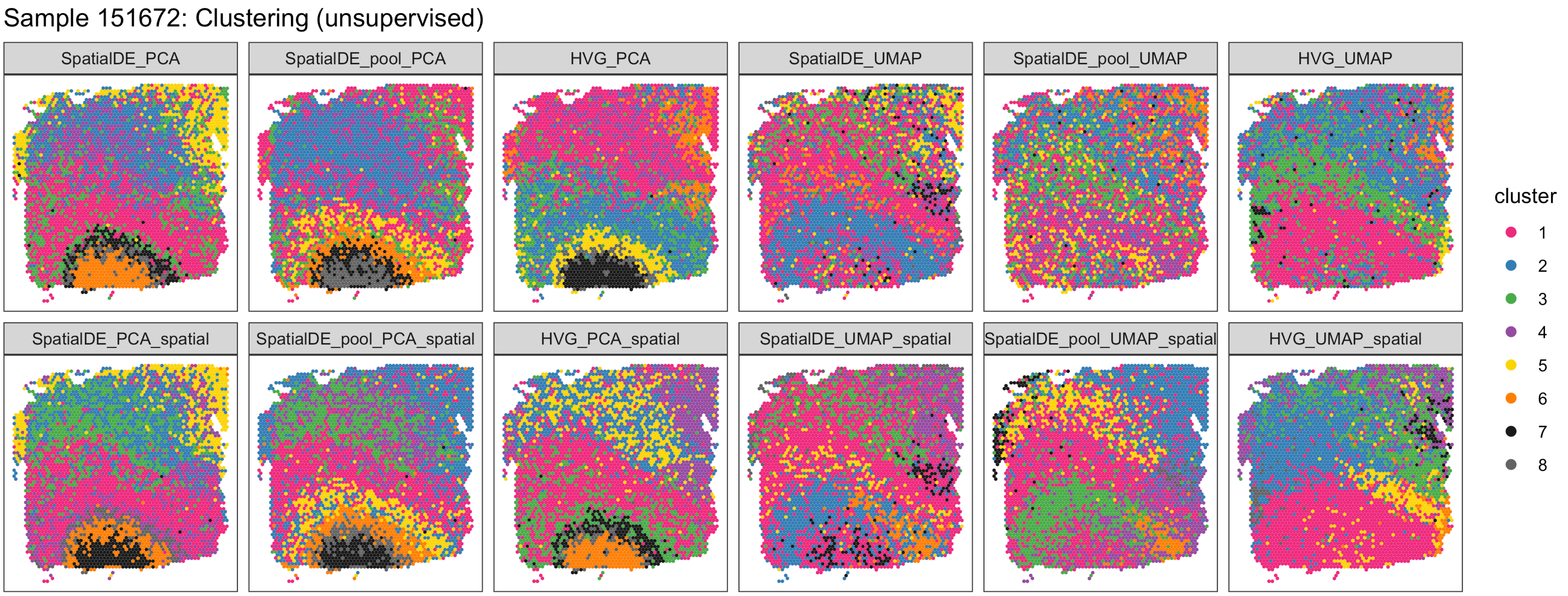

Sample 151672: Clustering (unsupervised)
SpatialDE_PCA
SpatialDE_pool_PCA
HVG_PCA
SpatialDE_UMAP
SpatialDE_pool_UMAP
HVG_UMAP
SpatialDE_PCA_spatial
SpatialDE_pool_PCA_spatial
HVG_PCA_spatial
SpatialDE_UMAP_spatial
SpatialDE_pool_UMAP_spatial
HVG_UMAP_spatial
cluster
1
2
3
4
5
6
7
8

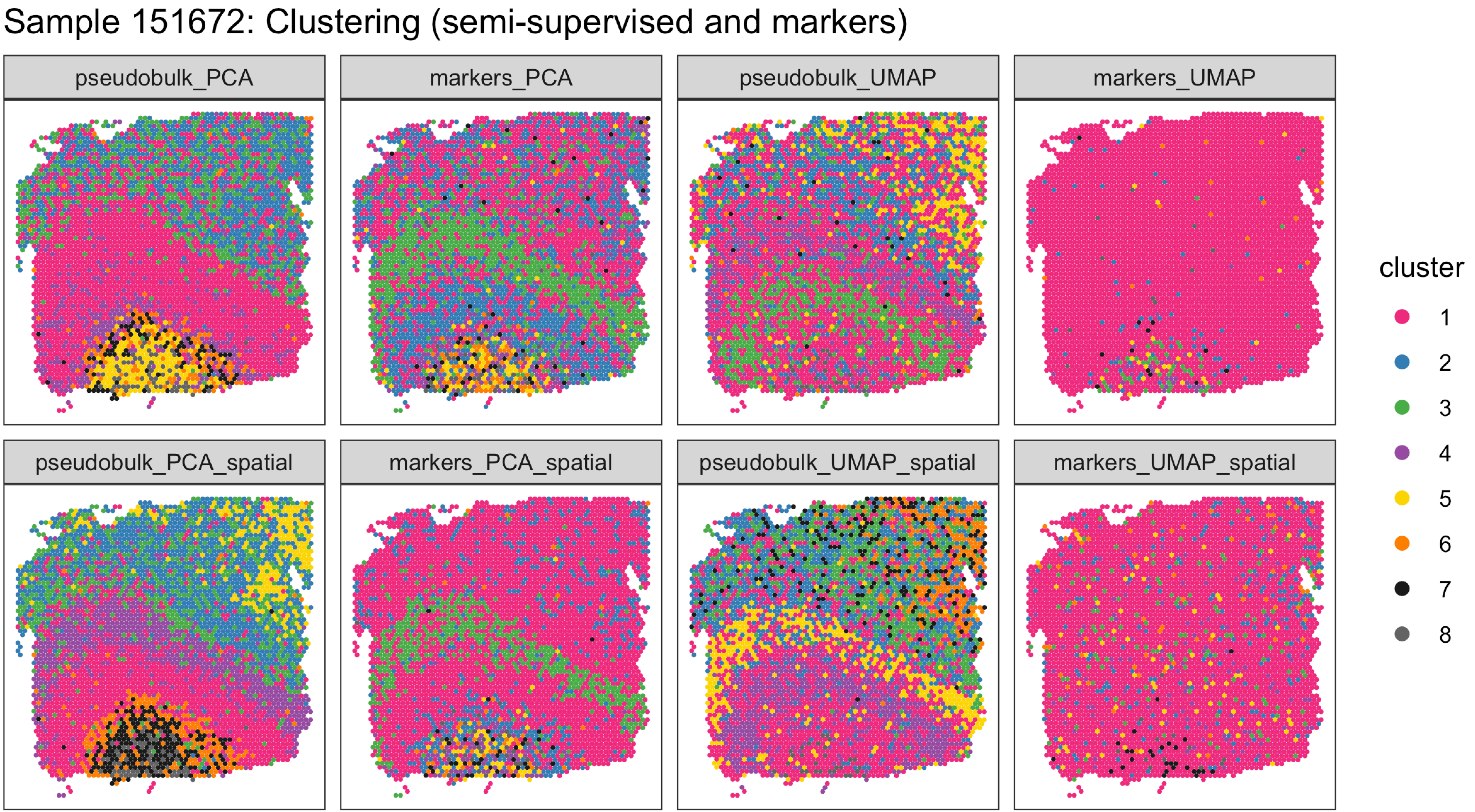

Sample 151672: Clustering (semi-supervised and markers)
pseudobulk_PCA
markers_PCA
pseudobulk_UMAP
markers_UMAP
pseudobulk_PCA_spatial
markers_PCA_spatial
pseudobulk_UMAP_spatial
markers_UMAP_spatial
cluster
1
2
3
4
5
6
7
8

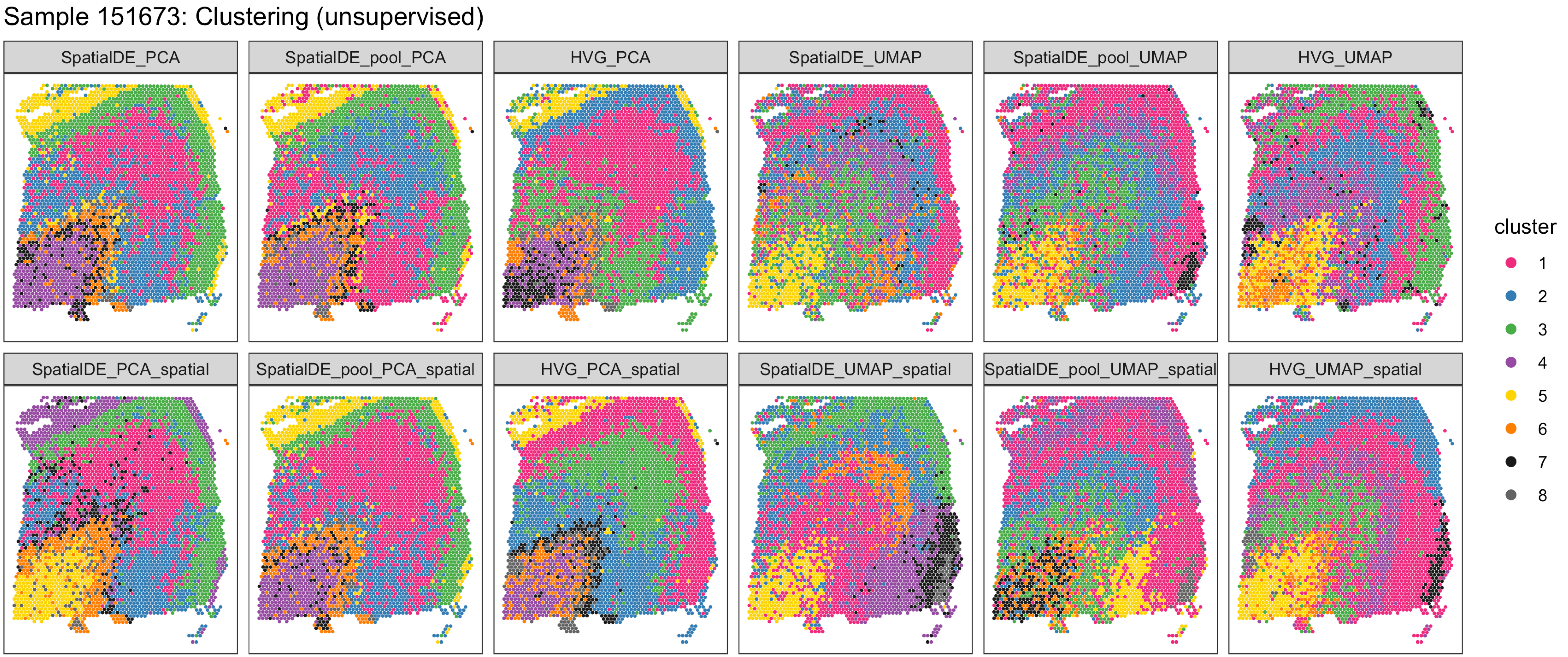
Sample 151673: Clustering (unsupervised)
SpatialDE_PCA
SpatialDE_pool_PCA
HVG_PCA
SpatialDE_UMAP
SpatialDE_pool_UMAP
HVG_UMAP
SpatialDE_PCA_spatial
SpatialDE_pool_PCA_spatial
HVG_PCA_spatial
SpatialDE_UMAP_spatial
SpatialDE_pool_UMAP_spatial
HVG_UMAP_spatial
cluster
1
2
3
4
5
6
7
8

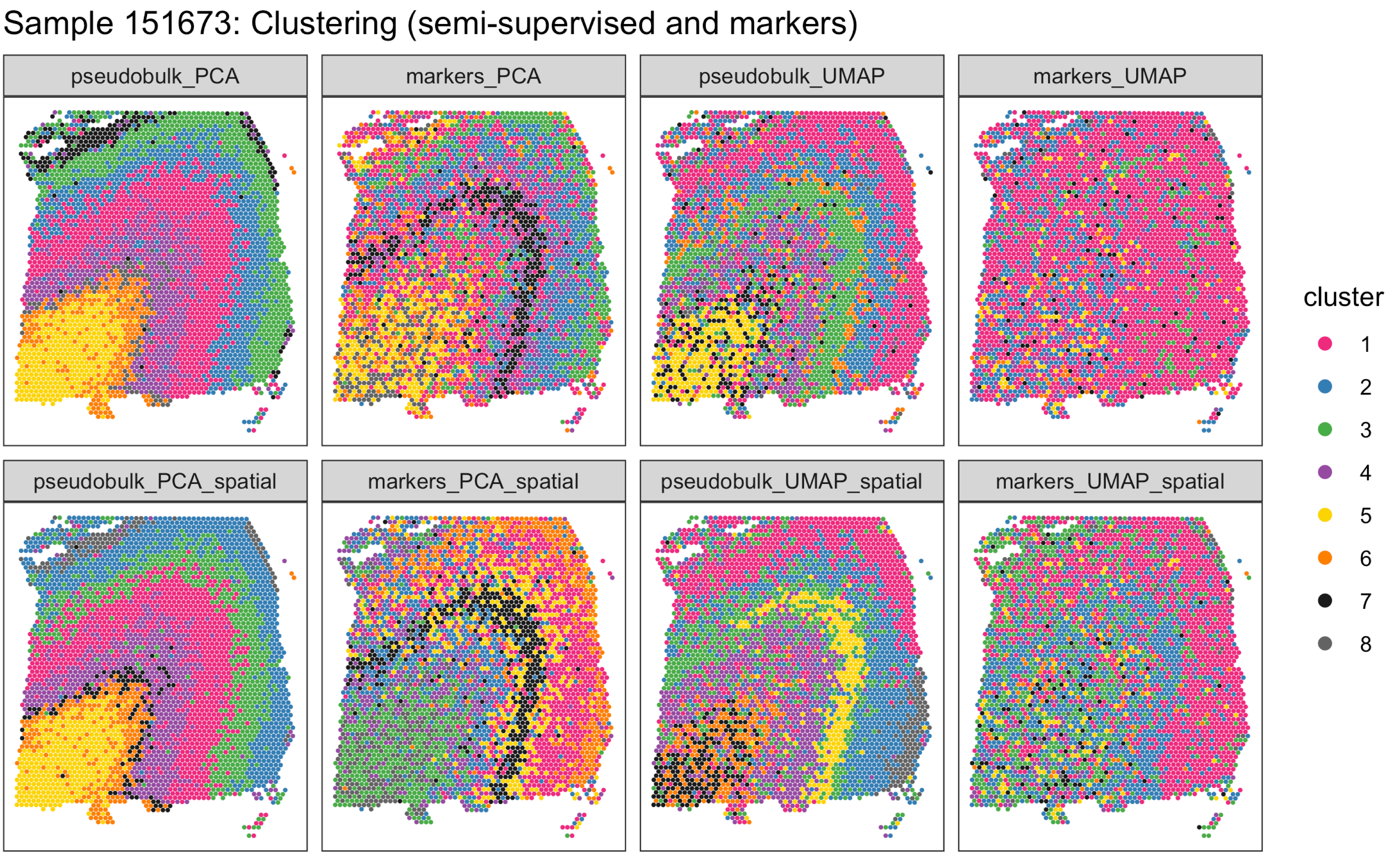
Sample 151673: Clustering (semi-supervised and markers)
pseudobulk_PCA
markers_PCA
pseudobulk_UMAP
markers_UMAP
pseudobulk_PCA_spatial
markers_PCA_spatial
pseudobulk_UMAP_spatial
markers_UMAP_spatial
cluster
1
2
3
4
5
6
7
8

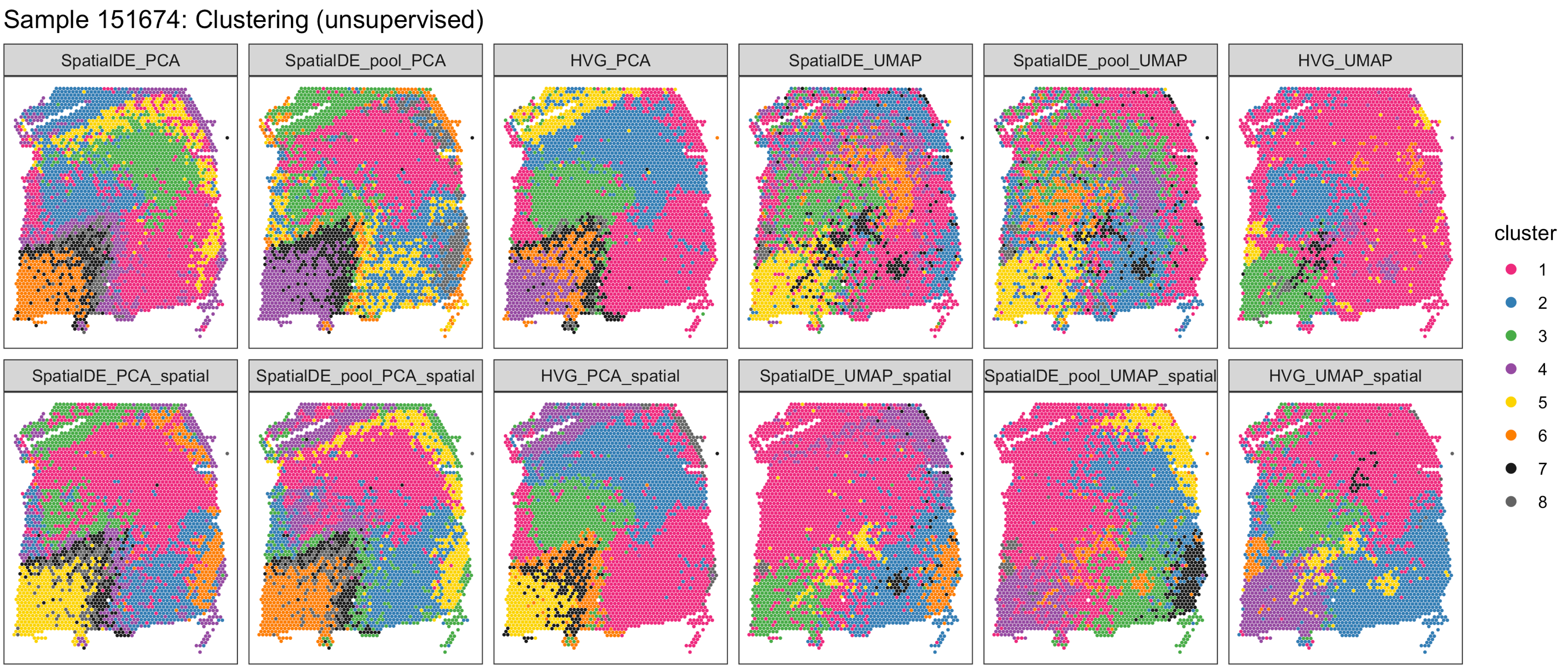
Sample 151674: Clustering (unsupervised)
SpatialDE_PCA
SpatialDE_pool_PCA
HVG_PCA
SpatialDE_UMAP
SpatialDE_pool_UMAP
HVG_UMAP
SpatialDE_PCA_spatial
SpatialDE_pool_PCA_spatial
HVG_PCA_spatial
SpatialDE_UMAP_spatial
SpatialDE_pool_UMAP_spatial
HVG_UMAP_spatial
cluster
1
2
3
4
5
6
7
8

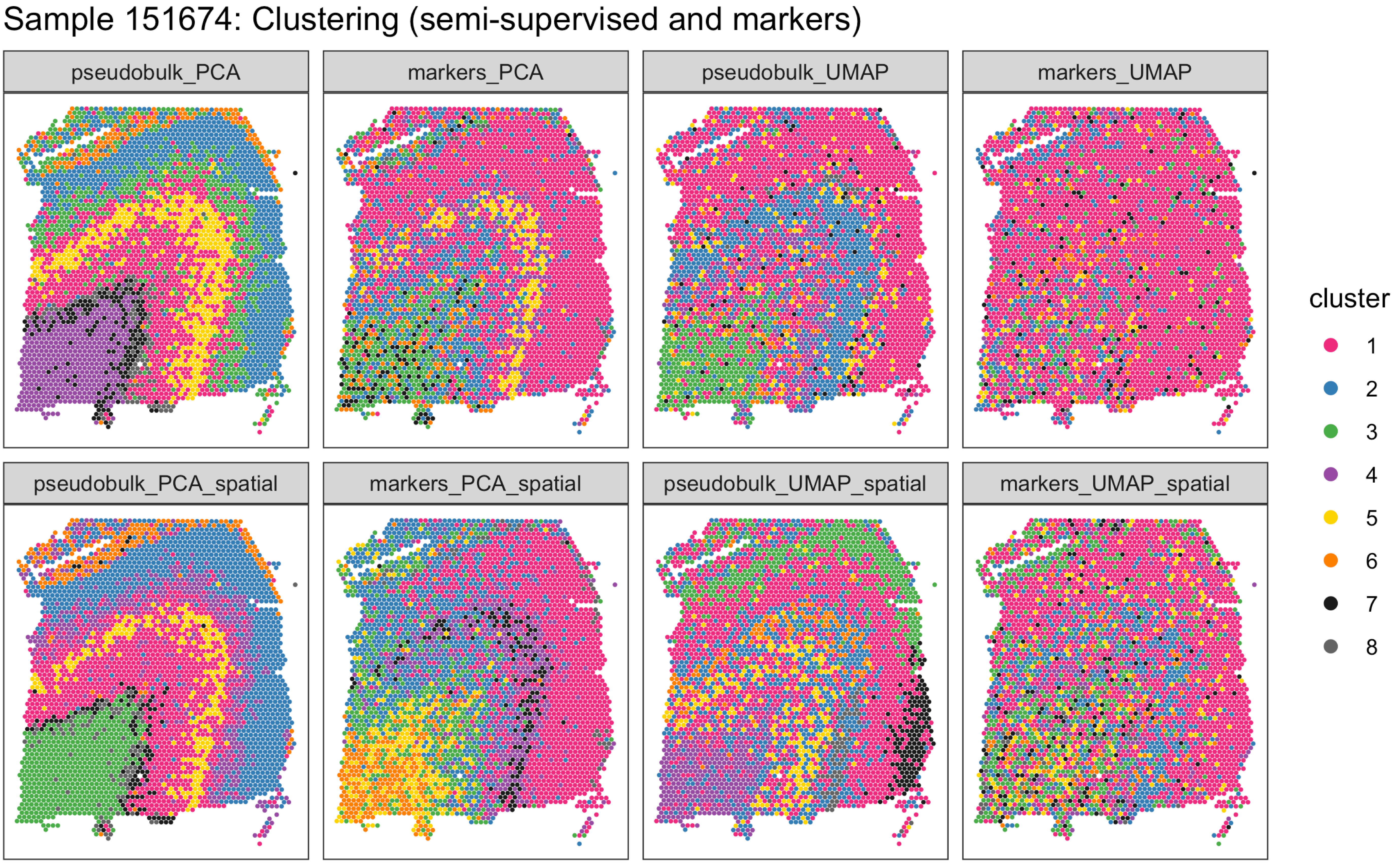

Sample 151674: Clustering (semi-supervised and markers)
pseudobulk_PCA
markers_PCA
pseudobulk_UMAP
markers_UMAP
pseudobulk_PCA_spatial
markers_PCA_spatial
pseudobulk_UMAP_spatial
markers_UMAP_spatial
cluster
1
2
3
4
5
6
7
8

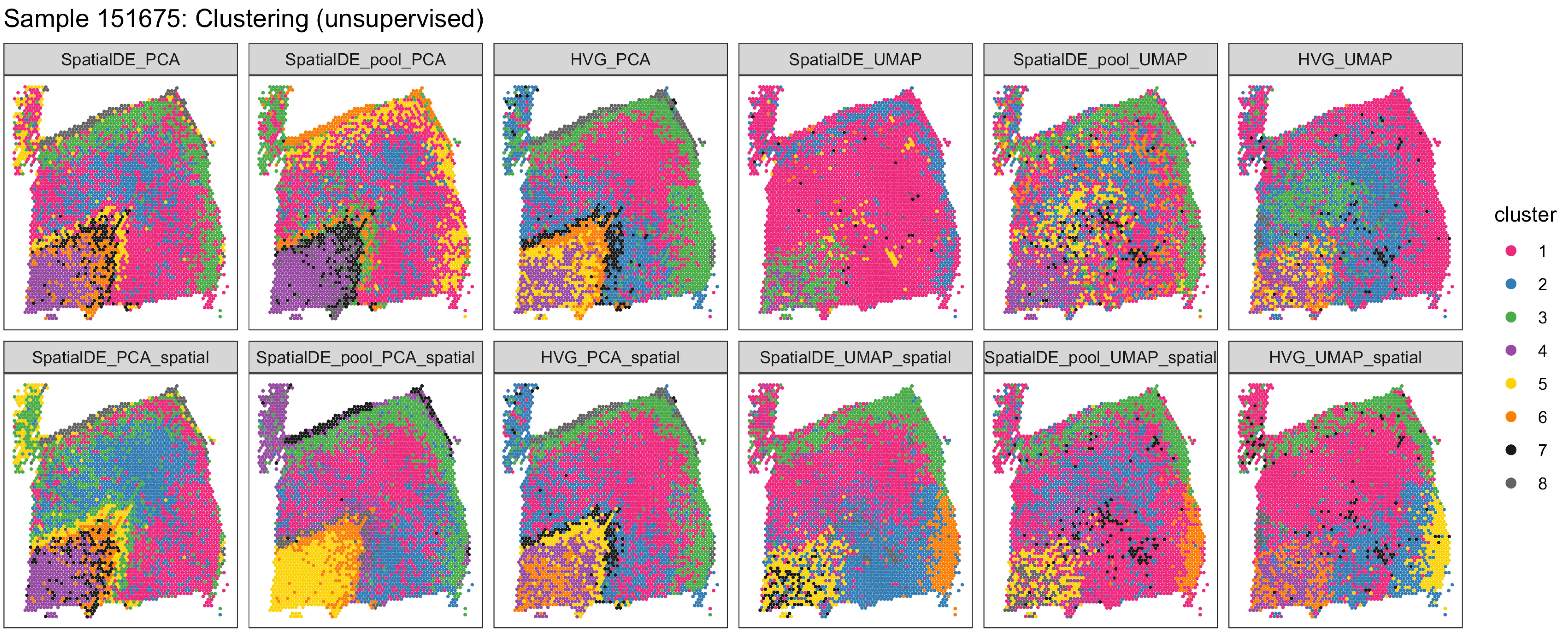

Sample 151675: Clustering (unsupervised)
SpatialDE_PCA
SpatialDE_pool_PCA
HVG_PCA
SpatialDE_UMAP
SpatialDE_pool_UMAP
HVG_UMAP
SpatialDE_PCA_spatial
SpatialDE_pool_PCA_spatial
HVG_PCA_spatial
SpatialDE_UMAP_spatial
SpatialDE_pool_UMAP_spatial
HVG_UMAP_spatial
cluster
1
2
3
4
5
6
7
8

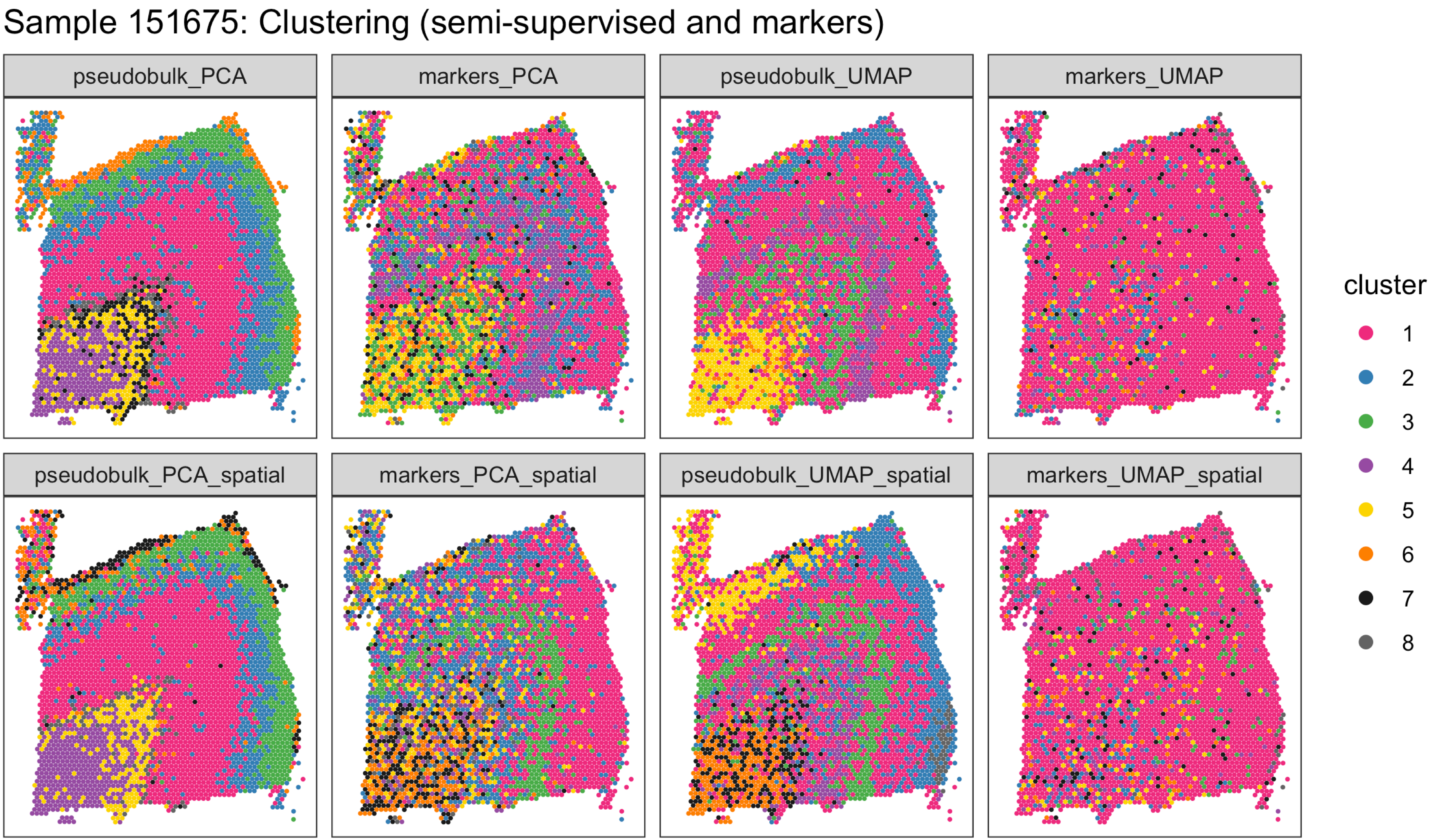

Sample 151675: Clustering (semi-supervised and markers)
pseudobulk_PCA
markers_PCA
pseudobulk_UMAP
markers_UMAP
pseudobulk_PCA_spatial
markers_PCA_spatial
pseudobulk_UMAP_spatial
markers_UMAP_spatial
cluster
1
2
3
4
5
6
7
8

Sample 151676: Clustering (unsupervised)

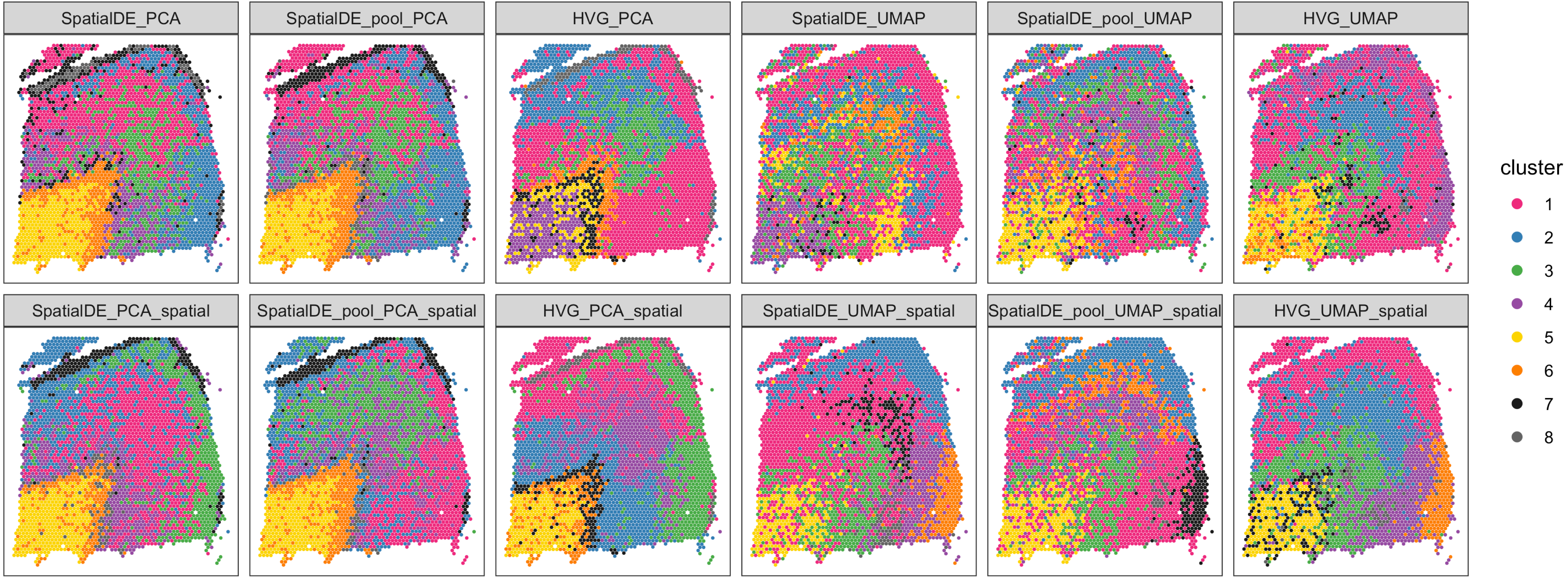

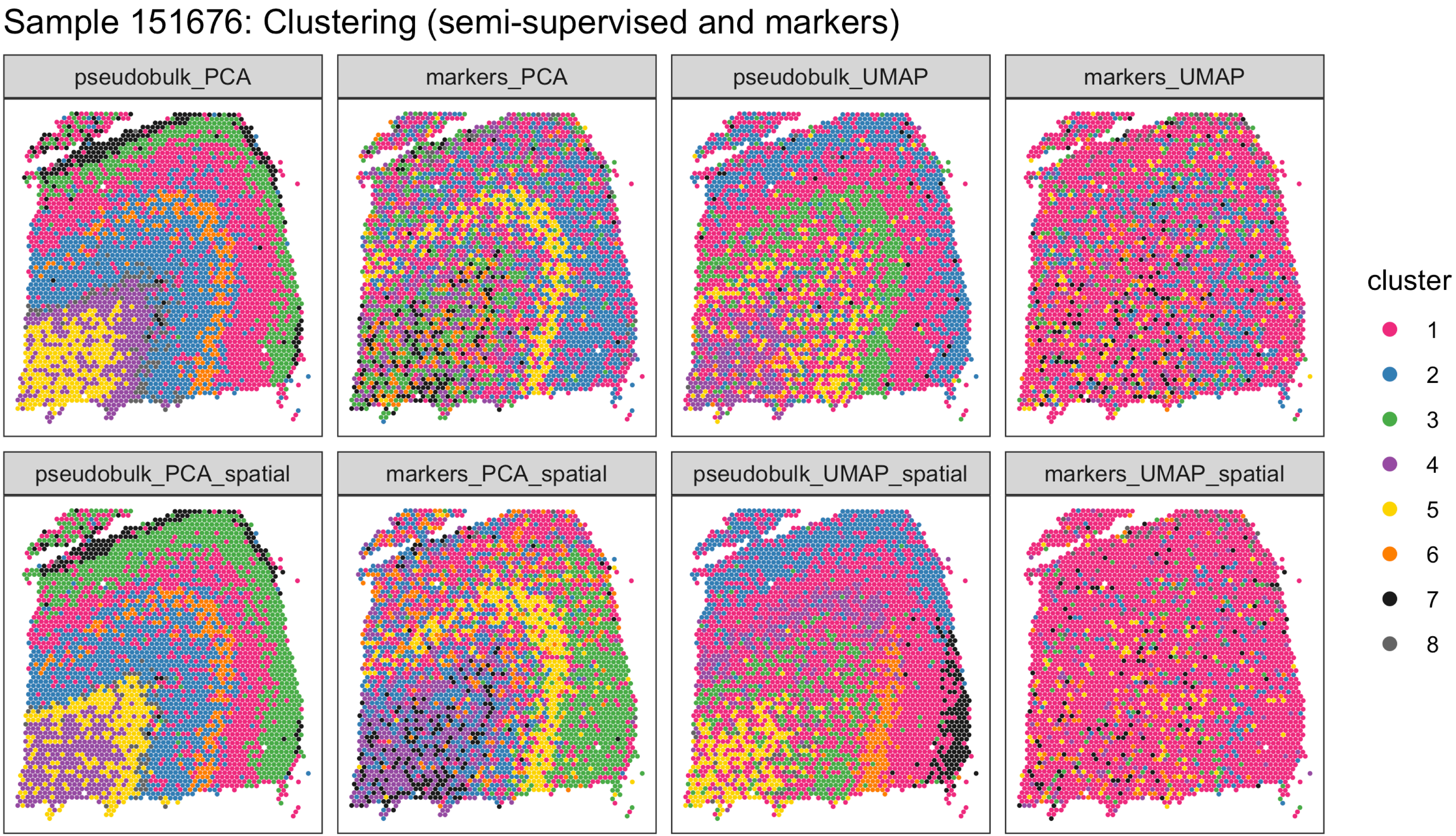
Sample 151676: Clustering (semi-supervised and markers)
pseudobulk_PCA
markers_PCA
pseudobulk_UMAP
markers_UMAP
pseudobulk_PCA_spatial
markers_PCA_spatial
pseudobulk_UMAP_spatial
markers_UMAP_spatial
cluster
1
2
3
4
5
6
7
8
